# Single-base substitutions switch RNA tertiary allostery between positive and negative coupling

**DOI:** 10.64898/2026.08.09.743768

**Authors:** Alabi Oladeji, Brandon Kircher, Joseph D Yesselman

## Abstract

Riboswitches regulate gene expression in response to ligand binding, remodeling their secondary structure to terminate transcription or block translation. This innate switching has driven efforts to repurpose riboswitches as synthetic control elements. Although promising, tuning riboswitch responses to new ligands and functions has proven difficult. Large RNAs such as the ribosome and RNase P exhibit an alternative switching mode via the docking and undocking of tertiary contacts. These transitions are buried deep within the machines, and there is no minimal system to study them in isolation. To overcome this limitation, we built a minimal 3D-structure ligand-inducible switch. We started with a previously designed nanostructure containing an ATP-aptamer and a tetraloop/tetraloop receptor (TL/TLR) tertiary contact. DMS-MaPseq screening of 3,375 mutants of the two linking motifs, a kink-turn and a 4-1 junction, identified 237 variants in which AMP binding undocked the TL/TLR and 120 in which it drove docking. Switching arose almost entirely from mutations in the conserved sheared G·A base pairs of the kink-turn. Fitting four representative constructs’ AMP titrations to a linked-equilibrium model yielded coupling free energies spanning 3.7 kcal/mol, and magnesium titrations with and without AMP independently confirmed these couplings while revealing a three-state docking pathway in mutants lacking wild-type coupling. Single nucleotides inserted into the kink-turn motif adjust the sign and strength of coupling without redesigning the secondary structure. This simple system thus provides a platform both as a model for tertiary switching in other large RNA machines and as a foundation for designing synthetic control elements based on tertiary interactions.

## Introduction

Structured RNAs perform essential biological functions, including peptide bond formation, tRNA maturation, and telomere extension (1–4). To perform these functions, the RNA must fold into a sophisticated three-dimensional (3D) structure (5–8). Many structured RNAs undergo dramatic structural changes upon stimulation (9). Riboswitches are prime examples of this process: binding of a ligand to an aptamer that detects a metabolite prompts a rearrangement of the downstream expression platform, resulting in termination of transcription or repression of translation (10, 11). Riboswitches switch function, with few exceptions, by adopting mutually exclusive base-pairing configurations (12, 13).

Naturally derived riboswitches have been engineered to bind new ligands and regulate new functional outputs, ranging from ribozymes to CRISPR effectors (14–16). However, tuning these switches has proven difficult. Switches that operate by changing base pairing often have slow kinetics (13, 17, 18), tend to misfold (19–21), and have limited dynamic range (22–24). Their directionality is also hard to invert. If a riboswitch terminates transcription upon ligand binding, engineering the opposite response typically requires redesigning the expression platform in concert with the aptamer, which can be difficult.

These limitations likely arise from the switching mechanism itself. Disrupting and reforming a series of base pairs is a slow, kinetic-trapping process and is highly sensitive to sequence changes. While significant progress has been made in developing simpler and more generalizable base-pairing switches (25–30), these designs still rely on base-pair stability, so the underlying limitations persist. The largest RNA machines are less constrained to base-pairing rearrangements. Although they also remodel base pairing, many of their functional transitions are driven by the docking and undocking of long-range tertiary contacts (4, 31–34), and the free energies of these contacts are comparable to those of ligand binding, placing the two within tuning range of one another. These motions have been studied intensively, but they have not been distilled into principles applicable to new synthetic RNAs. A minimal RNA whose state is toggled solely by tertiary rearrangement, driven by small-molecule binding, would allow us to study this mode of switching in isolation. Such a system would provide both a model for the principles of 3D allostery and an alternative engineering strategy in which sensing and output are energetically matched and structurally separable.

Here we report a 3D structure-switching platform controlled by ligand binding. We began with an engineered nanostructure comprised of an ATP aptamer (35, 36) and a peripheral GAAA tetraloop/tetraloop receptor motif (TL/TLR) (37–39). Thousands of mutations were placed between the TL/TLR and ATP aptamer in two structural motifs (kink-turn and 4-1 junction) to yield switching variants. Single-base mutations within the kink-turn yielded variants with a variety of allosteric outcomes: AMP triggered undocking of the contact in some variants and stabilization of the contact in other variants. Representative variants were titrated with AMP and fitted to a four-state linked-equilibrium model. The fits provide a direct measurement of how ligand binding affects tertiary docking, with resulting positive, negative, or zero coupling. Magnesium titrations confirmed the sign of coupling and revealed complex coupling between kink-turn mutants and TL/TLR docking. This system provides a single secondary structure control for tertiary allostery and a framework for studying RNA cooperativity, potentially applicable to other aptamers and tertiary contacts. This system is the first demonstrated minimal model system of tertiary allostery that does not involve a switch in secondary structure. This work lays the groundwork for exploring RNA 3D thermodynamic cooperativity, with potential expansion to other aptamers and tertiary interactions.

## Methods

### Designing a library of RNA mutants

We began with a previously characterized nanostructure with an ATP aptamer and TL/TLR tertiary contact (PDB: 6WLJ) (39). The construct includes five helices and two junctions, a kink-turn and a 4-1 junction that link the ATP aptamer to the TL/TLR contact. For helical-only variants, we randomized all helices excluding flanking pairs for each junction and the two GC pairs that are part of the tetraloop receptor. For junction variants, we generated single and double mutations in the kink-turn, the 4-1 junction, or both. After these junction mutations, we also performed a helix randomization as described above. We finally added an eight-base-pair barcode unique to each sequence at the beginning of the first helix. This also helped to increase sequence diversity. We added a common 5′ (GGAACAGCACUUCGGUGCAAAC) and 3′ (AAAGAAACAACAACAACAAC) sequence for PCR amplification, and finally a T7 promoter (TTCTAATACGACTCACTATA) at the 5′ end. We folded every candidate with ViennaRNA (40) and retained only those sequences with an ensemble defect <5, ensuring that all variants maintained the wild-type fold. The resulting 7,149 sequences were synthesized as an Agilent oligo pool (all sequences available in **Table S1**).

### Generating double-stranded DNA of the RNA variant library

The library oligonucleotide pool came dry and was solvated in 50 μL of 1x IDTE buffer at pH 8.0 (IDT #11-05-01-13). It was then amplified by PCR to generate double-stranded DNA templates for transcription. The 50 μL PCR reaction mixture comprised 25 μL of NEBNext High-Fidelity 2X PCR Master Mix (NEB #M0541S), 18 μL of RNase-free UltraPure water (ThermoFisher #10977015), 2 μL of oligonucleotide pool, and 2.5 μL of each of the forward primer (TTCTAATACGACTCACTATAGG) and reverse primer (GTTGTTGTTGTTGTTTCTTT), obtained from Integrated DNA Technologies. The PCR protocol included an initial denaturation at 98°C for 30 seconds, followed by 10 amplification cycles consisting of denaturation at 98°C for 10 seconds, annealing at 62°C for 15 seconds, and extension at 72°C for 15 seconds. A final 5-minute extension step at 72°C was performed to complete the last cycle. The PCR products were then separated via electrophoresis on a 2% agarose gel at 150 V for 1 hour. Subsequently, the DNA was purified from the gel using the ZymoClean Gel DNA Recovery Kit (GenScript Scientific #11-301C).

### Generating DNA templates through primer assembly and PCR amplification

The sequences for the wild type, aptamer alone, and four allosteric regime variants were generated using primer assembly at https://primerize.stanford.edu/. The primers were then ordered from IDT at 100 μM in 1x IDTE pH 8.0 buffer. The sequence of each primer is documented in Table S2. For each construct, the middle primers encompassing the P2-P5 sequences were combined and diluted to 1 μM in 100 μL of RNase-free UltraPure water (ThermoFisher #10977015). The reaction mixture for each construct was prepared with 19 μL of RNase-free UltraPure water, 2 μL of each of the first and last primers at 100 μM, 2 μL of the diluted middle primer solution, and 25 μL of Platinum II Hot-Start PCR master mix (ThermoFisher #14000012). Primer assembly began with an initial denaturation at 94°C for 30 seconds, followed by 30 cycles of denaturation, annealing, and extension. The denaturation, annealing, and extension phases were conducted for 30 seconds at 94°C, 60°C, and 72°C, respectively. A final 5-minute extension step at 72°C was then performed to complete the PCR cycle. To verify the correct sizes of the amplified products, the products were separated on a 2% agarose gel (150V, 1 hour) and subsequently purified using a ZymoClean Gel DNA Recovery Kit (GenScript Scientific #11-301C).

### In vitro transcription of RNA constructs

Each construct was transcribed *in vitro*. We prepared a 10x transcription buffer containing 400 mM Tris-HCl (pH 8.0), 10 mM spermidine, and 0.1% Triton X. Next, we prepared the transcription reaction mix. The transcription reaction mix contains 10 µL 10x transcription buffer, 5 µL 50 mM DTT, 16 µL 25 mM NTPs, 8 µL 250 mM MgCl_2_, 4 µL T7 polymerase (provided by the Eichhorn lab at UNL), 24 µL template DNA (adjusted to 0.3 µM), and 33 µL RNase-free water. We incubated the transcription run at 37 °C for 6 hours. We then digested the DNA template with DNase I and purified the RNA using an RNA Clean and Concentrator-5 kit (Genesee Scientific #R10 14). After purification, we used a Nanodrop to measure RNA concentration and ran a 4% agarose gel (150V, 1 hour) to confirm that the RNA was the correct length.

### Modifying RNA constructs using DMS-MaPseq

Each purified RNA construct was treated with DMS. For each condition, we started with 10 pmol of purified RNA in 1.75 μL of RNase-free water. Each RNA sample was denatured at 90 °C for 4 minutes, then allowed to cool for 3 minutes at 4 °C prior to folding. The folding and modification of RNA with DMS conditions will vary depending on the specific titration being performed. AMP titration conditions included 17 concentrations, evenly spaced on a logarithmic scale between 0 and 100 µM: 0, 0.1, 0.16, 0.25, 0.40, 0.63, 1.0, 1.58, 2.51, 3.98, 6.31, 10, 15.84, 25.11, 39.81, 63.09, 100 µM, with MgCl_2_ maintained at a final concentration of 10 mM. Given that Switch-On exhibits weaker AMP binding, it was titrated across a broader range of seventeen points up to 1,000 µM: 0, 0.35, 0.59, 1.01, 1.72, 2.92, 4.96, 8.43, 14.34, 24.37, 41.43, 70.43, 119.73, 203.54, 346.02, 588.24, and 1,000 µM. Magnesium titration was conducted following the same protocol under optimized conditions: a mixture of 1.75 μL RNA with folding solution, comprising 400 mM Sodium cacodylate buffer (18.75 μL) and 250 mM MgCl_2_ (1 μL), was prepared to achieve the desired concentrations. To achieve final concentrations of 50 mM Sodium cacodylate and 10 mM MgCl_2_, 18.75 μL of 66.67 mM Sodium cacodylate was combined with 1 μL of the target MgCl_2_ concentration. Subsequently, 1 μL of 2.5 mM AMP (to reach a final concentration of 100 μM AMP) or water (in the absence of AMP) was added and vortexed. RNA folding occurred in the thermocycler at 22 °C for 30 minutes, except for the Switch-On variant, for which we increased the temperature to 44 °C.

The DMS solution was prepared by mixing 15 μL of DMS (Sigma-Aldrich #D186309) with 85 μL of 100% ethanol (Decon Labs #2716). Immediately following the 30-minute incubation, 2.5 μL of DMS solution was added, and the mixture was incubated for 6 minutes. The reaction was quenched by adding 25 μL of BME (ThermoFisher #1254700). The modified RNA was purified utilizing the RNA Clean & Concentrator-5 kit (Genesee Scientific #R1014) and eluted in 7 μL of RNase-free water. Subsequently, 1 μL of the purified RNA sample was combined with the Qubit RNA BR Assay Kit (ThermoFisher #Q10211) for quantification.

TGIRT III reverse transcriptase (provided by the Eichhorn Lab at UNL) was used to convert methylation events into mutations in the cDNA strand during reverse transcription. The reverse transcription reaction mix consisted of 2.4 μL 5x TGIRT buffer, 1.2 μL 10 mM dNTPs, 0.6 μL 100 mM DTT, 0.5 μL TGIRT-III enzyme, 6.4 μL modified RNA diluted to 0.25 μM, and 1 μL 0.285 μM barcoded RTB RT primer (**Table S2**). The 5x TGIRT buffer consisted of 250 mM Tris-HCl pH 8.3, 375 mM KCl, and 15 mM MgCl_2_. Once the reaction mix was made, the sample was incubated for 2 hours at 57°, hydrolyzed with 0.4 M NaOH (5 μL), heated for 4 minutes at 90°, and cooled for 3 minutes at 4°. Quench acid was used to neutralize the NaOH. Quench acid consists of 1.43 M NaCl, 0.57 M HCl, and 1.29 M sodium acetate. 2.5 μL was used, but this varied depending on the batch made to neutralize the NaOH. The reaction was diluted to 50 μL by adding 30 μL of RNase-Free water, which is the minimum volume for the purification kit. Purification was conducted utilizing the Oligo Clean and Concentrator Kit (Genesee Scientific #11-380B). cDNA was eluted with 15 μL RNase-free water. Purified cDNA was then amplified using PCR. PCR reaction mix totaled 50 μL and consisted of: 25 μL NEBNext High-Fidelity 2X PCR Master Mix (NEB #M0541S), 18 μL RNase-free UltraPure water (ThermoFisher #10 977015), 2 μL cDNA, 2.5 μL forward primer, and 2.5 μL reverse primer. The forward primer sequence was: AATGATACGGCGACCACCGAGATCTACACTCTTTCCCTACACGCGCTCTTCCG. The reverse primer sequence was: CAAGCAGAAGACGGCATACGAGATCGGTCTCGGCATTCCTGCTGAACCGCTCTTCCGATCTGGAACAGCACTTCGGTGCAAA. PCR conditions for this run followed those of the oligo pool, except that 16 cycles were used for denaturation instead of 10. Bands of the correct sizes were visualized after running the products through a 2% E-gel EX Agarose Gel (Bio-Rad), and a Zymoclean Gel DNA Recovery Kit (GeneSyn Scientific #11-301C), respectively. 2% E-gel was run for 10 minutes on the E-Gel Power Snap Plus system (Thermo Fisher #G9 301). Finally, concentrations of the final DNA samples were measured using a Nanodrop and Qubit 1X dsDNA High Sensitivity Assay Kit (ThermoFisher #Q33230).

### Generating DMS reactivity from DMS-MaPseq sequencing data

We sequenced the DMS-MaPseq libraries on Illumina NovaSeq 6000 and NextSeq 1000 instruments using 300-cycle kits run in paired-end mode. We demultiplexed each run using the unique barcode added during reverse transcription to distinguish each experimental condition, using Novobarcode (https://www.novocraft.com/documentation/novobarcode/).

`novobarcode -b rtb_barcodes.fa -f test_R1_001.fastq test_R2_001.fastq`

The barcode file rtb_barcodes.fa lists one barcode per line, for example:

`Distance 4`

`Format 5`

`RTB021 CCAATGGGTGTA`

`RTB022 AGCCAAAACTGG`

`RTB023 GTGTGTTTGCCC`

`Distance` is the number of base differences allowed between a barcode and a read for assignment, and `Format 5` places the barcode at the 5′ end of read 1. This separated each experimental condition into unique demultiplexed FASTQ files. The demultiplexed FASTQ files are available at the Sequence Read Archive under accession PRJNA1483681.

Processing the demultiplexed FASTQ files into mutation fractions was performed using the rna-map software (https://github.com/YesselmanLab/rna_map) (41). With the following command for each replicate

`rna-map -fa <Fasta file> -fq1 <R2 fastq file> -fq2 <R1 fastq file> -- dot-bracket <csv file>`

Supplying each construct’s sequence and dot-bracket structure maps every read to the correct construct. For each position, `rna-map` reports the fraction of reads carrying a mismatch or deletion, which, under DMS, reflects N1-methyladenine and N3-methylcytosine adducts (42). We used these raw mutation fractions as the reactivity throughout, without library-wide normalization, and required at least 2,000 reads per construct in each condition; constructs below this depth were dropped before analysis.

### Classifying library variants into allosteric regimes

The mutation fraction of the GAAA tetraloop was calculated as the average of the mutation fractions of the three loop adenines. The mutation fraction of the tetraloop-receptor is represented by its sole reporter adenine in the receptor. We found it useful to merge these metrics into a single quantity, MF(TL/TLR), defined as the average of the mutation fractions of the tetraloop and the tetraloop-receptor (39). MF(TL/TLR) serves as a reporter for the state of the tertiary contact: when docked, these adenines are protected (low MF); when undocked, they are exposed (high MF). The mutation fraction of the aptamer, MF(apt), is calculated as the average of the mutation fractions of the two reporter adenines in the ATP-binding pocket (39, 43). For each construct tested in the ± AMP screen, we calculated the change in docking upon addition of AMP. We refer to this quantity as ΔMF(TL/TLR), where ΔMF(TL/TLR) = MF(TL/TLR, −AMP) - MF(TL/TLR, +AMP), the difference in mutation fraction of the TL/TLR in the absence and presence of AMP.

We classified the 3,375 junction variants into allosteric regimes using two cutoffs, one for TL/TLR docking and one for switching. For our switching cutoff, we determined the noise on ΔMF(TL/TLR) from the 3,590 helix-background variants that contain no junction mutations and were much less likely to switch. Their standard deviation was σ = 0.00096, and we designated a variant as AMP-responsive only if |ΔMF(TL/TLR)| was above 3σ = 0.00287. A variant is classified as Switch-On if ΔMF(TL/TLR) ≥ 3σ; that is, AMP causes docking by lowering reactivity. It is called Switch-Off if ΔMF(TL/TLR) ≤ −3σ; that is, AMP causes undocking by increasing reactivity. For variants that were not responsive to AMP, we used our second cutoff between Always-On and Always-Off. We assigned θ_low_ as the cutoff for Always-On, having low reactivity regardless of AMP, similar to the Wild-Type. We set θ_low_ = 1.1 × MF(TL/TLR of wild-type) = 0.0031. We set θ_high_ as the cutoff for when a contact is considered undocked. We set θ_high_ equal to the midpoint between the value for the average mutation fraction of helical variants (MF_helix_ = 0.0036) and the value for a fully undocked control (MFcontrol = 0.026, the mean of two controls that are incapable of forming the contact, one a UUCG tetraloop and the other a receptor deletion). This midpoint is θ_high_ = MFhelix + 0.5 × (MF_control_ − MF_helix_) = 0.015. A non-responsive variant is therefore Always-On below θ_low,_ Always-Off if its resting reactivity is at or above θ_high_, and Unclassified if its resting reactivity falls between θ_low_ and θ_high_, where we cannot assign it a state unambiguously.

### Fitting aptamer AMP affinity by the Hill equation

We fit the mutation fraction of the two aptamer reporter adenines relative to AMP concentration using a three-parameter Hill equation with the Hill coefficient fixed at one, expressed as MF(L) = MF_0_ + (MF_1_ − MF_0_) * (L / (Kd + L)), where MF_0_ and MF_1_ represent the ligand-free and saturated mutation fraction asymptotes respectively, K_d_ denotes the apparent dissociation constant, and L signifies the AMP concentration. The hill coefficient was fixed to 1 because only a single AMP molecule is bound by the aptamer. Before fitting, we averaged the mutation fraction over the replicates. We then fit by nonlinear least squares (scipy curve_fit). We fixed the coefficient at one because the aptamer binds a single AMP. Before fitting, we averaged the mutation fraction over the replicates that passed quality filters for each condition. We then fit by nonlinear least squares (scipy curve_fit). The fit is iterative and requires a starting value for each parameter, which we took from the data: the minimum and maximum mutation fractions (MF_0_ and MF_1_) from the 5th and 95th percentiles of the observed mutation fraction, and K_d_ from the AMP concentration closest to the midpoint of the mutation fraction. Because a poor starting value can lead to a poor fit on the wrong answer, we ran it from several starting values and kept the one with the smallest error. We also held each parameter to a physically sensible range: the two reactivities within the values we measured, and K_d_ positive and within the tested concentrations. To obtain a 95% confidence interval for each parameter, we resampled the fit residuals and refitted 500 times, each time starting from the point estimate, then took the 2.5th and 97.5th percentiles of the resulting distribution.

### Linked-equilibrium modeling of ligand-docking coupling

To quantify the degree of coupling between AMP binding and TL/TLR docking, both the mutation fraction titration curves of the TL/TLR and ATP aptamer were fitted to a four-state model (44, 45). This model comprises two elements, each possessing two states: the contact between TL/TLR (docked or undocked) and the aptamer (AMP-bound or unbound). These states collectively yield four configurations. The equilibrium is characterized by three parameters: the propensity for docking in the absence of AMP (K_TL0_), the affinity of the aptamer for AMP when the contact is undocked (K_AMP0_), and the coupling factor α, which determines the extent to which AMP binding influences docking. Specifically, α = 1 indicates no effect; α > 1 suggests AMP promotes docking; α < 1 implies AMP opposes docking. DMS mutation fraction depends on the state, as the reporter adenines are protected and yield low readings when the contact is docked or the aptamer is AMP-bound, whereas they are exposed and yield high readings when undocked or the aptamer is free. The model captures this behavior through four mutation fraction parameters: r_D_ (docked) and r_U_ (undocked) for the TL/TLR, and r_B_ (AMP-bound) and r_F_ (free) for the aptamer, translating the fraction of RNA in each state into a predicted signal. This results in seven parameters that are simultaneously fitted to both curves using nonlinear least-squares optimization (scipy.optimize.least_squares). The coupling is expressed as a free energy change, ΔΔG_C_ = −RT ln α (with R = 1.987 × 10⁻³ kcal mol⁻¹ K⁻¹; T = 295.15 K, or 317.15 K for Switch-On at 44 °C). The same fitting process yields an apparent AMP affinity, K_d_ = (1 + K_TL0_) / [K_AMP0_ (1 + α·K_TL0_)]. Consistent with Hill coefficient fits, reactivity values were averaged across replicates that passed quality control prior to fitting. Confidence intervals at 95% were estimated for α, ΔΔG_C_, and K_d_ via 500 samplings of the residuals followed by refitting.

### Measuring coupling direction from magnesium titrations

For each construct, we fit the −AMP and +AMP TL/TLR titration curve with two models: a two-state monotonic docking curve (undocked to docked) and a three-state model (unfolded to folded to docked). To decide what model to use, we applied the small-sample Akaike information criterion ΔAICc. We selected the three-state model only when ΔAICc < -2 and a nested F-test yielded a p-value < 0.05. Based on these criteria, we found that the wild-type and Always-On are better fit by a two-state model, but the rest fit better to a three-state model. We obtained the [Mg^2+^]_1/2_ from the midpoint of the two-state model and from the second transition in the three-state curves. The AMP-induced fold-shift is the log2 ratio of the +AMP and −AMP midpoints, and its sign gives the coupling direction (a shift to higher Mg^2+^ is negative coupling) (46, 47). We estimated its error by a robust bootstrap (2,000 resamples): the standard error is 1.4826 times the median absolute deviation of the bootstrap log2 fold-shifts, and the 95% interval is taken from the 2.5th and 97.5th percentiles of the bootstrap distribution.

### Per-helix energetic features

For the helix-background analysis (Supplemental Results 1), we computed three per-helix features. Per-helix folding free energy was the duplex ΔG of each helix, from a Turner 2004 nearest-neighbor RNAcofold of its two strands (ViennaRNA (40)), including duplex initiation and terminal AU/GU penalties, so values are comparable across variants. Per-helix DMS reactivity was the sum of mutation fraction over the A and C positions in each helix, taken with and without AMP and as their difference. Per-helix G·U wobble count was the number of G·U/U·G pairs in each duplex.

## Results

### AMP binding and tertiary docking in the characterized RNA nanostructure are thermodynamically coupled but do not switch

To build a 3D RNA switch, we started with a rationally designed nanostructure that we characterized previously (**Figure 1A**) (39). This minimal construct contains an ATP aptamer linked to a tetraloop/tetraloop receptor (TL/TLR) contact via a kink-turn and a 4-1 junction. (**Figure 1A**). The TL/TLR scaffolding preorganizes the bound conformation of the ATP aptamer, stabilizing its bound state (39). We performed DMS-MaPseq, which monitors both elements simultaneously: TL/TLR docking shields four adenines in the tetraloop and receptor, whereas AMP binding lowers the mutation fraction (MF) at two adenines in the aptamer core (**Figure 1B**). To measure the apparent binding affinity of AMP, we performed a 17-point titration, yielding a K_d_ of 0.46 ± 0.10 µM, down from 2.58 ± 0.63 µM for the aptamer alone, consistent with prior measurements (**Figure 1C, Supplemental Figures S1-S6**).

**Figure 1:**
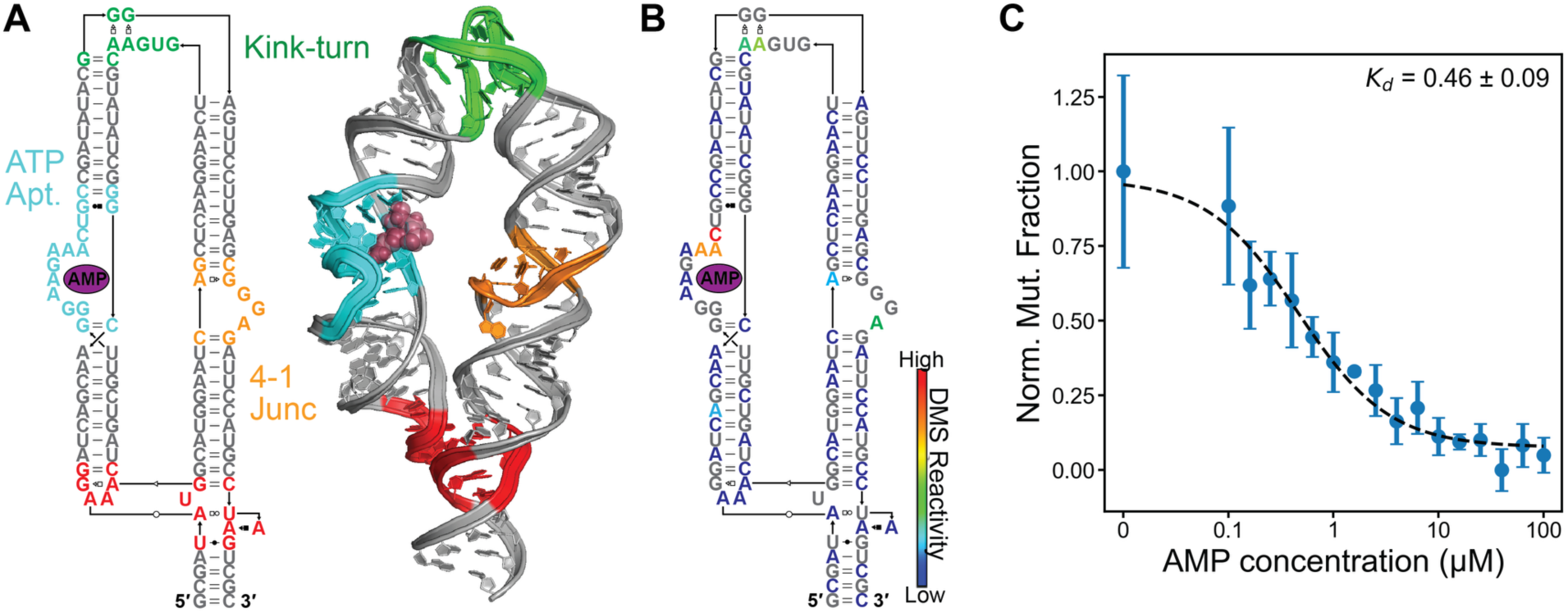
Overview of the rationally designed AMP-responsive 3D-switch RNA. (A) Secondary structure (left) and 3D model (right) of the wild-type construct, colored by motif: kink-turn (K-turn, green); ATP-binding aptamer (cyan); 4-1 junction (orange); and the peripheral GAAA tetraloop / tetraloop-receptor (TL/TLR, red) tertiary contact. The magenta oval marks the position of bound adenosine monophosphate (AMP) in the aptamer pocket. Secondary structure of the same construct, with each nucleotide colored by its per-nucleotide (B) DMS reactivity at saturating AMP (color scale at right: blue = low, yellow/orange = mid, red = high). Low reactivity at the GAAA tetraloop adenines (TL/TLR contact, bottom). (C) AMP titration of the wild-type construct measured by aptamer DMS reactivity (normalized to 0–1), fit with a three-parameter Hill equation (dashed line). Apparent K_d_ = 0.46 ± 0.09 μM. Points are replicate means; error bars are standard deviation across replicates.

Thermodynamic coupling is necessary for switching but not sufficient. Across the titration, aptamer and TL/TLR mutation fractions were uncorrelated (R² = 0.027; **Supplemental Figure S7**): the TL/TLR remained docked regardless of whether AMP was bound. This indicates the wild-type scaffold is thermodynamically coupled to ligand binding but too stable to switch. Here, the kink-turn and 4-1 junction tune the free-energy landscape that connects the aptamer and TL/TLR. In the wild-type scaffold, these junctions favor a low-energy ensemble in which the TL/TLR is formed and the aptamer is in a bound-like pose. Mutations in these motifs will alter their lowest-energy conformation, thereby changing the global conformation in the absence of AMP. The free energy of AMP binding could then bias the equilibrium, driving the TL/TLR to dock or undock.

### A massive library screen reveals four allosteric regimes

To systematically search for switching variants, we designed a library of 7,149 variants targeting the two junctions and the surrounding helices. We generated two types of variants. Helix-only variants that only changed the WC pairs in each helix. We also produced junction variants with a single or double mutation at the kink-turn and 4-1 junctions, as well as helix sequence changes, to increase sequence diversity (see Methods). We profiled the library by DMS-MaPseq at ±100 µM AMP. We selected constructs with more than 2,000 aligned reads across the ± AMP conditions, yielding 6,965 sequences for analysis.

To score whether the TL/TLR is docked or not, we created a TL/TLR average mutation fraction. We took the mean of the three adenines in the GAAA tetraloop and the receptor’s A that forms an A·A pair with the tetraloop, as we previously used (39) (See methods). To define a cutoff between docked and undocked, we used two controls that cannot form the TL/TLR interaction (the GAAA tetraloop replaced with a non-docking UUCG and the receptor replaced with a helix; Methods) (**Supplemental Figure S8**). We defined undocked as any mutation fraction greater than 50% of the TL/TLR mutation fraction in these two controls. To classify switching behavior, we computed a ΔMF(TL/TLR), which is the mutation fraction of the TL/TLR without AMP minus the TL/TLR mutation fraction with AMP, so a positive ΔMF(TL/TLR) means AMP stabilizes docking. We generated two class names to describe the docking state of the TL/TLR contact: “Always” variants maintain a single state across both AMP conditions, whereas “Switch” variants are responsive to AMP binding. Variants with ΔMF ≈ 0 and a low TL/TLR mutation fraction are Always-On (the contact stays docked in both AMP states); those with ΔMF ≈ 0 and a high mutation fraction are Always-Off (it stays undocked). Variants with ΔMF > 0 are Switch-On (positive cooperativity: AMP binding biases the equilibrium toward docked); variants with ΔMF < 0 are Switch-Off (negative cooperativity).

We first tested whether helix sequence alone could switch the contact, since helix changes can modulate tertiary-contact stability (43, 48, 49). No helix-only variant produced contact-specific ligand coupling. Of the 3,590 helix-only constructs, 110 exceeded the Switch-On threshold, but each carries a signature of weak global folding: less stable helices, more G·U wobbles, and elevated ΔMF summed across every helix rather than localized at the TL/TLR (**Supplemental Results 1; Supplemental Figure S9**). This distributed signal reflects AMP-induced stabilization of the entire secondary structure in marginally folded variants, not allosteric control of the tertiary contact. The ΔMF(TL/TLR) distribution of the helix-only pool therefore defines the experiment’s noise floor (σ = 0.00096), and we set the switching cutoff at three times this width (0.00287).

We next examined the 3,375 junction variants, comprising kink-turn-only, 4-1-only, and ones with mutations in both junctions. Unlike the helix-only constructs, the junction variants spanned all four allosteric regimes under the cutoffs described above (**Figure 2A-B**): 1,097 Always-On (32.5%) and 21 Always-Off (0.6%) hold a single docking state across both AMP conditions, whereas 237 Switch-Off (7.0%) and 120 Switch-On (3.6%) respond to the ligand. The remaining 1,900 variants (56.3%) had resting reactivities between our docked and undocked cutoffs and could not be assigned to a single state. Rather than experimental noise, these likely represent partially docked ensembles: their reactivities are distributed continuously between the Always-On and Always-Off populations, and the magnesium titrations below resolve a partially docked intermediate in three of four representative constructs.

**Figure 2:**
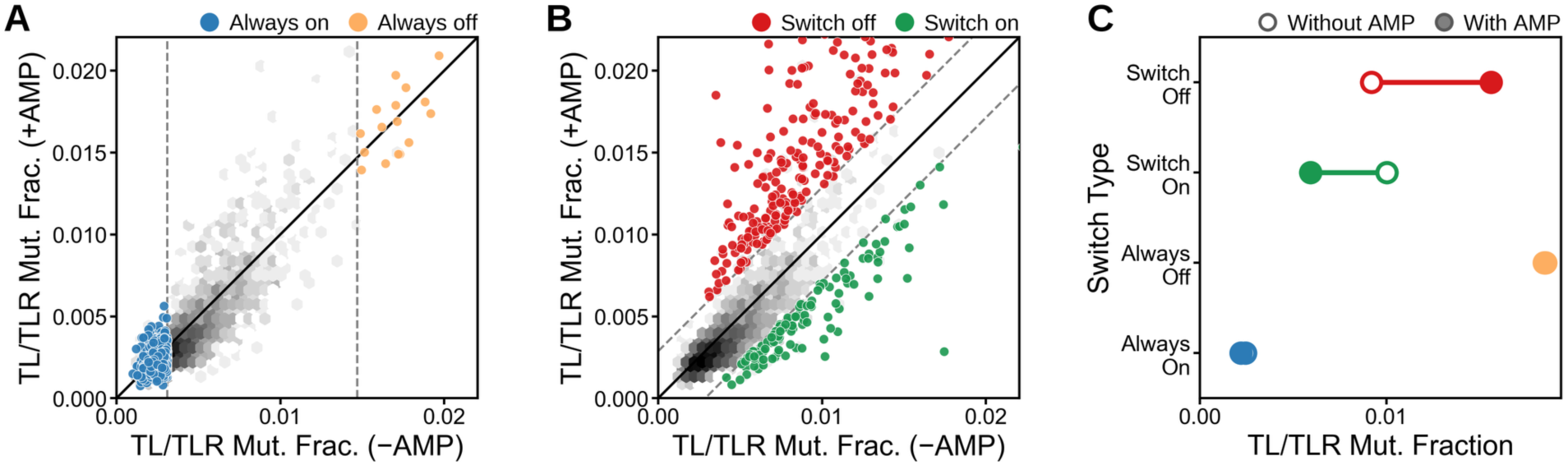
Junction mutations fall into four allosteric regimes. Each junction variant is placed by its TL/TLR reactivity without AMP versus with AMP (mean of the GAAA tetraloop and receptor docking reporters). The solid line is y = x, and the gray density shows all 3,375 junction variants. (A) Always-On (blue) and Always-Off (orange) variants. The dashed verticals are the baseline cutoffs that split the non-switching variants into docked (Always-On, low baseline) and undocked (Always-Off, high baseline). (B) Switch-Off (red) and Switch-On (green) variants. The dashed diagonals are the switching cutoffs on ΔMF(TL/TLR). Variants above the upper diagonal lose docking on AMP binding (Switch-Off, negative cooperativity), and variants below the lower diagonal gain it (Switch-On, positive cooperativity). (C) Representatives TL/TLR reactivity per class without AMP (open) and with AMP (filled), with a line marking the AMP-induced change. Always-On and Always-Off stay flat, Switch-Off rises, and Switch-On falls.

### The kink-turn, not the 4-1 junction, drives allostery

To determine which mutations alter allosteric behavior, we grouped our variant data into bins by mutation location: kink-turn only, 4-1 junction only, and both regions. Across all variant classes exhibiting divergent allostery relative to wild-type (Always-On), the kink-turn category was enriched. Of Switch-Off variants (N=237), 90.7% were kink-turn only (N=215), 8.9% were double mutants (N=21), and 0.4% were 4-1 only (N=1). Of Always-Off variants (N=21), 81.0% were kink-turn only (N=17), 19.0% were double mutants (N=4), and zero were 4-1-only. For Switch-On variants (N=120), 57.5% were kink-turn only (N=69), 37.5% were 4-1 only (N=45), and 5.0% were double (N=6). Thus, based on these variant populations, allosteric behavior was primarily determined by mutations in the kink-turn region, which was therefore subjected to further analysis.

The kink-turn is a conserved motif consisting of two tandem sheared A·G pairs adjacent to a three-nucleotide bulge (see **Supplemental Figure S10** for kink-turn residue numbering). It can switch between linear and kinked (120°) conformations, and the populations of these two states depend on metal ions, temperature, and protein binding (50–52). Based on the cryo-EM structure, the kinked state is required for TL/TLR docking (53). Mutations within the two sheared A·G pairs (1b·1n, 2b·2n) have been previously shown to disrupt the ability of the kink-turn to bend in solution (51, 54, 55). Thus, a mutation at these positions likely alters the preferred lowest-energy structure from a kinked to a linear conformation, inhibiting docking. Mutation of position A35 (1b) was found to be the predominant Switch-Off site: 88% of single mutants made at this position are Switch-Off (A35U: 95%, A35C: 86%, A35G: 80%) (**Supplemental Figures S11, S12**). However, all four regimes are present at this position at much lower populations: Always-On (5%), Switch-On (3%), and Always-Off (4%). Switch-On appears when mutations are made at 2b or bulge position 34 (G34A: 38%). Mutations at the remaining bulge positions (32, 33) are Always-On. G34U is the only bulge mutation that is not predominantly Always-On and leads to Switch-Off. The Always-Off regime is rarely observed and occurs most frequently with A35C, converting 1b·1n to a Watson-Crick C·G pair, or with G93A, converting 2b·2n to an A·A pair (10%). Mutations within the kink-turn, largely those in the conserved A·G, account for most of what dictates which regime a variant falls into. However, changes to base pairs in the helices also affect distribution; unfortunately, no single feature within the helices can predict a mutant’s regime.

### Kink-turn substitutions tune the sign and magnitude of effective coupling

Accurate quantification of the sign and magnitude of coupling between TL/TLR docking and ligand binding necessitates an AMP titration. Conducting sufficient sequencing to encompass the entire library for this titration would require substantial sequencing depth; therefore, we selected a representative from each allosteric class. The Always-On representative is a G109C variant (a mutation at the 4-1 junction) that preserves the native kink-turn sequence. The Always-Off variant is an A35C mutation that transforms the 1b·1n sheared A·G base pair into a Watson-Crick C·G pair. The Switch-Off variant includes A35G, which converts the 1b·1n into a noncanonical G·G pair, along with an additional neutral G109C mutation (see Methods). The ‘Switch-On’ variant is an A36U mutation, which converts the 2b·2n sheared A·G into a naturally occurring kink-turn variant, a U·G wobble (52) (**Figure 2C, Supplemental Figure S13)**.

The Always-On variant exhibits a titration curve comparable to that of the wild-type; its TL/TLR mutation fraction remains flat and low throughout as a function of AMP concentration (**Supplemental Figures S14-16**). Conversely, the TL/TLR mutation fraction was high across the Always-Off AMP titration, indicating that it largely remained undocked as AMP concentrations increased (**Supplemental Figures S17-19**). The Switch-Off construct is docked at 0 µM AMP; however, as AMP concentration rises, the mutation fraction at TL/TLR increases, suggesting that AMP induces undocking of the contact (**Supplemental Figures S20-S22**). The Switch-On construct exhibits opposite behavior; however, it yields an inconsistent signal at 22 °C (**Supplemental Figures S23-S26**). To amplify the effect size, we ran the AMP titrations at 44 °C, where we observed a much more consistent decrease in the TL/TLR mutation fraction as AMP increased (**Supplemental Figures S27-S30**). Increasing the temperature did not alter the wild-type’s behavior, which retained its Always-On state (**Supplemental Figures S31-S32**). In all constructs, the mutation fraction of the aptamer decreased as AMP saturated its binding pocket, demonstrating that all constructs successfully bound AMP and that variations were attributable to the interactions of their TL/TLR contacts.

To convert the titration curves into a singular coupling number, we simultaneously fit both the TL/TLR docking and AMP binding curves to a four-state linked-equilibrium model (see Methods) (44, 45). This model assumes that both events, the TL/TLR formation (docked or undocked) and the aptamer binding (AMP-bound or unbound), are coupled. One parameter, α, measures the extent to which one process influences the other: α = 1 indicates independent switches, α > 1 signifies positive cooperativity (where AMP stabilizes docking), and α < 1 indicates negative cooperativity (where AMP destabilizes docking). This coupling constant is expressed as a free-energy change, ΔΔG_C_ = −RT ln α (see Figure 3C). 95% confidence intervals are derived from a residual bootstrap (see **Figure 3**, Methods).

**Figure 3:**
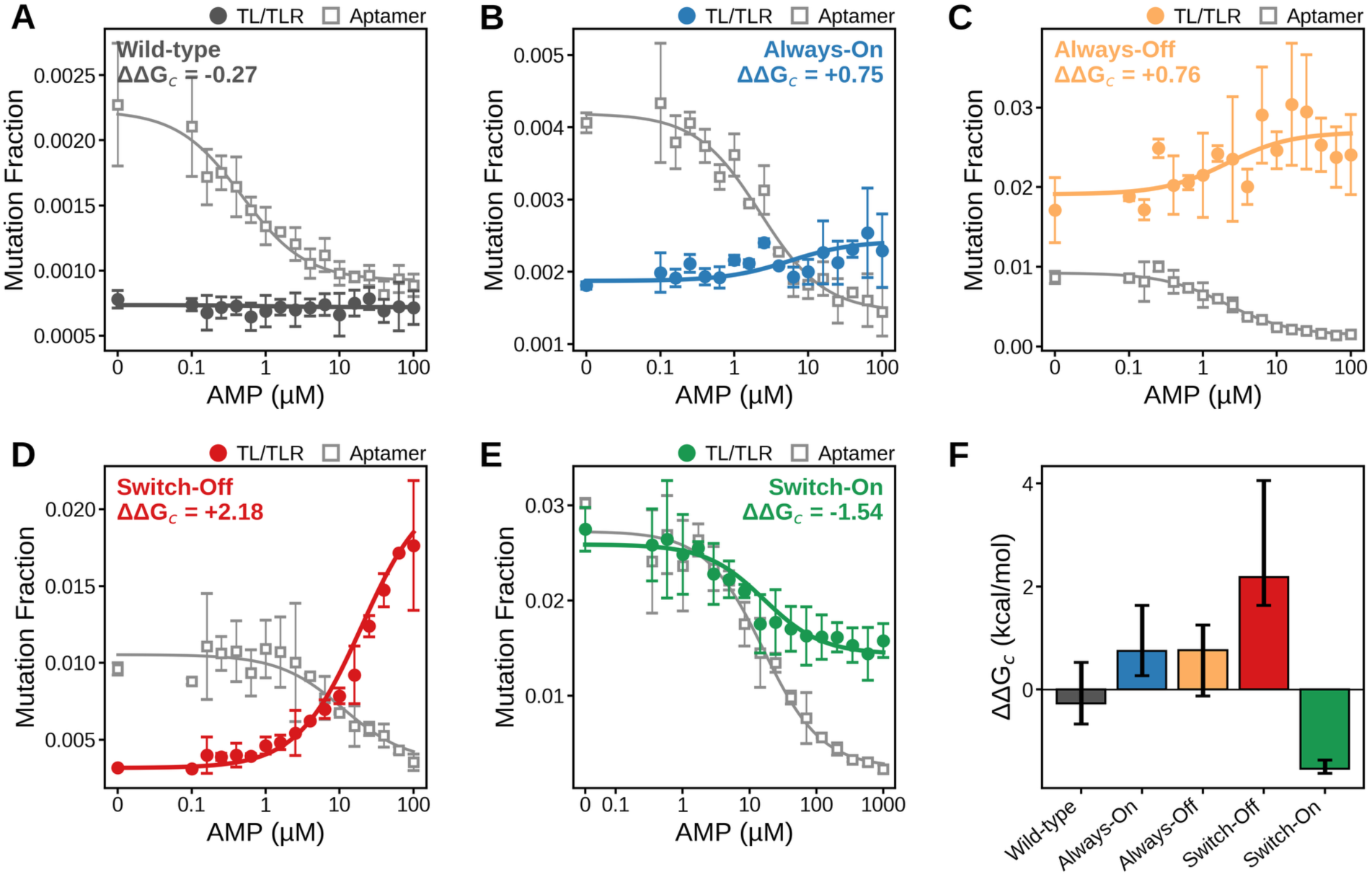
AMP titrations resolve the sign and magnitude of coupling between ligand binding and tertiary docking across the four regimes. (A to E) DMS-MaPseq AMP titrations for the wild-type and the four representative mutants. Filled symbols are TL/TLR docking reactivity (mean of the GAAA tetraloop and receptor reporter adenines); open gray squares are ATP-aptamer mutation fraction (the binding-site adenines). A higher mutation fraction marks the undocked (TL/TLR) or ligand-free (aptamer) state. Aptamer mutation fraction falls as AMP rises in every construct, confirming AMP binding in all five so AMP binding drives docking. Curves are model fits, and the effective coupling free energy ΔΔG_C_ = −RT ln α, where α is the coupling factor, is annotated on each panel. Points are replicate means; error bars are standard deviation across replicates. Switch-On (E) was profiled at 44 °C over a wider AMP range (to 1000 μM). (F) ΔΔG_C_ (kcal/mol) for all five constructs, colored by regime.

As a baseline for comparison, the wild-type exhibited an α value of 1.58 (ΔΔG_C_ = −0.27 kcal/mol), indicating weak positive coupling, with AMP stabilizing the TL/TLR contact. This observation is consistent with prior research on this nanostructure (39). The Always-On and Always-Off variants demonstrate weak negative coupling, with α values of 0.28 (+0.75 kcal/mol) and 0.27 (+0.76 kcal/mol), respectively. Both are far smaller than the switching variants, and the Always-On magnesium titration gives a fold shift whose interval includes 1 (**Figure 4F**), so its residual coupling is weak. For the switching constructs, the effects observed were markedly stronger. The Switch-Off configuration yielded an α of 0.024 (+2.18 kcal/mol) at 22 °C, corresponding to a fortyfold reduction in docking induced by AMP. Conversely, the Switch-On configuration exhibited an α of 11.4 (−1.54 kcal/mol) at 44 °C, indicating an elevenfold increase. Across all four regimes, ΔΔG_C_ varies by approximately 3.7 kcal/mol.

**Figure 4:**
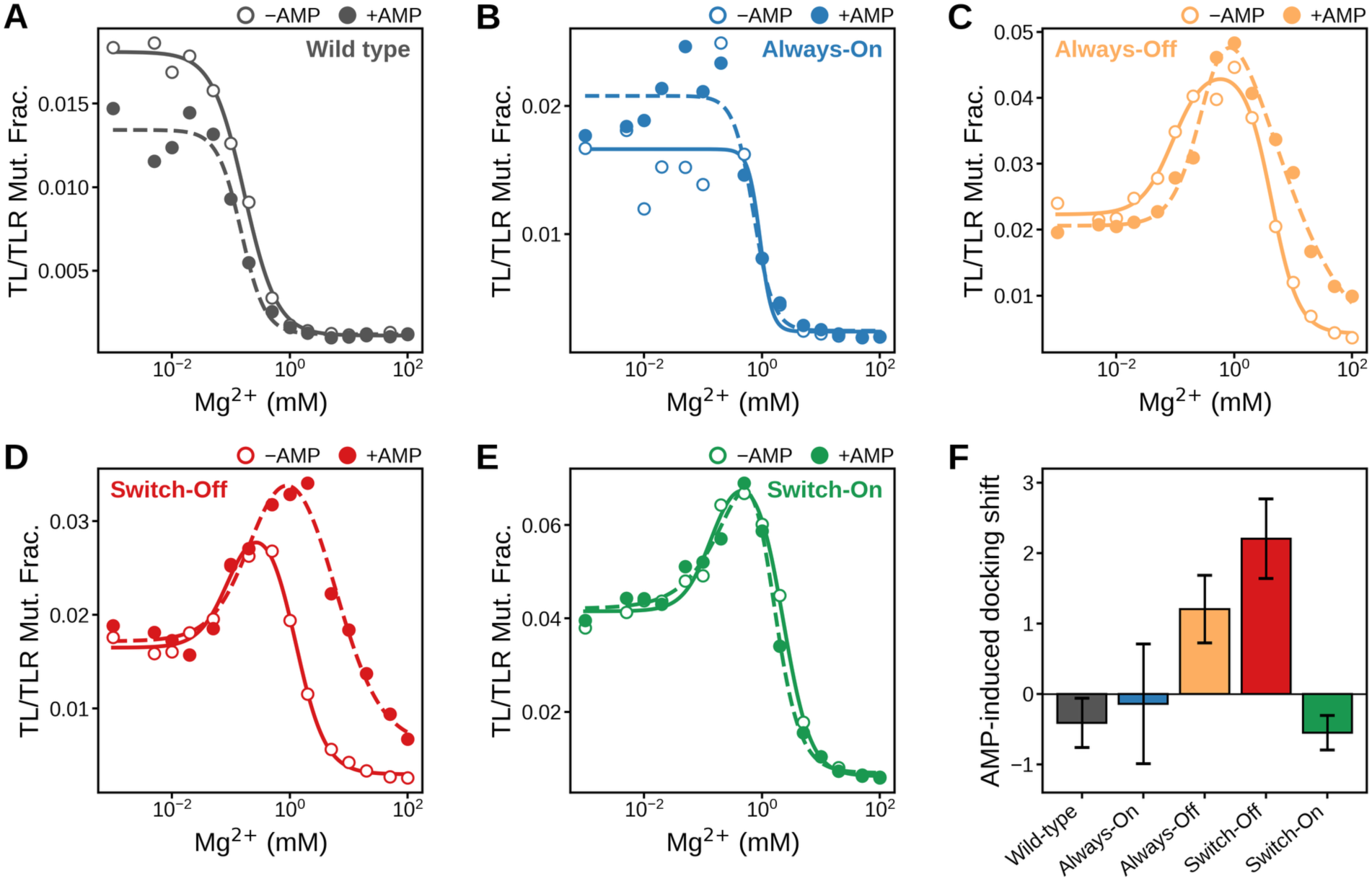
Magnesium titrations reshape the docking profile and independently confirm the coupling sign. (A to E) TL/TLR docking reactivity versus Mg^2+^ for the wild-type and the four representative mutants, without AMP (open symbols, solid fit) and with saturating AMP (filled symbols, dashed fit). A higher mutation fraction marks the undocked state; raising Mg^2+^ drives folding and docking. Curves are fits of a three-state model (A↔B↔C). Wild-type and Always-On titrate monotonically and change little with AMP. Always-Off and Switch-Off are non-monotonic, and AMP shifts their docking transition to higher Mg^2+^, the signature of negative coupling (AMP opposes Mg-driven docking). Switch-On (measured at 44 °C) binds AMP, seen as protection of the aptamer reporter, but its TL/TLR Mg²⁺ titration is nearly AMP-independent and does not independently resolve a coupling sign. Points are replicate means and error bars the standard deviation across replicates. (F) AMP-induced shift of the docking midpoint, as the signed log2 ratio of the model-free [Mg^2+^]_1/2_ with AMP over without AMP.

We also calculated an apparent ligand-binding Kd from standard two-state Hill fits. The values follow trends in the coupling. Always-On and Always-Off bind at 2.2 µM and 2.1 µM, respectively, values close to the aptamer-alone value of 2.58 µM (**Supplemental Figure S33**). Switch-Off and Switch-On bind AMP over 10-fold weaker than wild-type and about 5-fold weaker than the aptamer alone at 11.1 µM and 16.0 µM, respectively. These increases suggest that some of the AMP binding energy is diverted toward breaking (Switch-Off) or stabilizing (Switch-On) the TL/TLR contact. The linkage model returns an apparent K_d_ of its own, serving as an independent check on the coupling constant (**Supplemental Figure S33**). The two are in agreement except in the flat wild-type and Always-On cases, where there is negligible movement of docking upon AMP binding. There, the model K_d_ is underdetermined and its confidence interval is large (0.96 versus 0.46 µM; 5.6 versus 2.2 µM); we report the direct Hill value in those cases. These two-state Hill fits are strong additional evidence of the linked-equilibrium model.

### Magnesium titrations confirm the coupling sign and reveal three-state docking in the kink-turn mutants

Magnesium titrations gave an orthogonal test of the AMP-derived coupling. Mg^2+^ stabilizes the TL/TLR and drives the kink-turn into its kinked state, and we previously demonstrated that AMP lowers the Mg^2+^ required to dock (39). We ran a 16-point Mg²⁺ titration from 0 to 100 mM on the wild type and the four representatives, with and without AMP (**Figure 4**, **Supplemental Figure S34-S45**). The wild type and Always-On titrate as a simple two-state docking (undocked to docked), the model we used before (**Figure 4A, 4B**) (39). The three kink-turn mutants do not follow a two-state model. Always-Off, Switch-Off, and Switch-On all fit a three-state model, in which TL/TLR reactivity first rises at low Mg^2+^ and only then falls as docking sets in (**Figures 4C to E**). The rise suggests the contact is already partially docked at zero Mg^2+^. We do not resolve the intermediate directly, but the simplest path consistent with the curves is partially docked to undocked to docked, in which kink-turn mutations shift the stability of the extended state relative to the kinked state: the extended state forms first and breaks the contact, and above a critical Mg^2+^, the TL/TLR forms and pulls the kink-turn back into the kinked state.

To evaluate how AMP acts on each construct, we compared the magnesium midpoint ([Mg^2+^]_1/2_) with and without AMP (**Figure 4F**). For the three-state fits, we took the second transition, the one that leads to the docked state. Linked equilibria predict that ligand stabilization lowers the Mg²⁺ requirement (positive coupling), destabilization raises it (negative coupling), and no coupling leaves the two curves overlapping (46, 47). As seen before, AMP lowered the wild-type [Mg^2+^]_1/2_ from 0.17 to 0.13 mM (fold-shift 0.75x, 95% CI 0.59 to 0.96) (39). Always-On moved from 0.63 to 0.57 mM (0.91x, CI 0.50 to 1.63), the only construct whose interval includes 1, consistent with docking and ligand binding no longer being coupled. AMP raised [Mg^2+^]_1/2_ in both negatively coupled constructs: Always-Off from 4.08 to 9.40 mM (2.30x, CI 1.65 to 3.21) and Switch-Off from 1.51 to 6.98 mM (4.61x, CI 3.12 to 6.82), the largest shift we measured and consistent with its +2.18 kcal/mol AMP-titration assignment. Always-Off already demands a large amount of Mg^2+^ to dock at all, so AMP acts on an equilibrium that is weak to begin with. Switch-On moved the other way, from 2.67 to 1.82 mM (0.68, 0.58 to 0.81), so AMP helps the TL/TLR form. Every construct, except the Always-On control, resolved a shift whose direction matches the sign of its AMP α (**Figure 3**), providing additional evidence for the coupling we observed with the ligand titrations.

## Discussion

Here we demonstrate that a ligand-responsive switch can be constructed entirely through tertiary rearrangement, and that its response can be flipped by a single base change. Resolution of this coupling behavior into a single term α (effective ΔΔG_c_ = −RT ln α) within a four-state linked-equilibrium framework spans ≈3.7 kcal/mol (44, 45, 56). This term encompasses both the sign of the AMP-induced change in TL/TLR docking and the direction of the AMP-induced shift in the Mg^2+^ midpoint. Tertiary coupling therefore presents an alternative design framework to that of secondary-structure switches in which both the direction and magnitude of response can be tuned in a more modular fashion.

Kink-turn sequence controls the coupling between ligand binding and tertiary docking. Decades of structural and FRET work establish that single-base substitutions at the conserved G·A pairs tune the kinked/extended equilibrium and the Mg^2+^ requirement for kinking (50–52, 54, 55). This study demonstrates that the kinked/extended equilibrium of the kink-turn can be directly exploited to control thermodynamic coupling when it is placed between an aptamer and a tertiary contact. Substitutions that preserve the sheared G·A pairs leave the contact stably docked and not responsive to AMP binding. Substitutions that abolish kinking (G·A to G:C) render it constitutively undocked. Only substitutions that destabilize kinking without abolishing it (G·A to G·G, A·G to U·G) produce ligand-responsive switching, with the sign of the response set by which kink-turn conformation is favored in the absence of ligand. Switching therefore requires the kink-turn to be poised near its conformational midpoint, where the free energy of ligand binding is sufficient to tip the balance.

There is one potential caveat in our coupled linkage model. The model treats the RNA as two binary switches, docking and AMP binding, so α is collapsing all the kink-turn state changes into this term. The kink-turn is itself dimorphic, interconverting between kinked and extended states in response to Mg^2+^ and temperature (50, 52, 54), and our substitutions act primarily on that equilibrium rather than on docking directly. The measured α is therefore an effective coupling that folds the kink-turn conformational change into the docking response. This is the parameter relevant to engineering, since it predicts the change in docking per ligand bound, but it does not separate kink-turn folding from docking. Doing so would require resolving the kink-turn conformational states directly, which our DMS data cannot provide, since the positions that report on kinking are themselves mutated in three of the four representatives. An independent readout of kink-turn geometry, such as single-molecule FRET, would be needed to decompose α into its underlying terms.

To our knowledge, this is the first engineered RNA switch in which a ligand controls a discrete long-range tertiary contact with no change in base pairing, and in which the sign of that control can be inverted by a single substitution. Switching without base-pair rearrangement should avoid the misfolding pathways and kinetic barriers associated with base-pair changes, and because both the TL/TLR contact, aptamer, and kink-turn are modular, the same architecture could be rebuilt in other scaffolds with equivalent junctions. Switch-Off shifts the docking equilibrium 40-fold, corresponding to about 2 kcal/mol, which likely can be improved. The system addresses a broader question: how large RNAs use a free-energy change in one region to control folding in another. This minimal system provides a route to dissecting allostery in natural RNA machines and to designing synthetic ones of greater complexity.

## Data availability

Demultiplexed sequencing reads are deposited at the NCBI Sequence Read Archive under BioProject PRJNA1483681. Processed data sufficient to regenerate every figure and number in this paper, including the classified library of 7,147 variants, the titration data for the wild type and the four representatives, the fitted parameters, and a full data dictionary, are archived at Zenodo (doi:10.5281/zenodo.21843901). The analysis code is available at https://github.com/YesselmanLabPublications/2026_atp_ttr_switch and archived at Zenodo (doi: 10.5281/zenodo.21865199)

## Acknowledgements

We would like to thank Catherine Eichhorn for her thoughtful comments, which strengthened the paper.

## Contributions

J.D.Y designed the project and experiments. A.O. ran the experiments with help from B.K. J.D.Y performed the analysis with the help of A.O. J.D.Y wrote the paper.

## Supplemental Results

### Supplemental Results 1: Helix-only negative control and classification thresholds

Helix sequence can modulate tertiary-contact stability (43), so we screened 3,590 variants in which only the helix base pairs were randomized, keeping the kink-turn, TL/TLR, and aptamer intact. This pool serves two roles. As a noise model, it has a tight ΔMF(TL/TLR) distribution (Methods). As a negative control, it tests whether helix-sequence variation alone can switch the contact.

Under the classification cutoffs, no helix variant crossed the Switch-Off threshold, two crossed Always-Off, and 110 crossed Switch-On. These 112 apparent calls share a structural signature of weak global folding: weaker TL-helix folding free energy (means −11.8 and −10.6 kcal/mol vs −13.3 kcal/mol for the Always-On majority), more G·U wobbles (2.3 and 4.0 vs 1.8 pairs), and elevated total ΔMF summed over all helix A and C positions (Mann-Whitney p < 2 × 10⁻⁵ for each feature in the Switch-On). The breadth of this total-ΔMF signal, present across every helix rather than localized at the TL/TLR, indicates AMP-induced stabilization of the global secondary structure of these weakly folded variants rather than TL/TLR-specific coupling. Helix-sequence variation alone therefore does not generate ligand-induced TL/TLR allostery in this scaffold.

## Supplemental Figures

**Figure S1:**
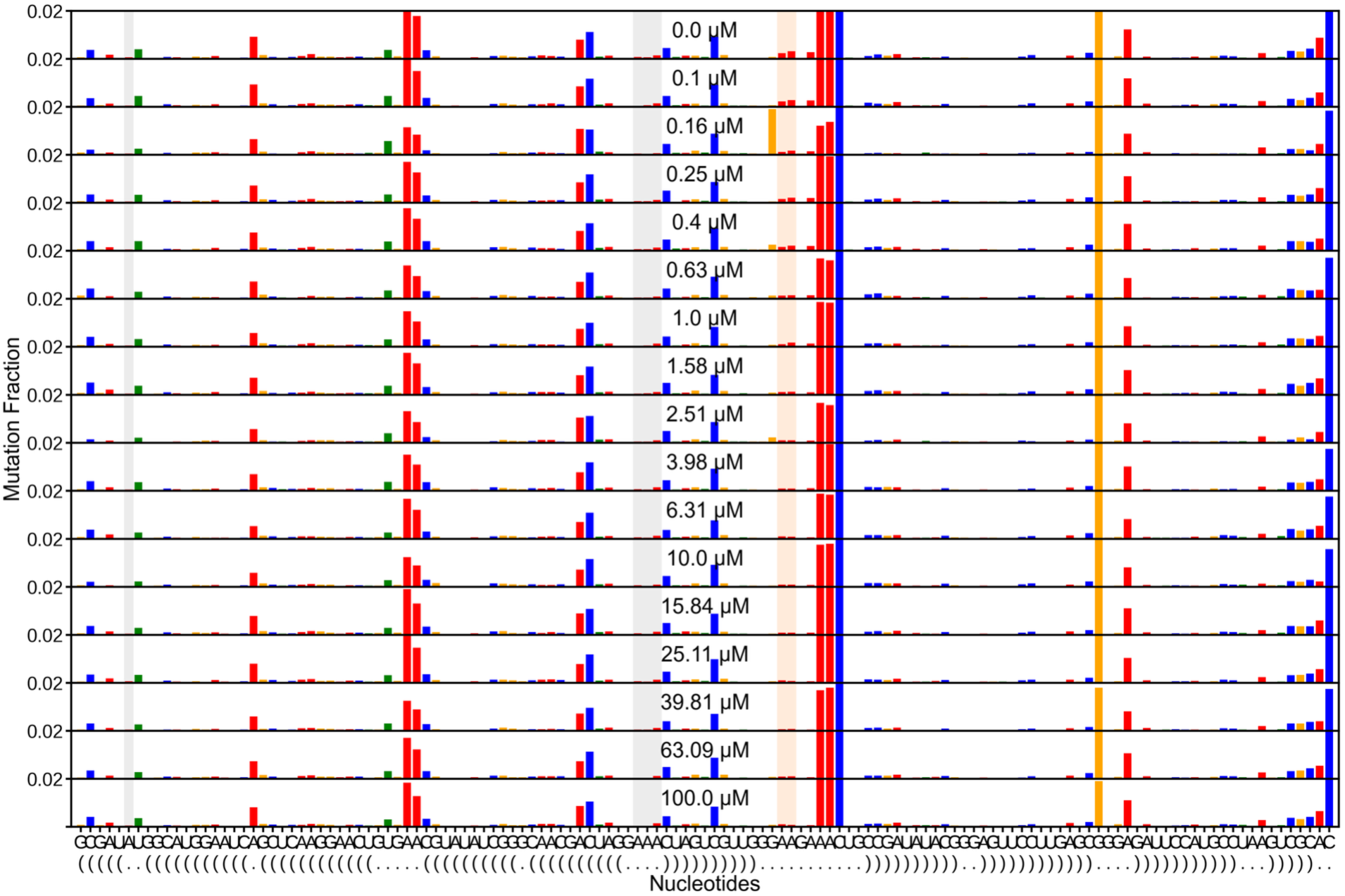
Wild-type per-nucleotide reactivity across the AMP titration, replicate 1. Full per-nucleotide DMS-MaPseq reactivity for the wild-type construct at each point of the AMP titration (replicate 1), stacked from low AMP (top) to high AMP (bottom); each row is the mutation fraction at every nucleotide for one AMP concentration. Gray shading marks the TL/TLR docking reporters (the GAAA tetraloop adenines and the receptor adenine) and orange shading the ATP-aptamer binding-site adenines. Aptamer reactivity falls as AMP rises, reporting ligand binding, while the TL/TLR reporters stay low and roughly constant, so the tertiary contact remains docked across the titration. This is the raw per-replicate data underlying the averaged wild-type curves in Figure 3.

**Figure S2:**
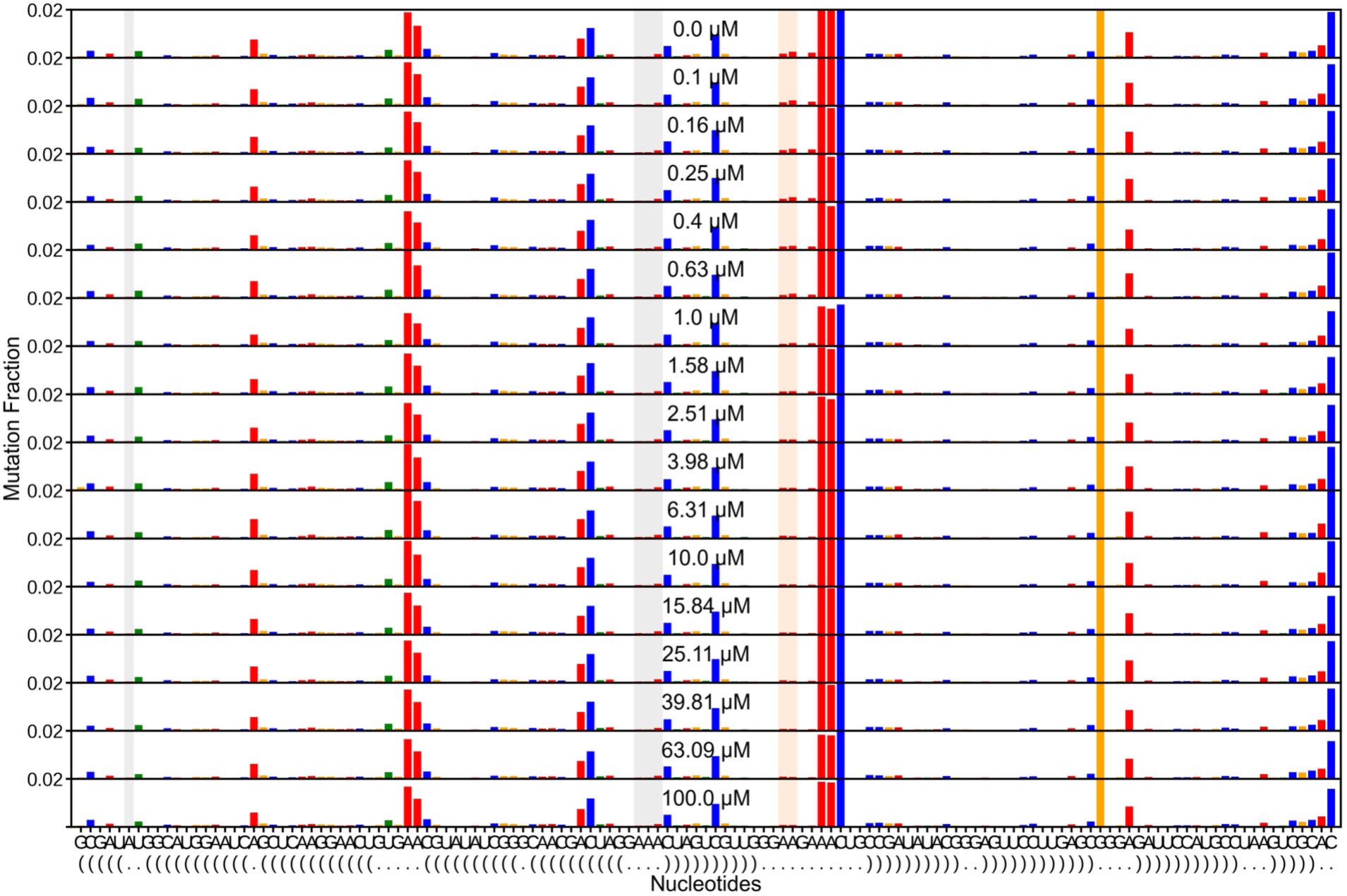
Wild-type per-nucleotide reactivity across the AMP titration, replicate 2. As in Figure S1 for the second wild-type replicate. Full per-nucleotide DMS-MaPseq reactivity at each AMP concentration, with the TL/TLR docking reporters shaded gray and the ATP-aptamer binding-site adenines shaded orange. Aptamer reactivity falls with AMP while TL/TLR reactivity stays low, reproducing the docked, weakly coupled wild-type behavior.

**Figure S3:**
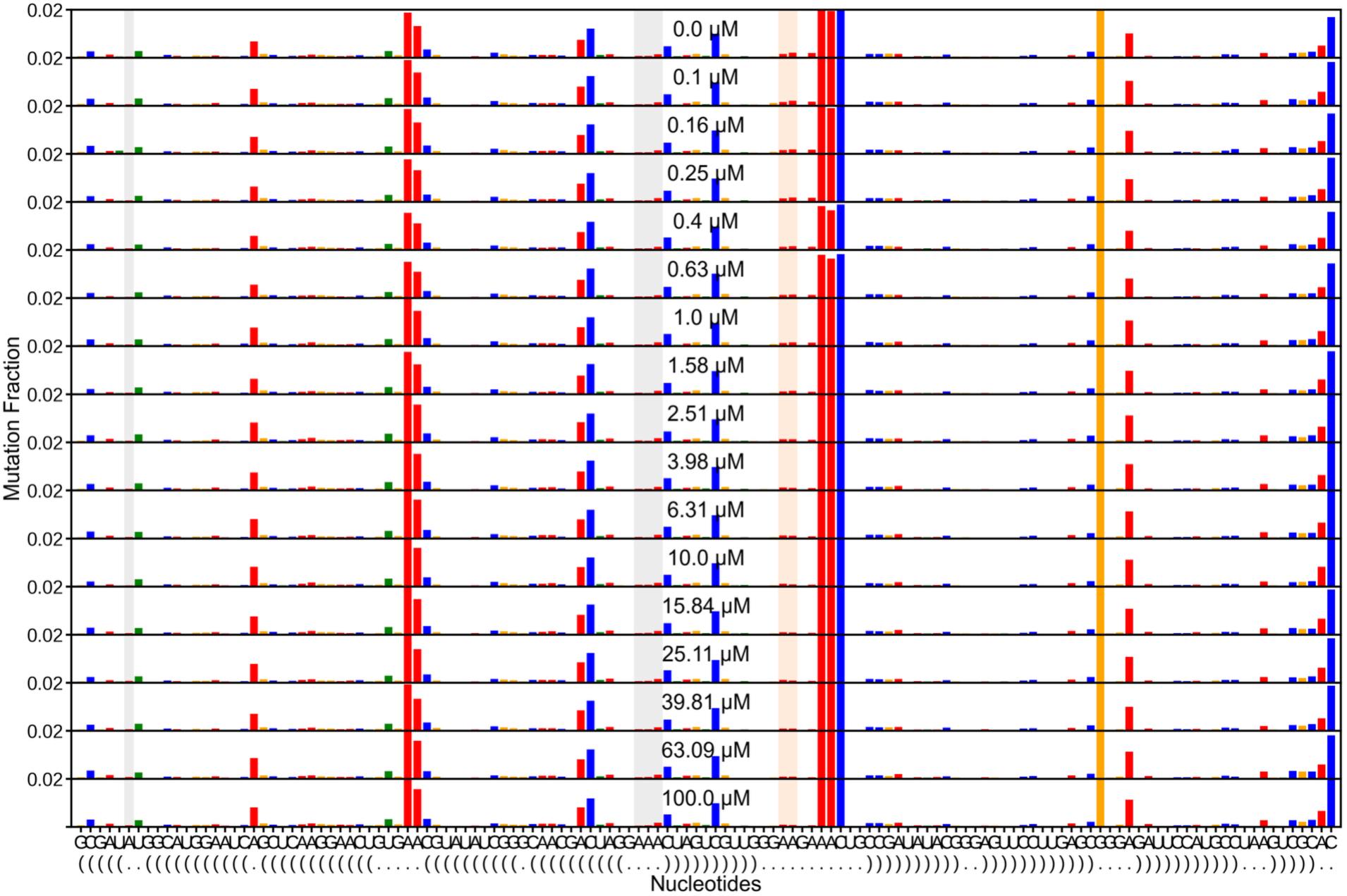
Wild-type per-nucleotide reactivity across the AMP titration, replicate 3. As in Figure S1 for the third wild-type replicate. The aptamer adenines (orange) lose reactivity as AMP increases and the TL/TLR reporters (gray) remain low throughout, consistent across all three replicates.

**Figure S4:**
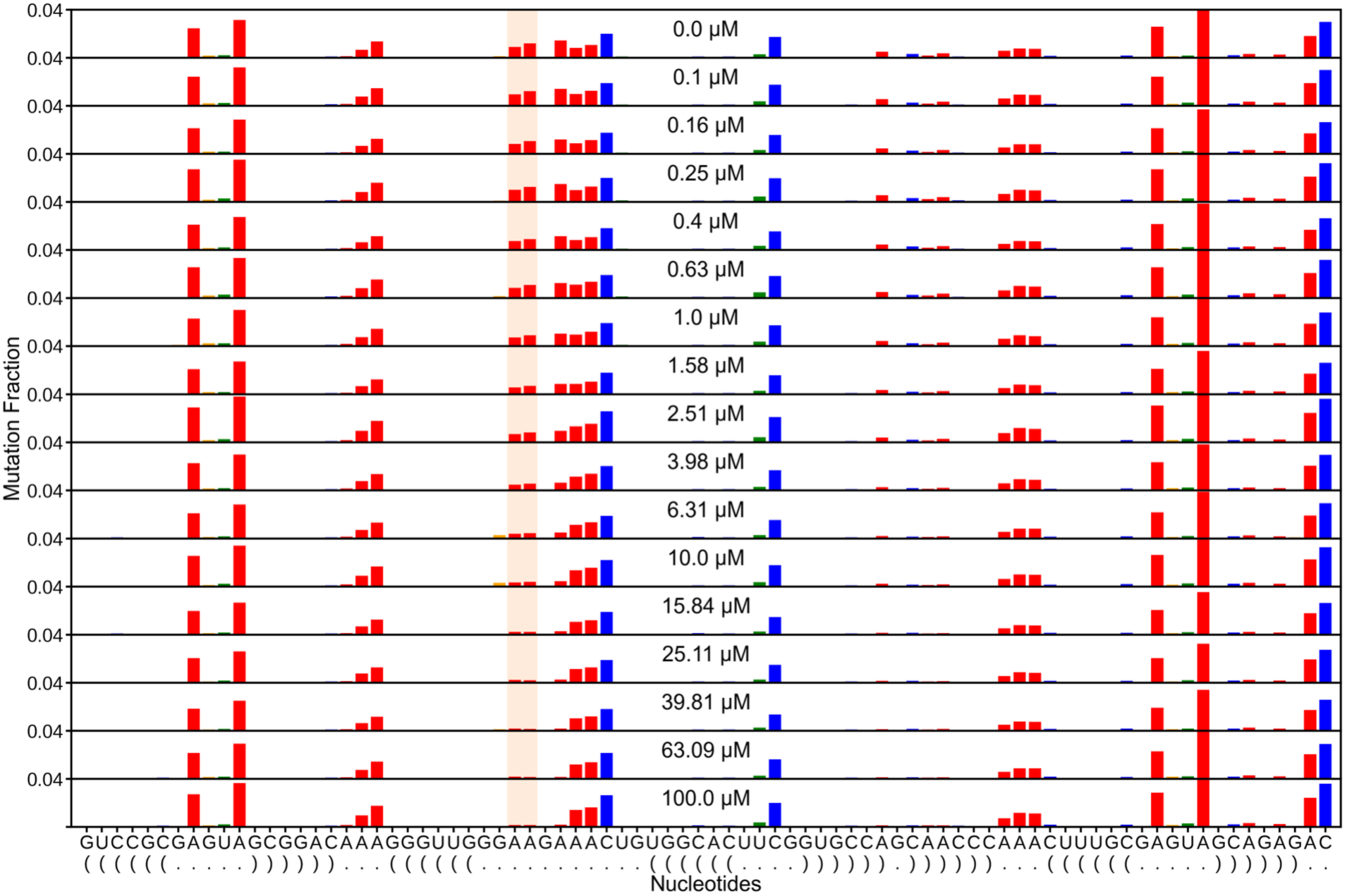
ATP-aptamer-alone per-nucleotide reactivity across the AMP titration, replicate 1. Full per-nucleotide DMS-MaPseq reactivity for the isolated ATP aptamer (no TL/TLR present) at each point of the AMP titration (replicate 1), stacked by AMP concentration. Orange shading marks the binding-site adenines, whose reactivity falls as AMP rises, reporting ligand binding by the aptamer on its own. These per-replicate profiles underlie the aptamer dissociation-constant fit.

**Figure S5:**
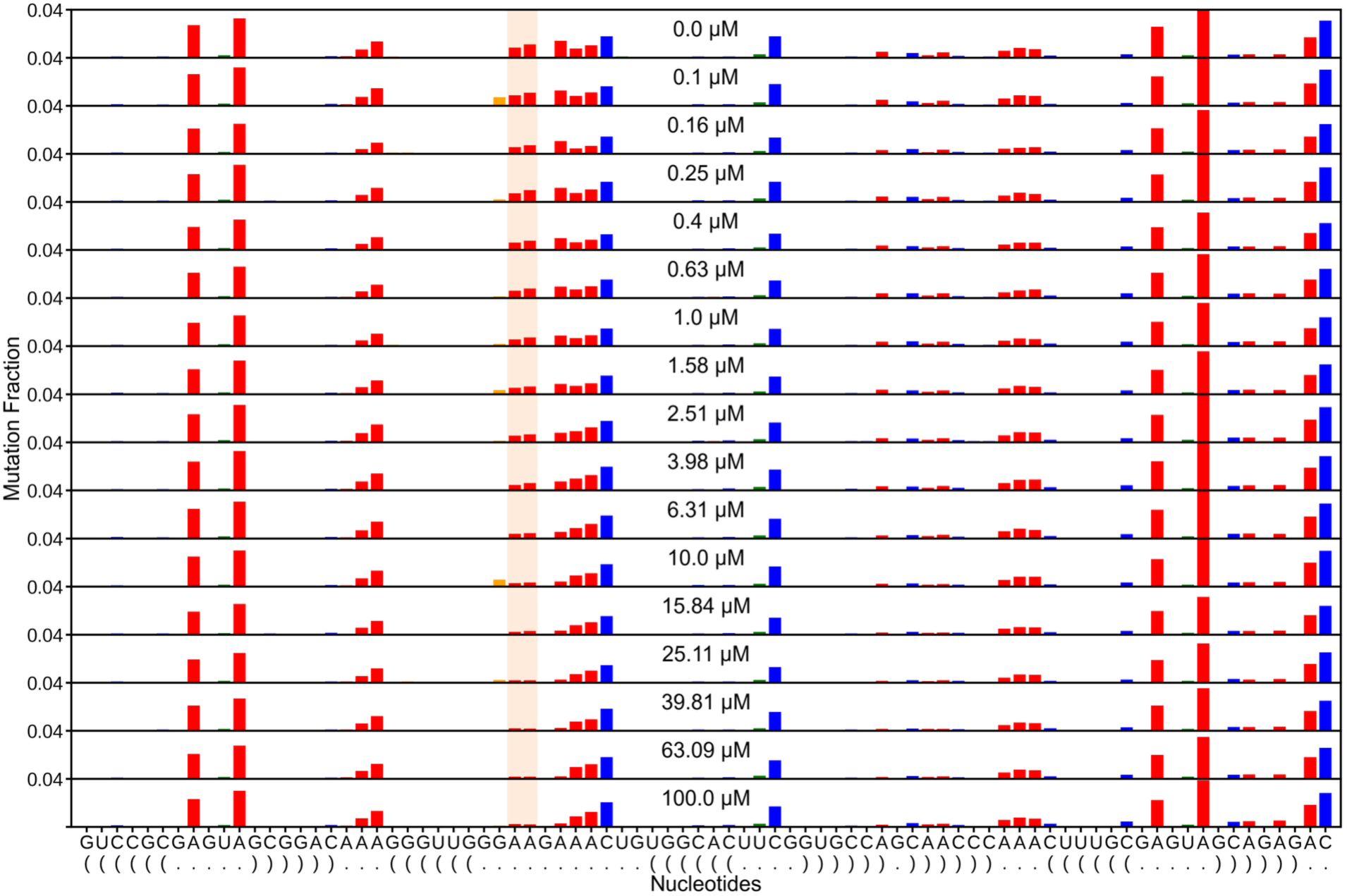
ATP-aptamer-alone per-nucleotide reactivity across the AMP titration, replicate 2. As in Figure S4 for the second ATP-aptamer-alone replicate. The binding-site adenines (orange) are progressively protected as AMP increases, reproducing the isolated-aptamer binding response.

**Figure S6:**
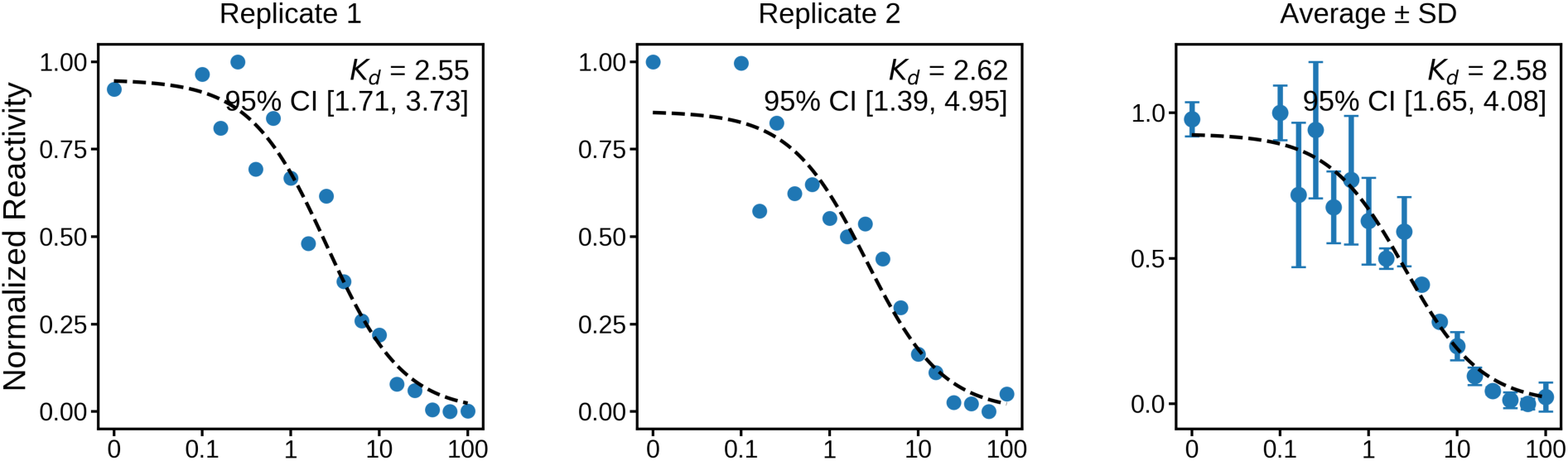
ATP-aptamer-alone AMP binding affinity. Apparent dissociation constant of the isolated ATP aptamer (no TL/TLR). Mean reactivity of the two AMP-sensitive binding-site adenines, normalized to [0, 1], plotted versus AMP concentration for each replicate and for the inter-replicate average (± SD). Dashed lines are three-parameter Hill fits, and the apparent Kd from each fit is annotated. Averaging the replicates gives K_d_ = 2.58 ± 0.67 μM, consistent across replicates (2.55 ± 0.58 μM and 2.62 ± 0.90 μM).

**Figure S7:**
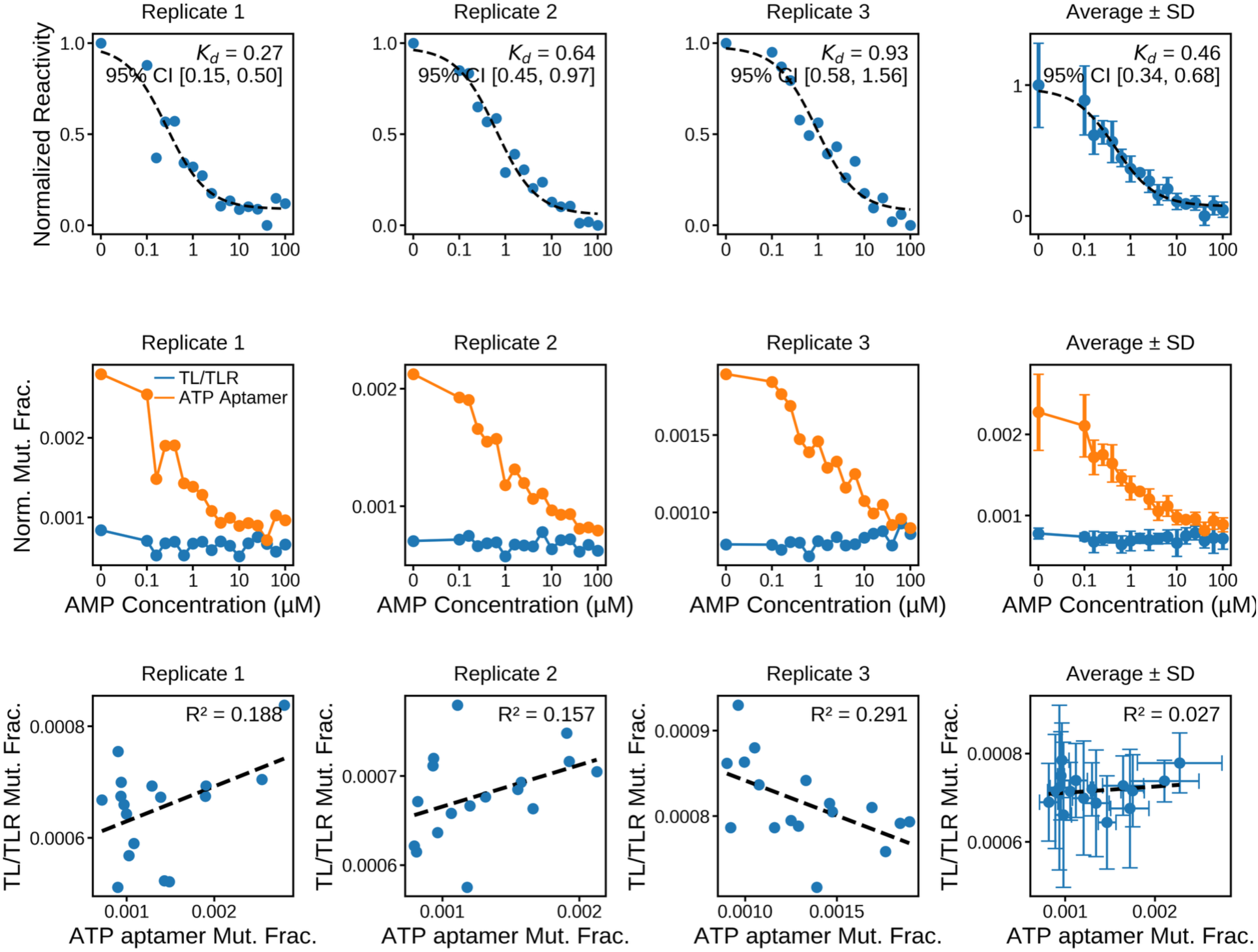
Wild-type AMP binding affinity and weak coupling to TL/TLR docking. (Top) Mean reactivity of the AMP-sensitive ATP-aptamer binding-site adenines, normalized to [0, 1], versus AMP concentration for each wild-type replicate and for the inter-replicate average (± SD). Dashed lines are three-parameter Hill fits, and the apparent Kd from each fit is annotated. The replicate fits give K_d_ = 0.27 ± 0.10, 0.64 ± 0.14, and 0.93 ± 0.24 μM, and averaging the replicates gives K_d_ = 0.46 ± 0.09 μM (the value reported in Figure 1C). (Middle) Per-nucleotide normalized mutation fraction across the same titration for the TL/TLR docking reporters (blue) and the ATP-aptamer binding-site adenines (orange). Aptamer reactivity falls as AMP rises, reporting binding, while the TL/TLR reporters stay low and flat, so the tertiary contact stays docked throughout. (Bottom) TL/TLR docking reactivity versus ATP-aptamer reactivity across the titration, per replicate and averaged, with the coefficient of determination annotated.

**Figure S8:**
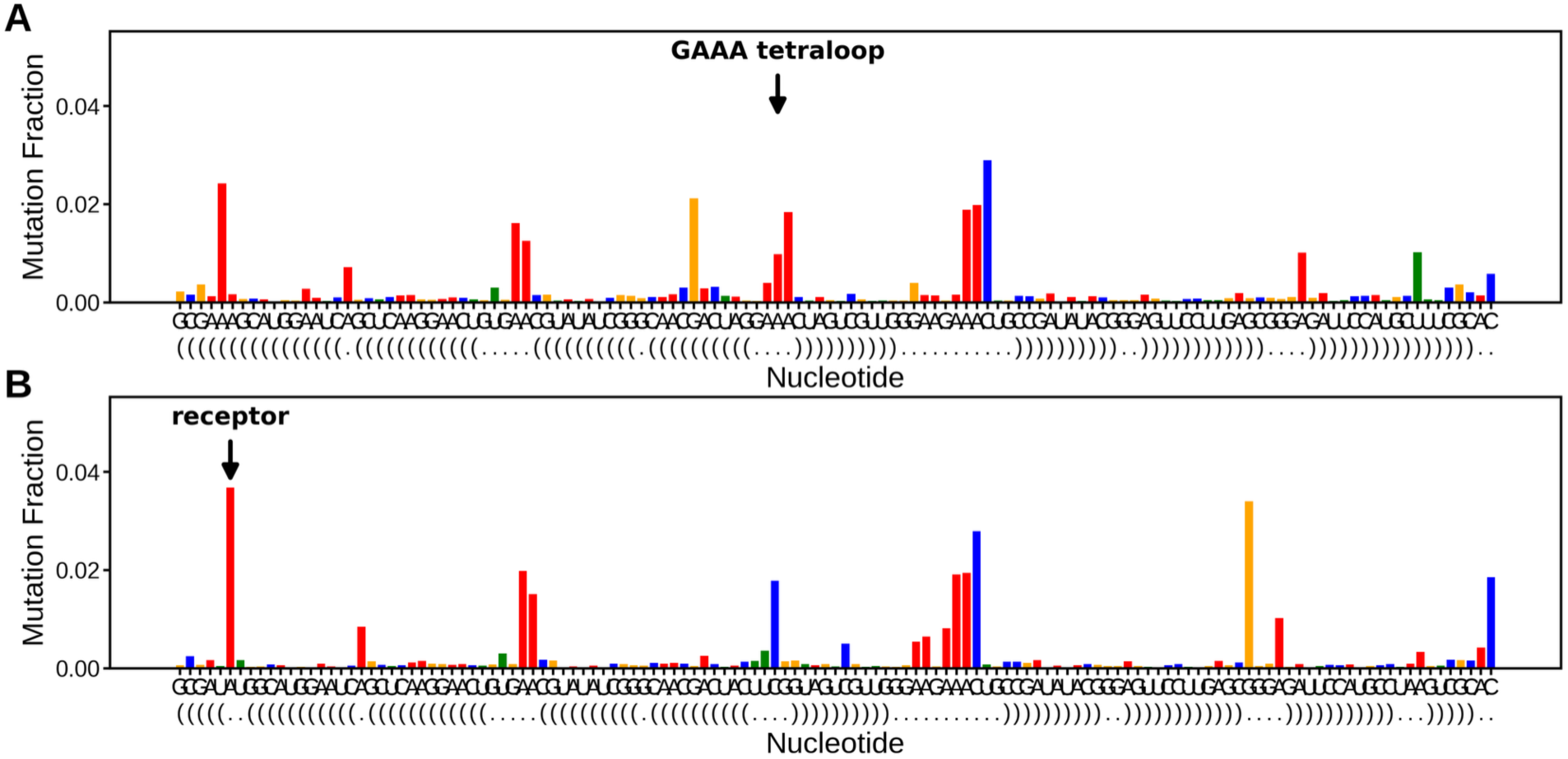
Non-docking controls define the undocked TL/TLR reactivity baseline. Per-nucleotide DMS-MaPseq mutation fraction for control constructs in which the TL/TLR tertiary contact cannot form, so its docking-reporter adenines are fully solvent-exposed. (A) A control probing the GAAA tetraloop (arrow): the tetraloop adenines are highly reactive. (B) A control probing the receptor (arrow): the receptor adenine receptor is highly reactive. These high, undocked reactivities set the reference for a fully open contact and anchor the upper end of the docking scale used to normalize TL/TLR reactivity throughout the study. The dot-bracket secondary structure is drawn beneath each panel.

**Figure S9:**
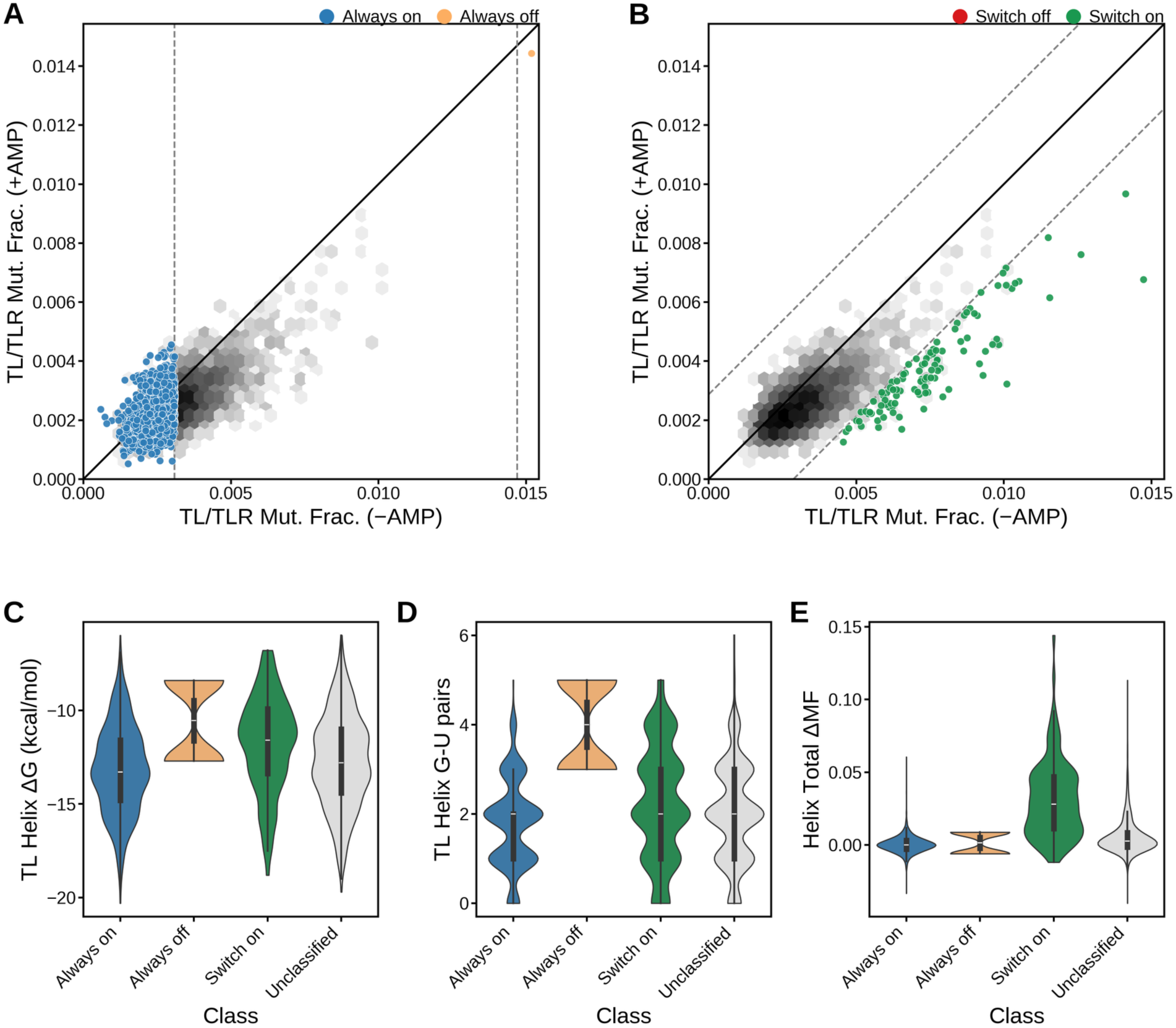
Helix sequence background does not predict switching phenotype. (A and B) TL/TLR reactivity without AMP versus with AMP for all junction variants (gray density), as in Figure 2, with (A) Always-On (blue) and Always-Off (orange) and (B) Switch-Off (red) and Switch-On (green) overlaid. Dashed lines are the same baseline cutoffs (A) and ±3σ helix TL/TLR variation cutoffs (B). (C to E) Distributions of helix-background properties by phenotype class (Always-On blue, Always-Off orange, Switch-On green, Unclassified gray): (C) predicted TL-helix folding free energy ΔG, (D) number of G·U wobble pairs in the TL helix, and (E) total helix ΔMF. The class distributions overlap broadly in ΔG and G·U content, so the randomized helix background does not separate the phenotypes. The only class-linked trend is the elevated helix ΔMF of Switch-On variants in (E). Phenotype is therefore set by junction identity, not by the surrounding helix sequence.

**Figure S10:**
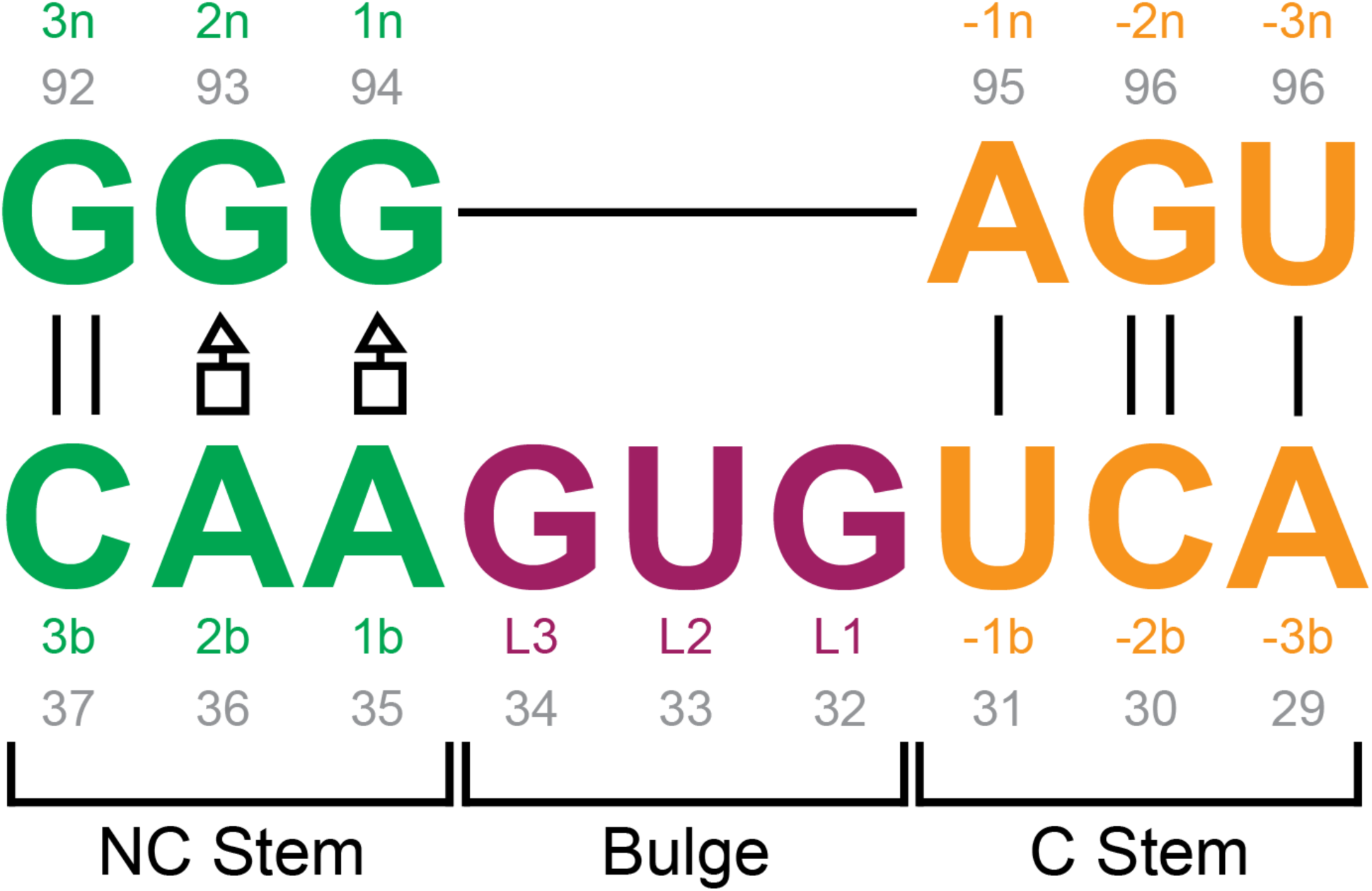
Kink-turn nomenclature and residue numbering. Kink-turn positions in atp_ttr_3, numbered by the standard Leontis-Westhof and Lilley convention. The three-nucleotide bulge is L1 to L3 (positions 32 to 34). Base pairs are counted outward from the bulge: into the NC-stem as 1b·1n (35:94), 2b·2n (36:93), and 3b·3n (37:92), and into the C-stem as -1b:-1n (31:95) and beyond, where b and n denote the bulged and non-bulged strands. The two conserved sheared A·G pairs (1b·1n and 2b·2n) are marked with Leontis-Westhof symbols (open square, Hoogsteen edge; open triangle, sugar edge), and Watson-Crick pairs with vertical lines. Numbers by each base give its atp_ttr_3 residue position.

**Figure S11:**
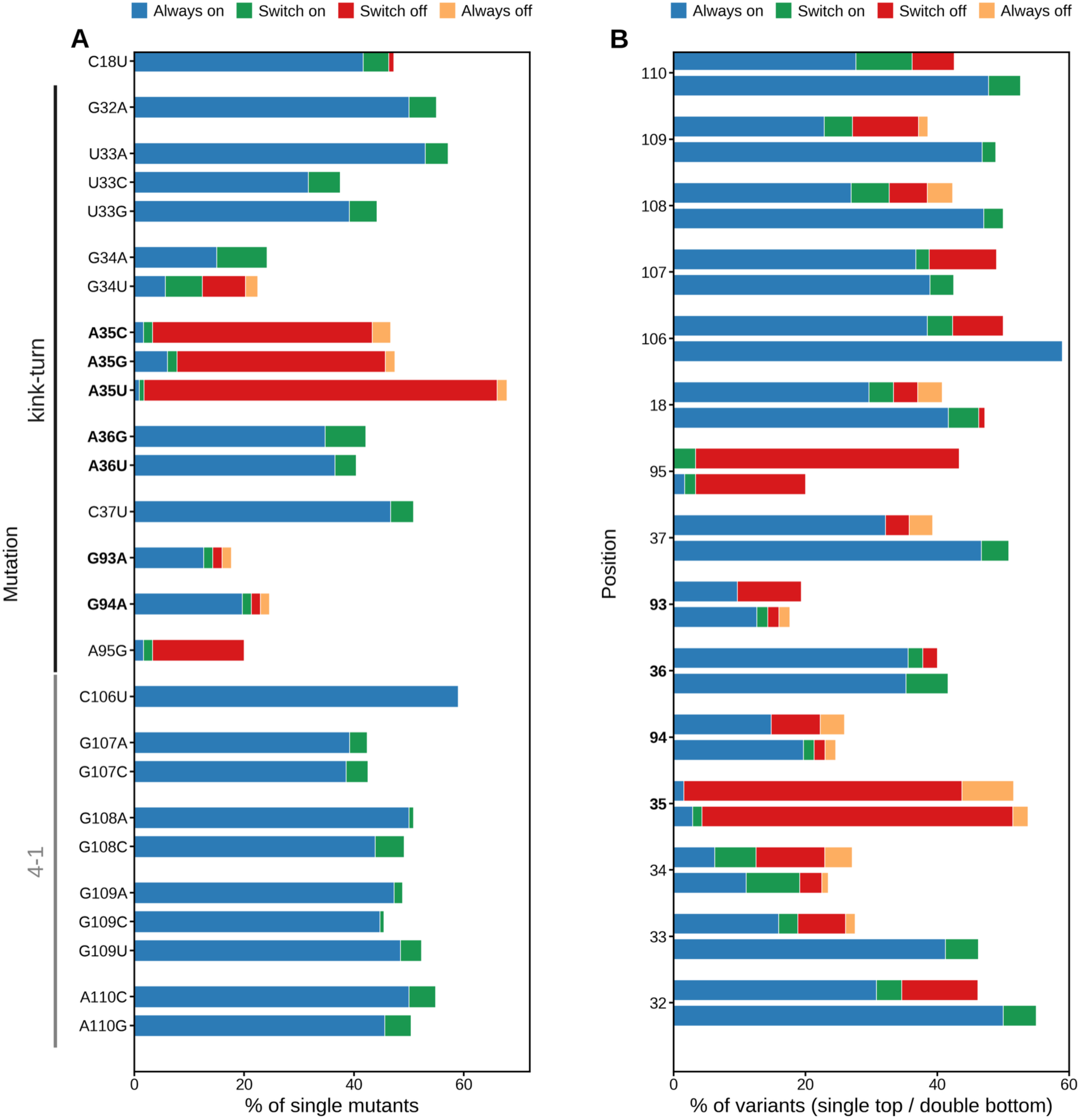
Junction single mutations map to the allosteric regimes by position. (A) Regime composition of each single junction mutation across roughly 120 randomized helix backgrounds, colored by class (Always-On blue, Switch-On green, Switch-Off red, Always-Off orange). Mutations are ordered along the construct and grouped by junction (kink-turn versus 4-1). The kink-turn 1b position dominates: A35C, A35G, and A35U are largely Switch-Off, and A95G at the −1n position is also enriched for Switch-Off, while the 4-1 junction mutations (106 to 110) are almost uniformly Always-On. (B) The same composition per position, comparing single mutants (top bar) with double mutants (bottom bar) that carry a second junction change.

**Figure S12:**
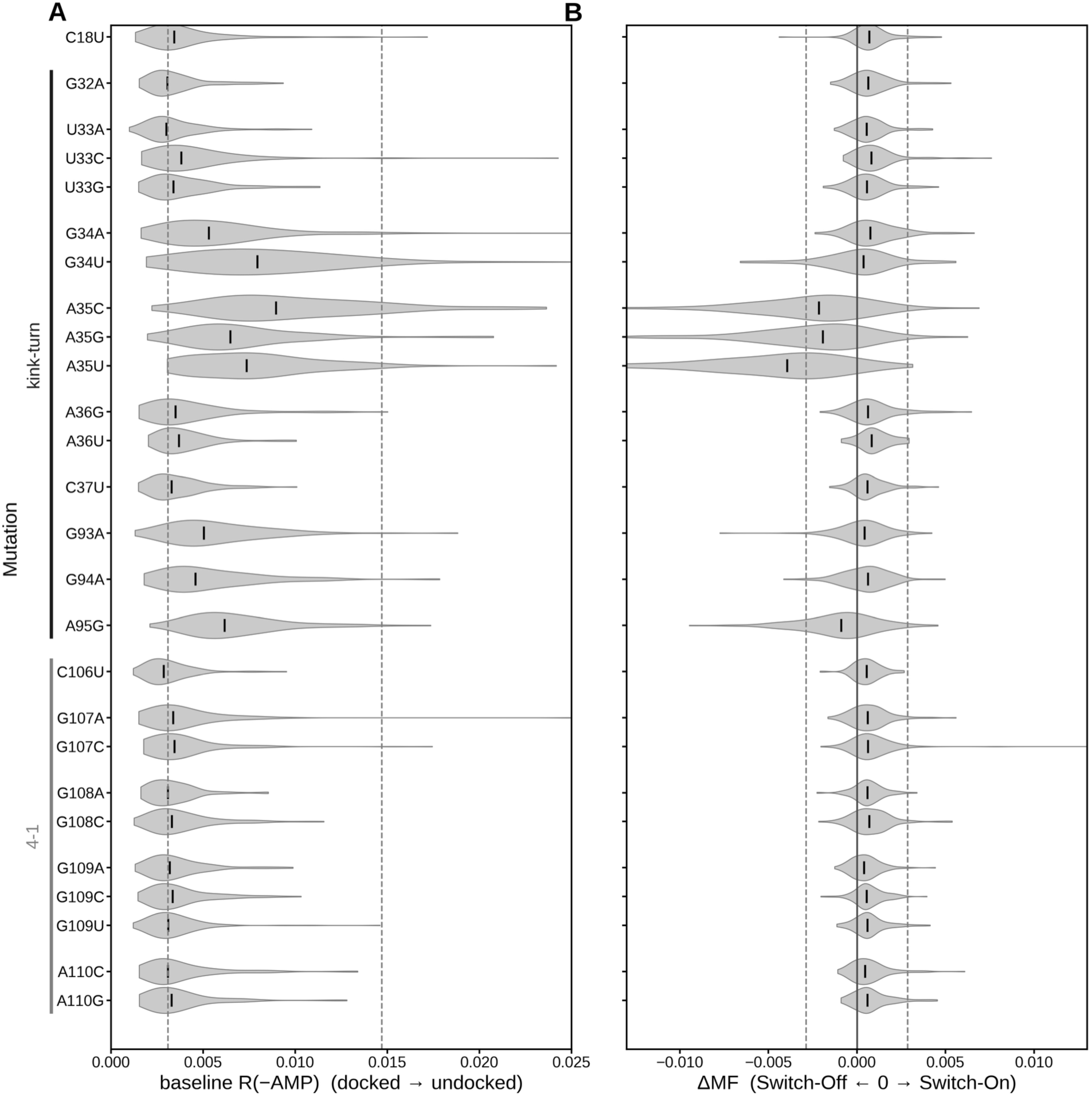
Junction single mutations shift the TL/TLR docking baseline and the AMP response by position. For each single junction mutation (rows, ordered along the construct and grouped by junction, kink-turn versus 4-1), the distribution across roughly 120 randomized helix backgrounds of (A) baseline TL/TLR reactivity without AMP, from docked (low) to undocked (high); the dashed lines are the θ_low and θ_high baseline cutoffs from Figure 2. (B) The AMP-induced change ΔMF(TL/TLR), where negative values are Switch-Off and positive values are Switch-On; the dashed lines are the ±3σ_helix cutoffs. The kink-turn 1b mutations A35C, A35G, and A35U shift the baseline toward undocked and push ΔMF negative, and A95G behaves the same way, while the 4-1 junction mutations stay docked with near-zero ΔMF.

**Figure S13:**
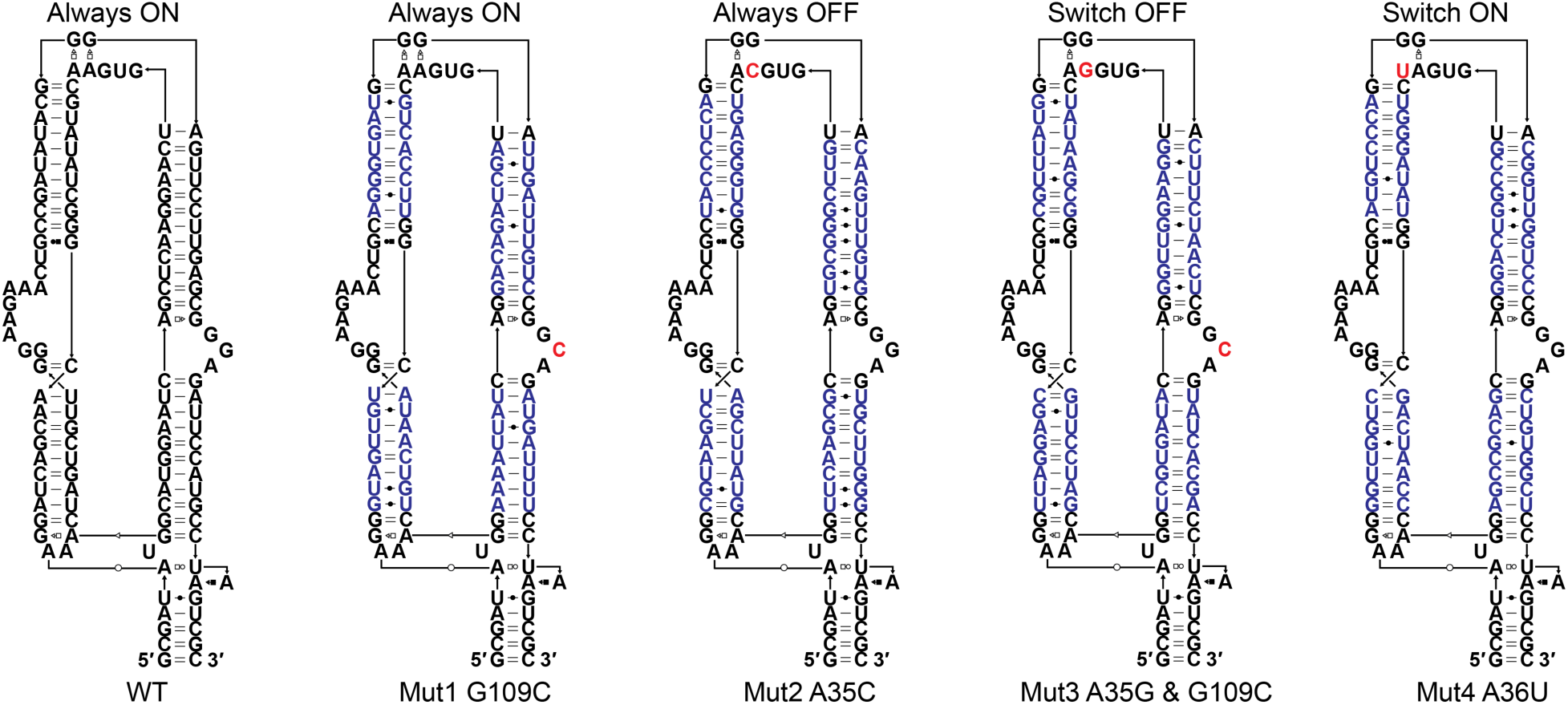
Secondary structures of the wild-type and the four representative mutants. Secondary-structure diagrams of the wild-type construct and the four phenotype representatives, labeled by allosteric regime (top) and by construct and mutation (bottom): wild-type (Always-On), Mut1 G109C (4-1 junction, Always-On), Mut2 A35C (kink-turn 1b, Always-Off), Mut3 A35G and G109C (kink-turn 1b plus 4-1, Switch-Off), and Mut4 A36U (kink-turn 2b, Switch-On). The mutated nucleotide is highlighted in red. Blue marks the helix-background positions that are randomized across library variants, drawn here as each representative’s actual library sequence, while the wild-type reference is shown without randomization; black marks the fixed aptamer, junction, and TL/TLR positions.

**Figure S14:**
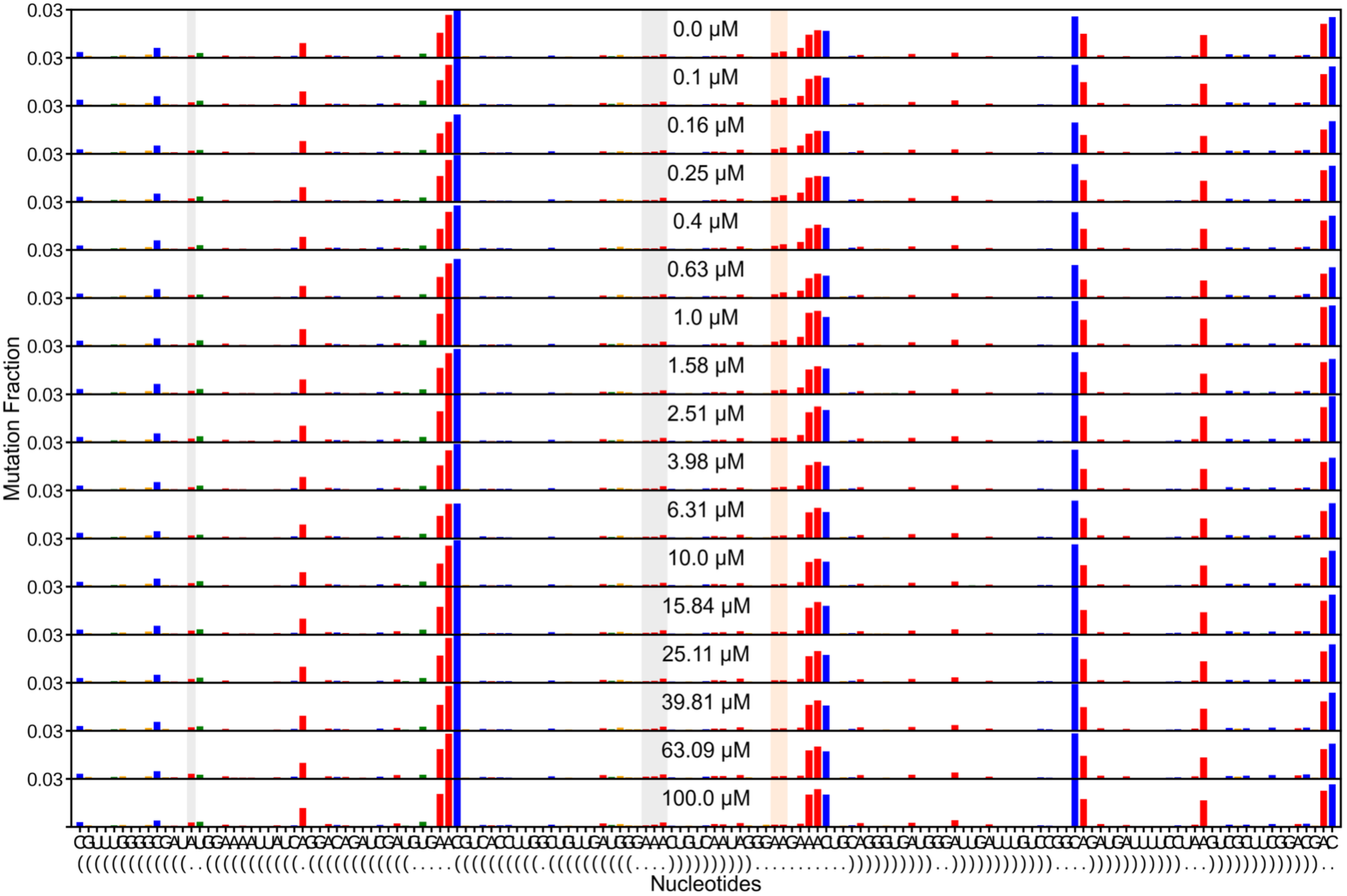
Always-On per-nucleotide reactivity across the AMP titration, replicate 1. Full per-nucleotide DMS-MaPseq reactivity for the Always-On representative (C01EI) at each point of the AMP titration (replicate 1), stacked by AMP concentration. Gray shading marks the TL/TLR docking reporters and orange shading the ATP-aptamer binding-site adenines. The aptamer adenines lose reactivity as AMP rises, so the aptamer binds AMP, while the TL/TLR reporters stay low across the titration, so the tertiary contact is docked regardless of ligand. These are the raw per-replicate data behind the Always-On curves in Figure 3B.

**Figure S15:**
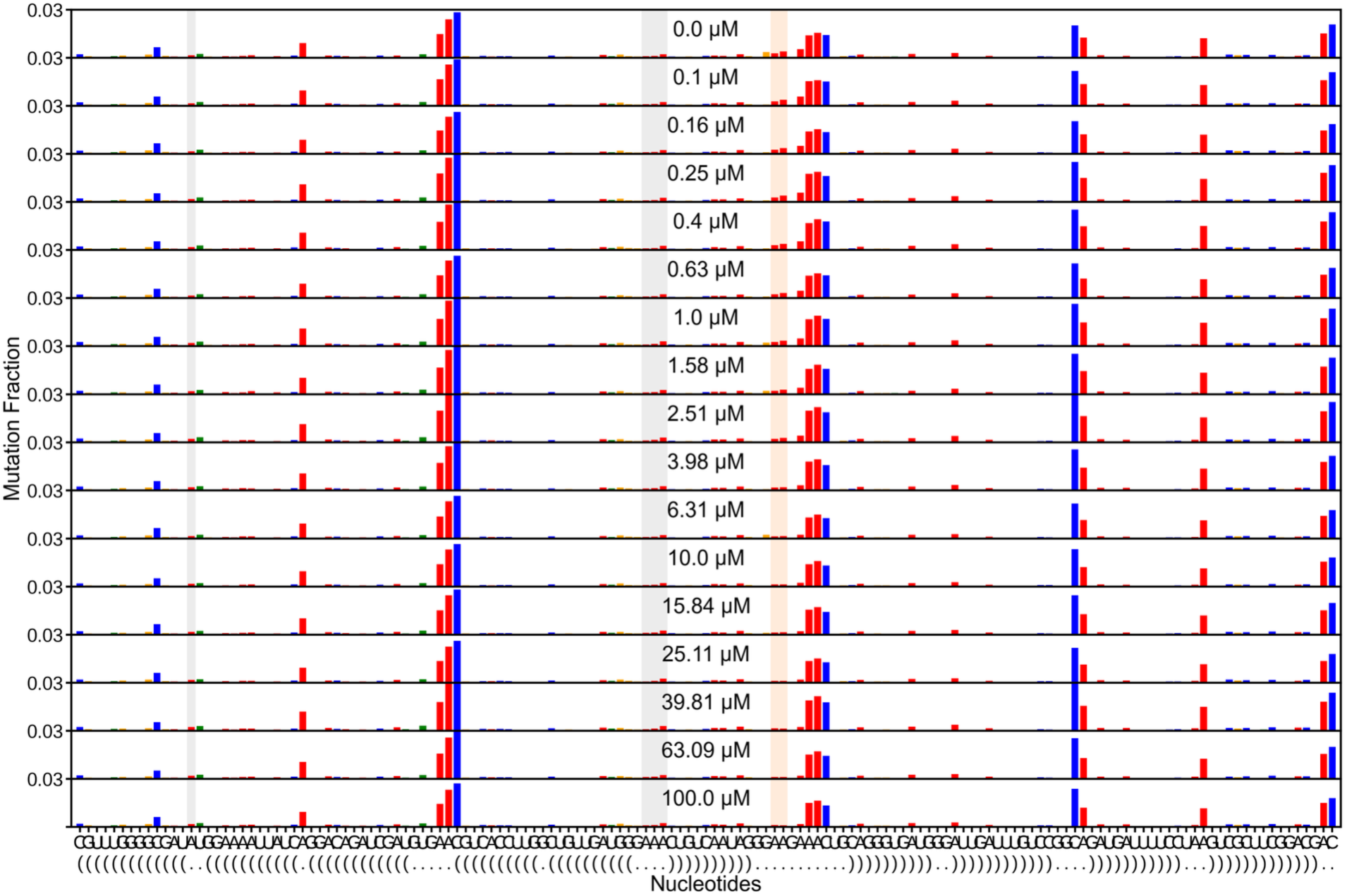
Always-On per-nucleotide reactivity across the AMP titration, replicate 2. As in Figure S13 for the second Always-On replicate. The aptamer adenines (orange) are protected as AMP increases while the TL/TLR reporters (gray) stay low, reproducing the docked, ligand-insensitive behavior of the Always-On regime.

**Figure S16:**
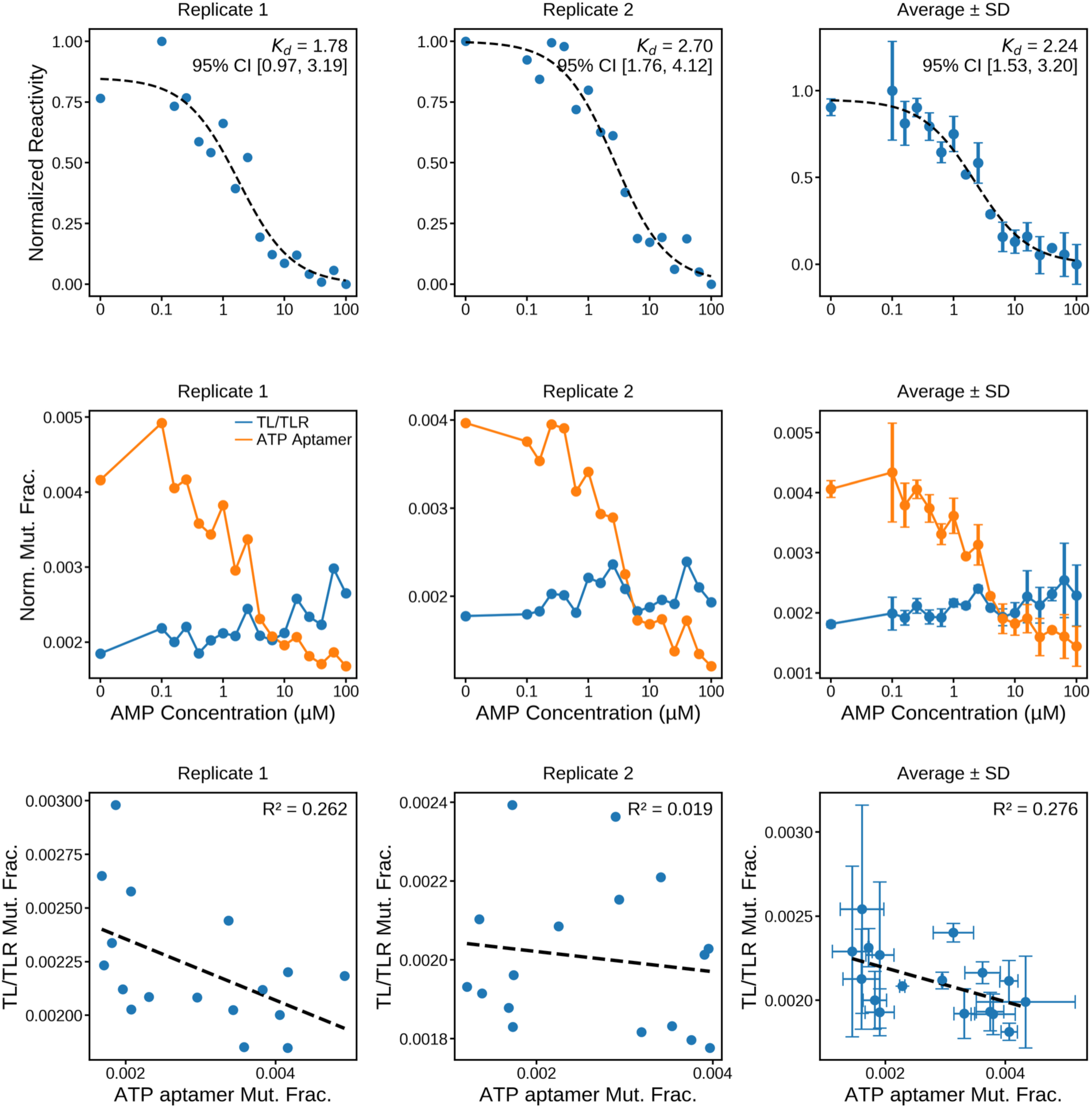
Always-On AMP binding affinity and weak coupling to TL/TLR docking. (Top) Mean reactivity of the AMP-sensitive ATP-aptamer binding-site adenines, normalized to [0, 1], versus AMP concentration for each Always-On replicate and for the inter-replicate average (± SD). Dashed lines are three-parameter Hill fits, with the apparent Kd annotated. The replicate fits give Kd = 1.78 ± 0.66 and 2.70 ± 0.60 μM, and averaging gives Kd = 2.24 ± 0.43 μM, so the Always-On representative binds AMP with wild-type-like affinity. (Middle) Per-nucleotide normalized mutation fraction across the same titration for the TL/TLR docking reporters (blue) and the ATP-aptamer binding-site adenines (orange). Aptamer reactivity falls with AMP while the TL/TLR reporters stay low, so the contact stays docked throughout. (Bottom) TL/TLR docking reactivity versus ATP-aptamer reactivity across the titration, per replicate and averaged, with the coefficient of determination annotated.

**Figure S17:**
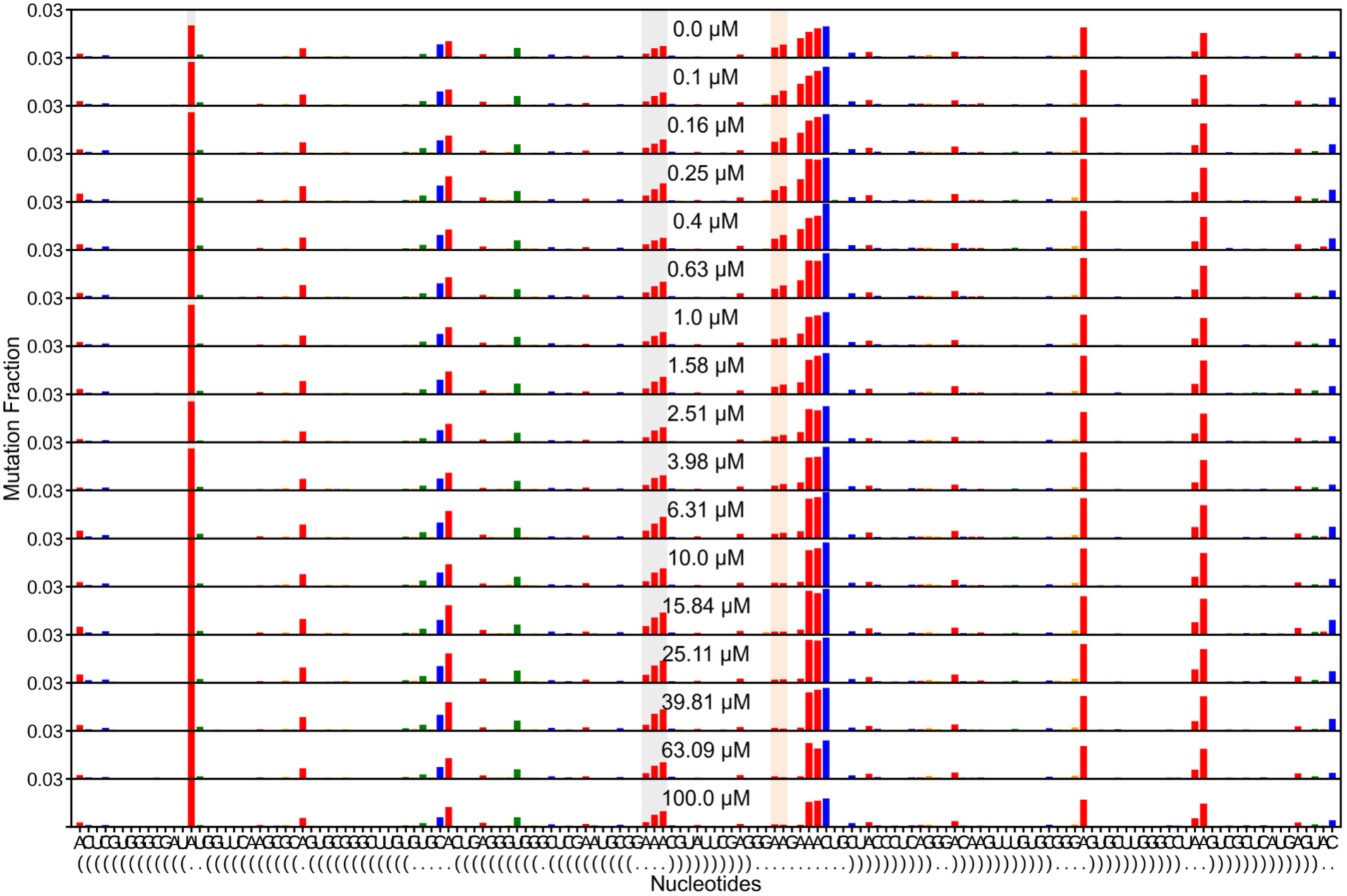
Always-Off per-nucleotide reactivity across the AMP titration, replicate 1. Full per-nucleotide DMS-MaPseq reactivity for the Always-Off representative (C01EK) at each point of the AMP titration (replicate 1), stacked from low AMP (top) to high AMP (bottom); each row is the mutation fraction at every nucleotide for one AMP concentration. Gray shading marks the TL/TLR docking reporters (the GAAA tetraloop adenines and the receptor adenine) and orange shading the ATP-aptamer binding-site adenines. The aptamer adenines lose reactivity as AMP rises, so the aptamer binds AMP, while the TL/TLR reporters stay high across the titration, so the tertiary contact is undocked regardless of ligand. These are the raw per-replicate data behind the Always-Off curves in Figure 3C.

**Figure S18:**
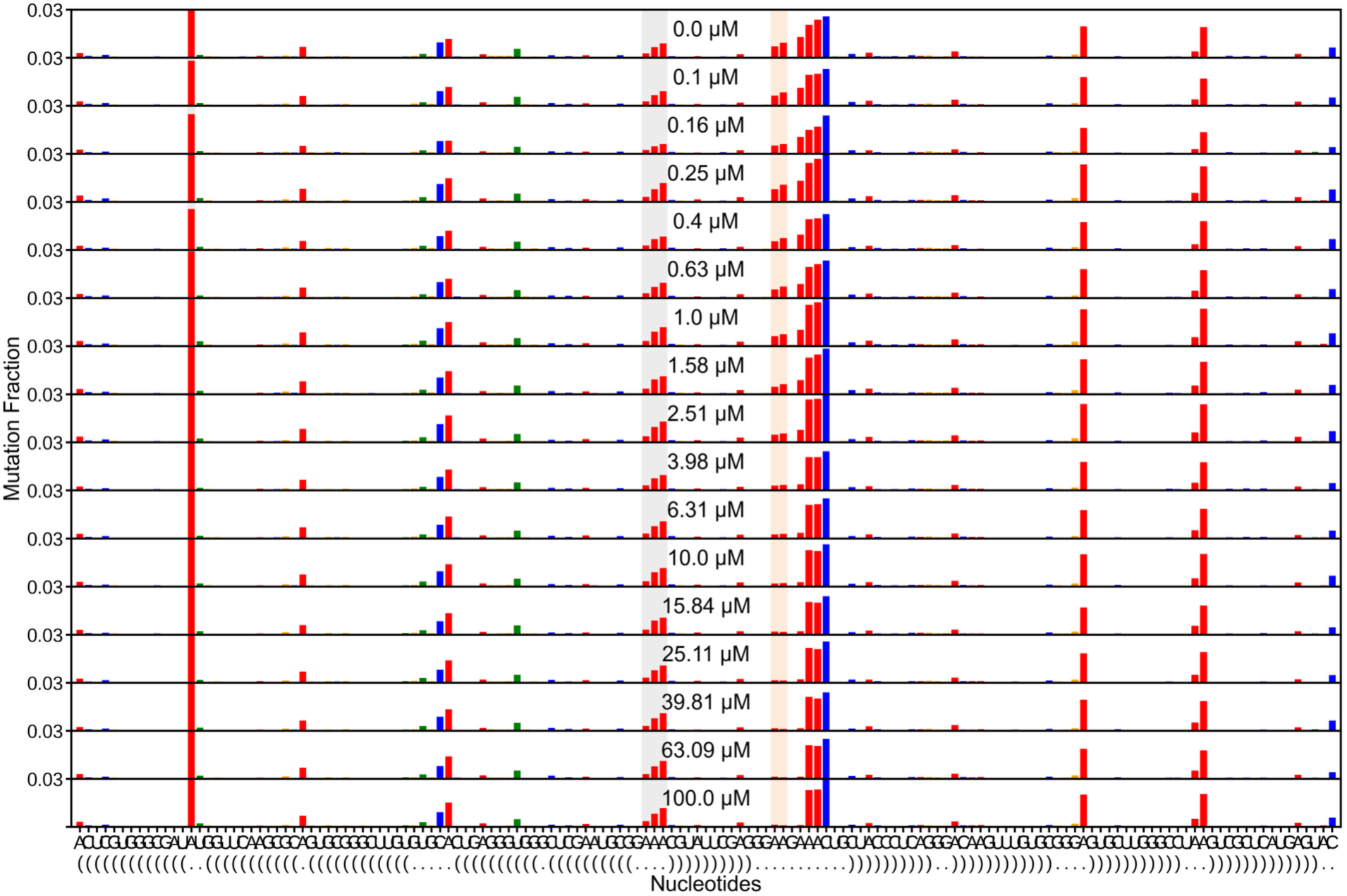
Always-Off per-nucleotide reactivity across the AMP titration, replicate 2. As in Figure S16 for the second Always-Off replicate. The aptamer adenines (orange) are protected as AMP increases while the TL/TLR reporters (gray) stay high, reproducing the undocked, ligand-insensitive docking of the Always-Off regime.

**Figure S19:**
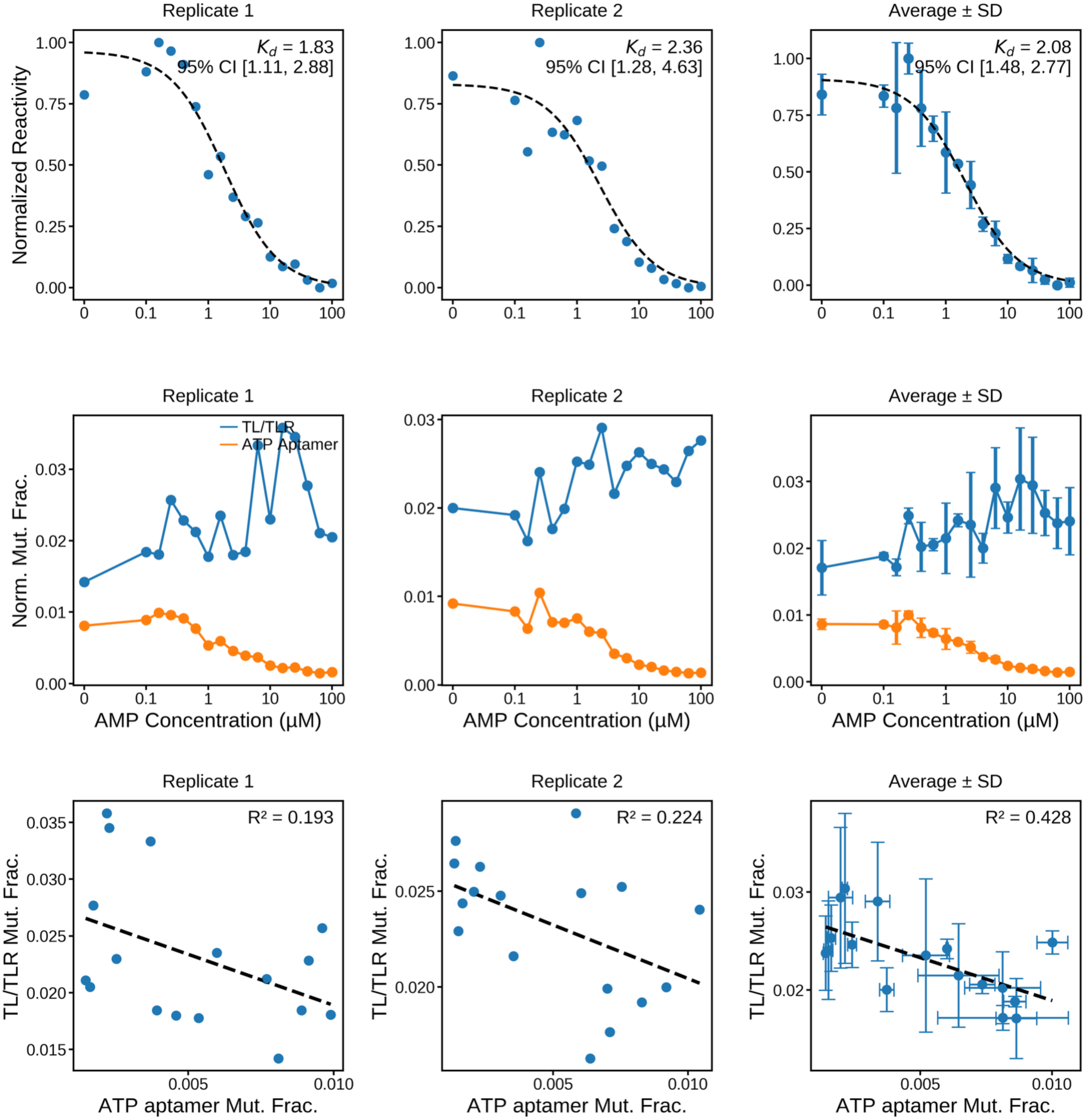
Always-Off AMP binding affinity and weak negative coupling to TL/TLR docking. (Top) Mean reactivity of the AMP-sensitive ATP-aptamer binding-site adenines, normalized to [0, 1], versus AMP concentration for each Always-Off replicate and for the inter-replicate average (± SD). Dashed lines are three-parameter Hill fits, with the apparent Kd annotated. The replicate fits give Kd = 1.83 ± 0.49 and 2.36 ± 0.76 μM, and averaging the replicates gives Kd = 2.08 ± 0.33 μM, so the Always-Off representative binds AMP with wild-type-like affinity. (Middle) Per-nucleotide normalized mutation fraction across the same titration for the TL/TLR docking reporters (blue) and the ATP-aptamer binding-site adenines (orange). Aptamer reactivity falls with AMP, while the TL/TLR reporters stay high and rise modestly, so the contact is undocked with or without ligand. (Bottom) TL/TLR docking reactivity versus ATP-aptamer reactivity across the titration, per replicate and averaged, with the coefficient of determination annotated. The average anticorrelation (R² = 0.428) reflects the small AMP-induced rise in TL/TLR reactivity, the weak negative coupling quantified in Figure 3C.

**Figure S20:**
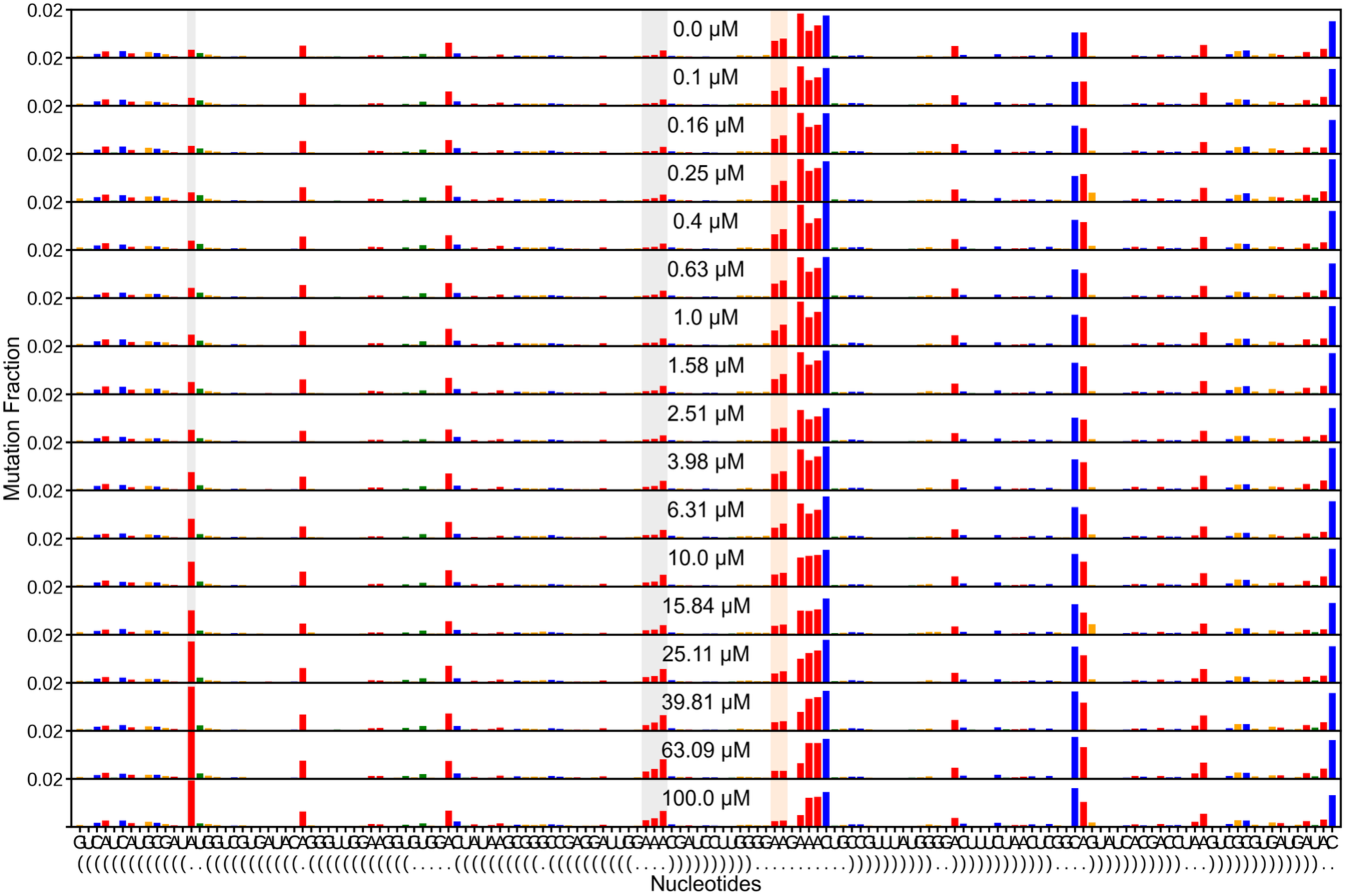
Switch-Off per-nucleotide reactivity across the AMP titration, replicate 1. Full per-nucleotide DMS-MaPseq reactivity for the Switch-Off representative (C01EJ) at each point of the AMP titration (replicate 1), stacked by AMP concentration. Gray shading marks the TL/TLR docking reporters and orange shading the ATP-aptamer binding-site adenines. The aptamer adenines lose reactivity as AMP rises, so the aptamer binds AMP, and the TL/TLR reporters gain reactivity over the same range, so AMP binding undocks the tertiary contact. These are the raw per-replicate data behind the Switch-Off curves in Figure 3D.

**Figure S21:**
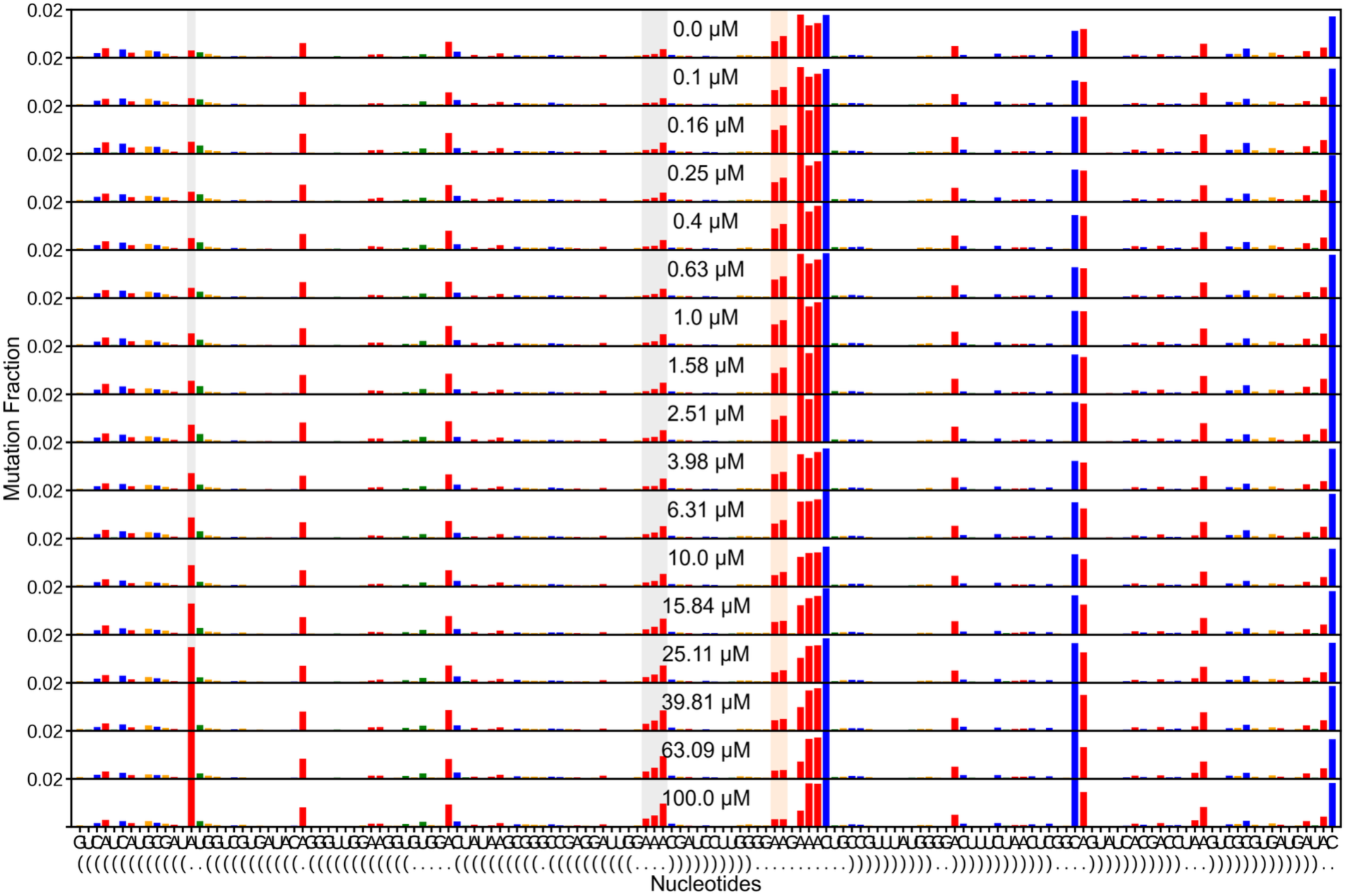
Switch-Off per-nucleotide reactivity across the AMP titration, replicate 2. As in Figure S19 for the second Switch-Off replicate. The aptamer adenines (orange) are protected while the TL/TLR reporters (gray) gain reactivity as AMP increases, reproducing the AMP-induced loss of docking that defines the Switch-Off regime.

**Figure S22:**
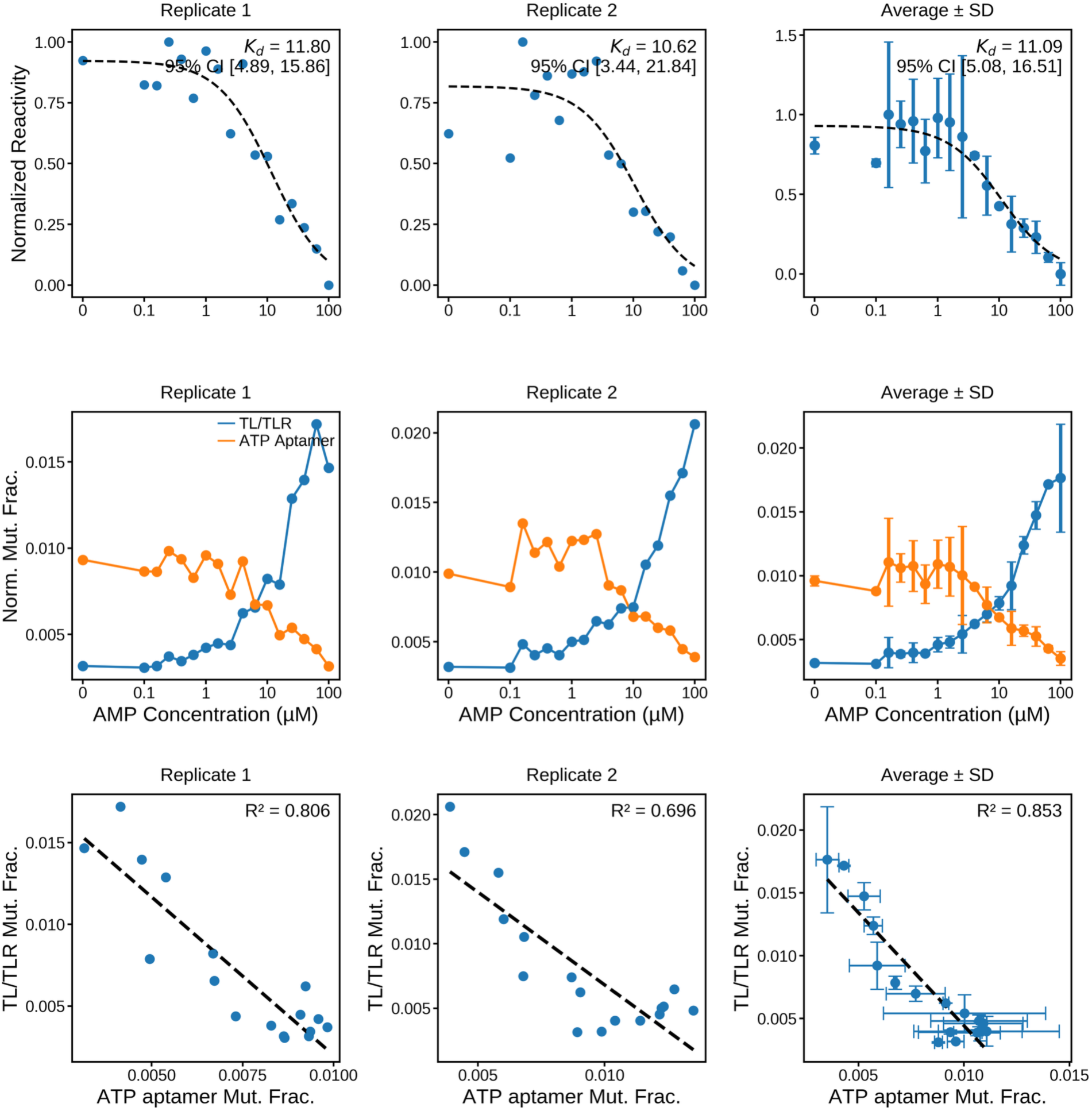
Switch-Off AMP binding drives loss of TL/TLR docking. (Top) Mean reactivity of the AMP-sensitive ATP-aptamer binding-site adenines, normalized to [0, 1], versus AMP concentration for each Switch-Off replicate and for the inter-replicate average (± SD). Dashed lines are three-parameter Hill fits, with the apparent Kd annotated. The replicate fits give Kd = 11.80 ± 2.98 and 10.62 ± 4.68 μM, and averaging gives Kd = 11.09 ± 2.99 μM. (Middle) Per-nucleotide normalized mutation fraction across the same titration for the TL/TLR docking reporters (blue) and the ATP-aptamer binding-site adenines (orange). The two arms move in opposite directions: aptamer reactivity falls as TL/TLR reactivity rises, so AMP binding undocks the contact. (Bottom) TL/TLR docking reactivity versus ATP-aptamer reactivity across the titration, per replicate and averaged, with the coefficient of determination annotated. The strong average anticorrelation (R² = 0.853) reflects the tight AMP-coupled undocking, the negative coupling quantified in Figure 3D.

**Figure S23:**
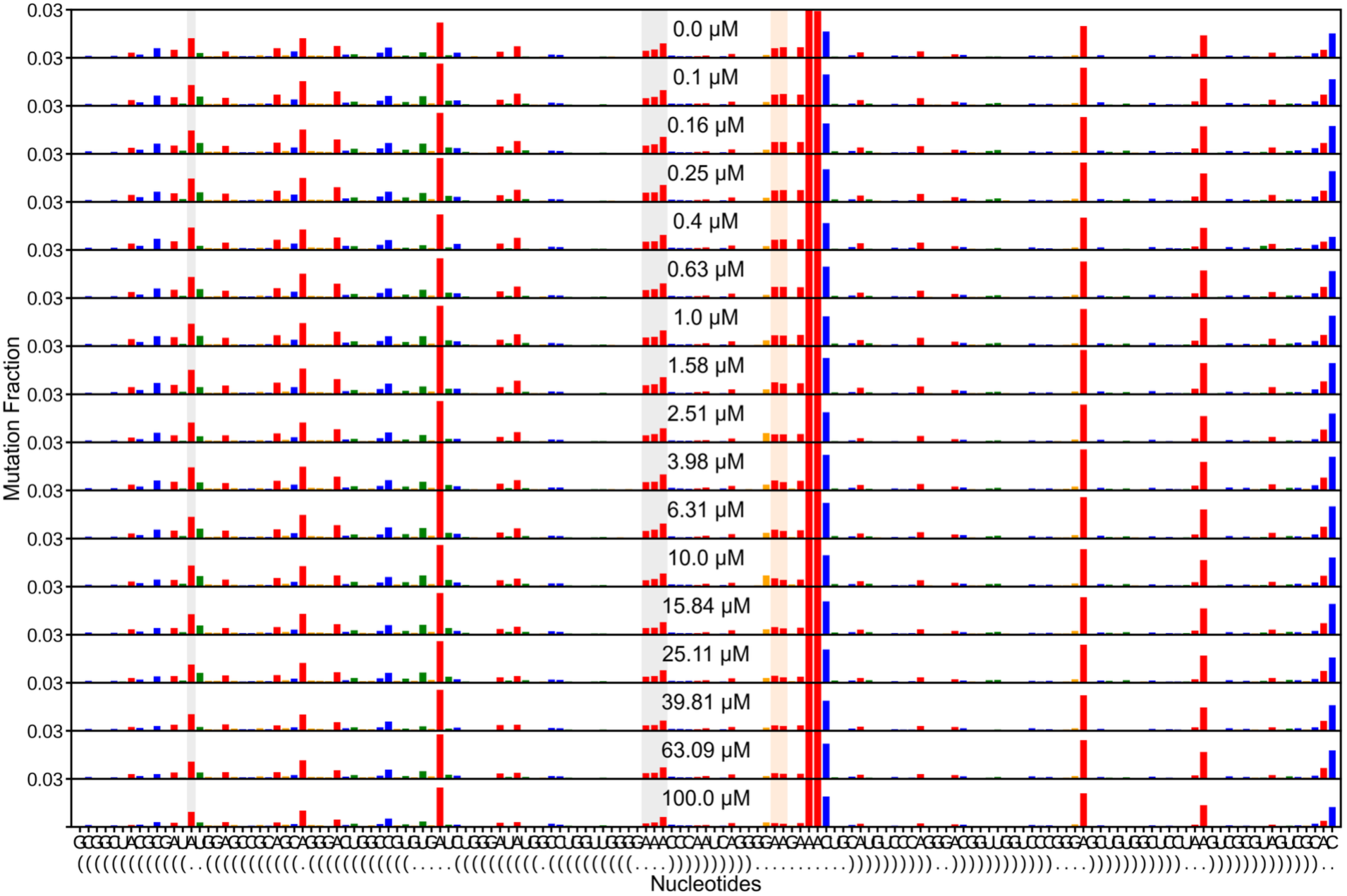
Switch-On per-nucleotide reactivity across the AMP titration at 22 °C, replicate 1. Full per-nucleotide DMS-MaPseq reactivity for the Switch-On representative (C01FL) at each point of the AMP titration (replicate 1), measured at 22 °C and stacked by AMP concentration. Gray shading marks the TL/TLR docking reporters and orange shading the ATP-aptamer binding-site adenines. The aptamer adenines lose reactivity as AMP rises, so AMP still binds at 22 °C, but the TL/TLR docking response is weaker and less consistent than at 44 °C (Figure S22). These are the raw per-replicate data behind the 22 °C fit in Figure S29.

**Figure S24:**
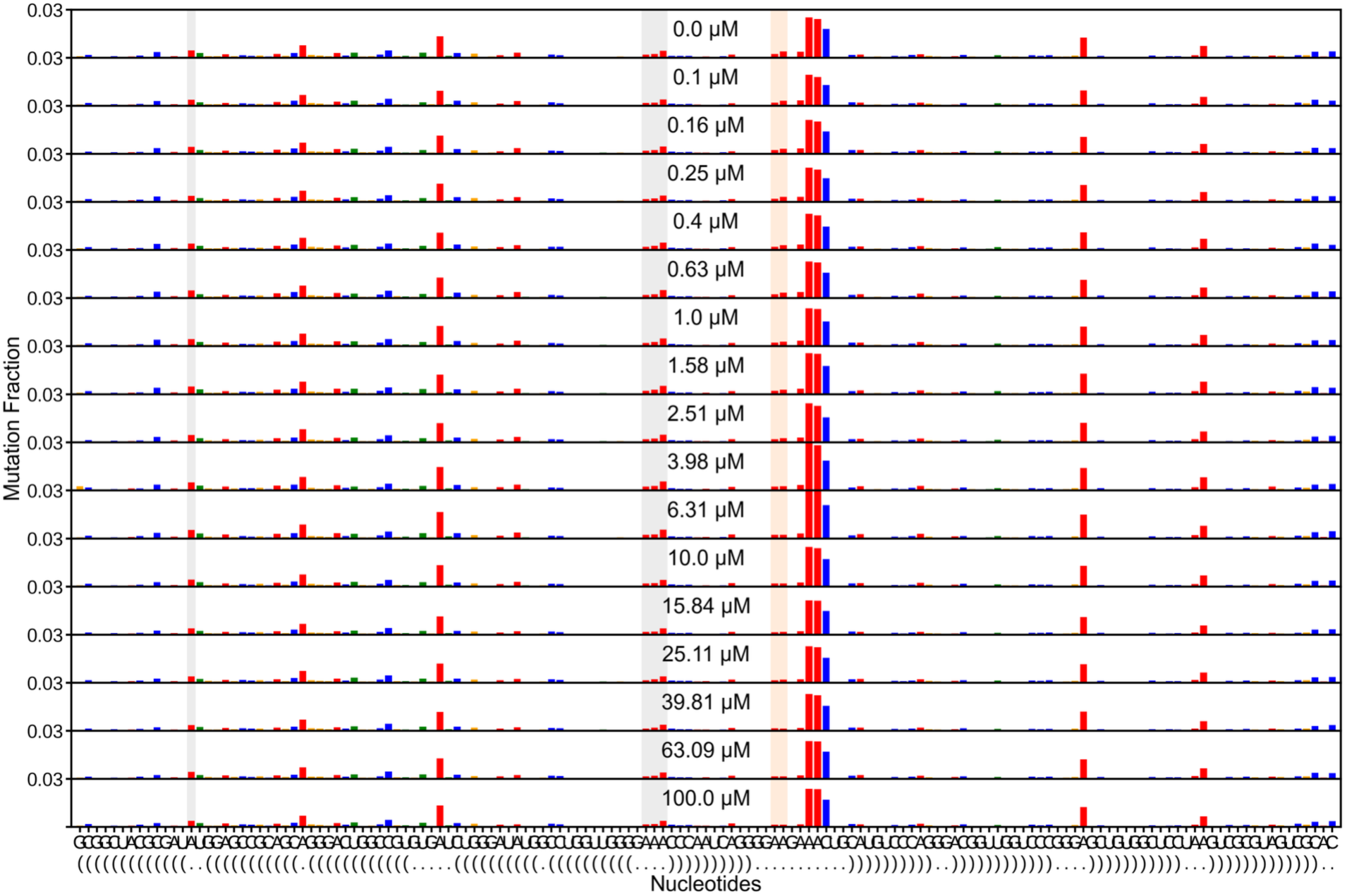
Switch-On per-nucleotide reactivity across the AMP titration at 22 °C, replicate 2. As in Figure S26 for the second 22 °C replicate. AMP protects the aptamer adenines (orange), but the TL/TLR reporters (blue) track binding only weakly, the replicate-variable docking response that motivated measuring Switch-On at 44 °C.

**Figure S25:**
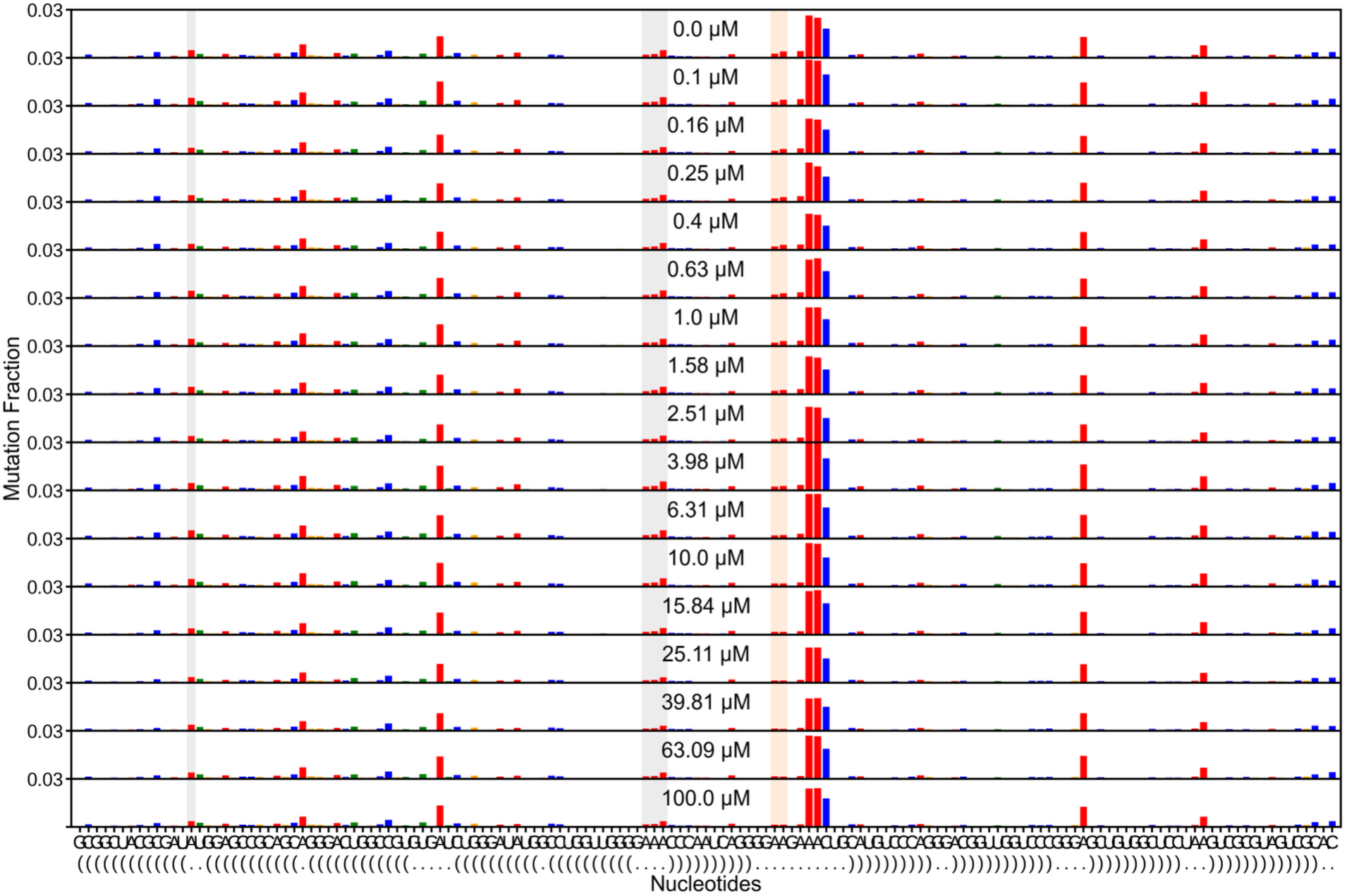
Switch-On per-nucleotide reactivity across the AMP titration at 22 °C, replicate 3. As in Figure S26 for the third 22 °C replicate. AMP again protects the aptamer adenines while the TL/TLR docking signal stays small, consistent with the attenuated 22 °C coupling.

**Figure S26:**
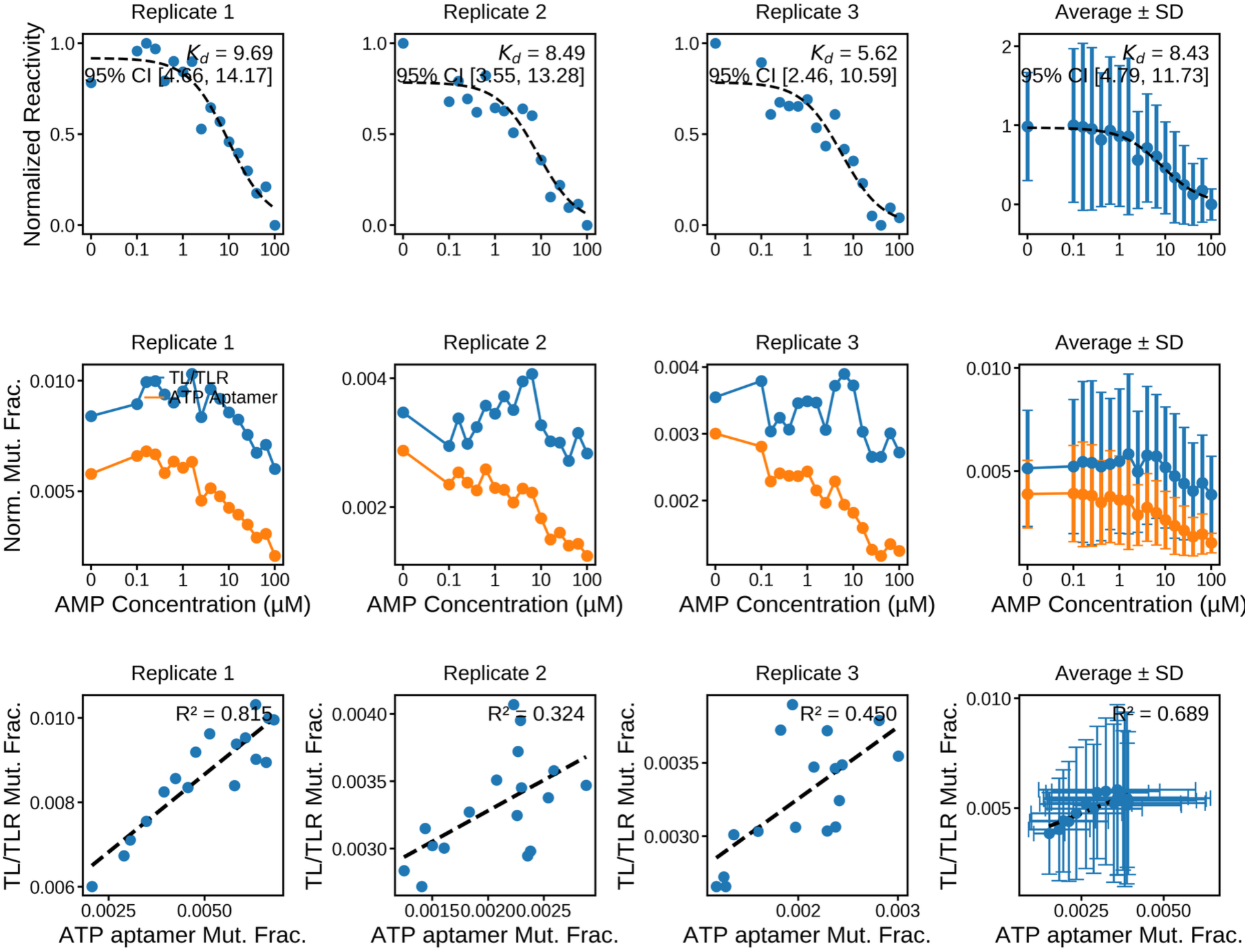
Switch-On AMP binding at 22 °C with attenuated TL/TLR docking. Measured at 22 °C over the same AMP range as the 44 °C panel (Figure S25). (Top) Mean reactivity of the AMP-sensitive ATP-aptamer binding-site adenines, normalized to [0, 1], versus AMP concentration for each replicate and the inter-replicate average (± SD). Dashed lines are three-parameter Hill fits, with the apparent Kd annotated. The replicate fits give Kd = 9.69 ± 2.40, 8.49 ± 2.78, and 5.62 ± 1.93 μM, and averaging gives Kd = 8.43 ± 1.57 μM, so the aptamer binds AMP at 22 °C. (Middle) Per-nucleotide normalized mutation fraction across the same titration for the TL/TLR docking reporters (blue) and the ATP-aptamer binding-site adenines (orange). The aptamer arm falls with AMP while the TL/TLR arm responds only weakly. (Bottom) TL/TLR docking reactivity versus ATP-aptamer reactivity across the titration, per replicate and averaged, with the coefficient of determination annotated. The correlation is weak and replicate-variable (R² = 0.815, 0.324, and 0.450; average 0.689), far below the 44 °C value (R² = 0.943, Figure S25). AMP binds at 22 °C but drives little docking; the same coupling amplifies into a reliable signal at 44 °C.

**Figure S27:**
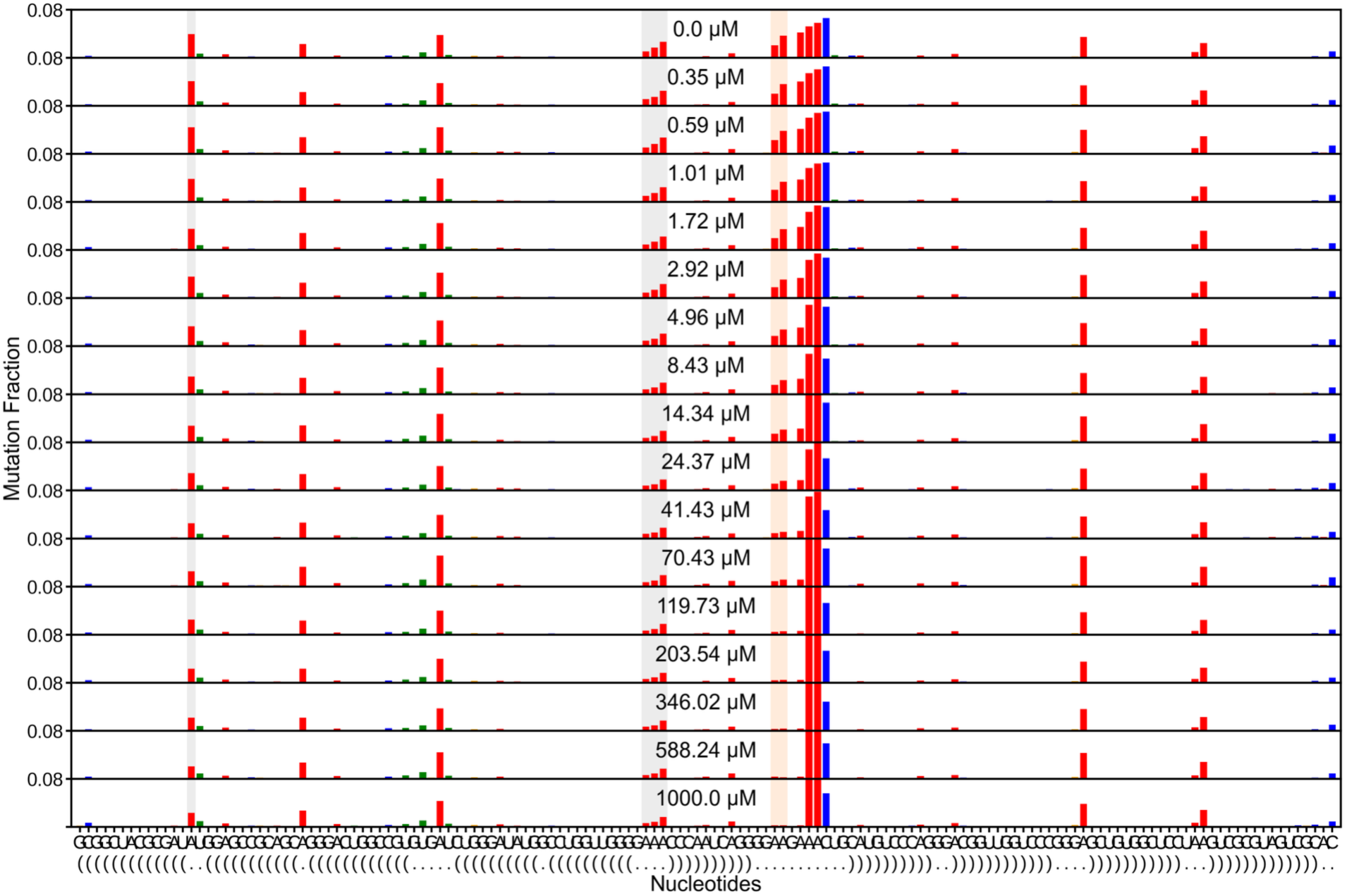
Switch-On per-nucleotide reactivity across the AMP titration, replicate 1. Full per-nucleotide DMS-MaPseq reactivity for the Switch-On representative (C01FL) at each point of the AMP titration (replicate 1), measured at 44 °C over a wider AMP range (to 1000 μM) and stacked by AMP concentration. Gray shading marks the TL/TLR docking reporters and orange shading the ATP-aptamer binding-site adenines. Both sets of reporters lose reactivity as AMP rises: the aptamer adenines report binding and the TL/TLR reporters report docking, so AMP binding drives the tertiary contact to dock. These are the raw per-replicate data behind the Switch-On curves in Figure 3E.

**Figure S28:**
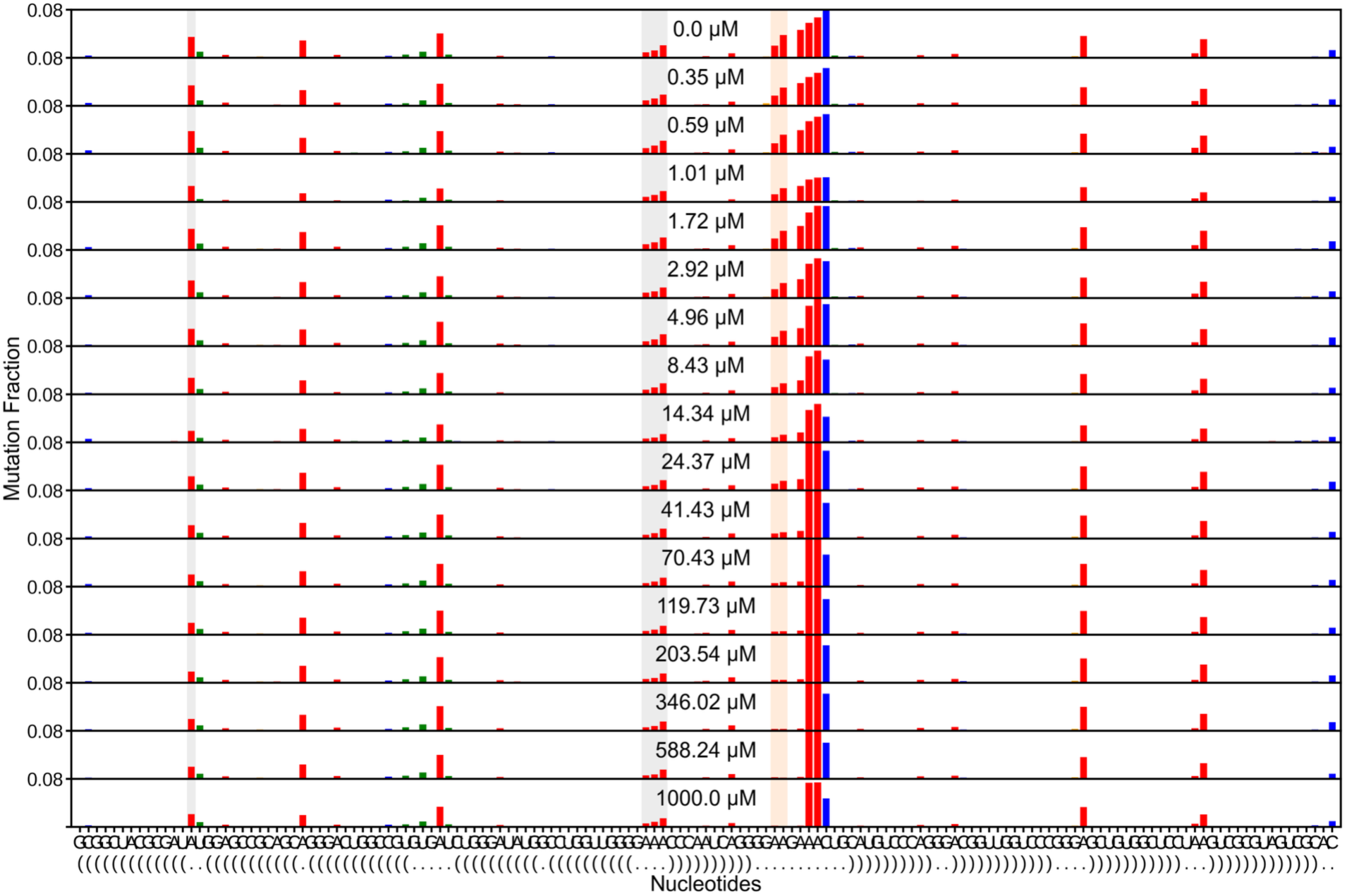
Switch-On per-nucleotide reactivity across the AMP titration, replicate 2. As in Figure S22 for the second Switch-On replicate. Both the aptamer (orange) and TL/TLR (gray) reporters are protected as AMP increases, reproducing the AMP-induced gain of docking that defines the Switch-On regime.

**Figure S29:**
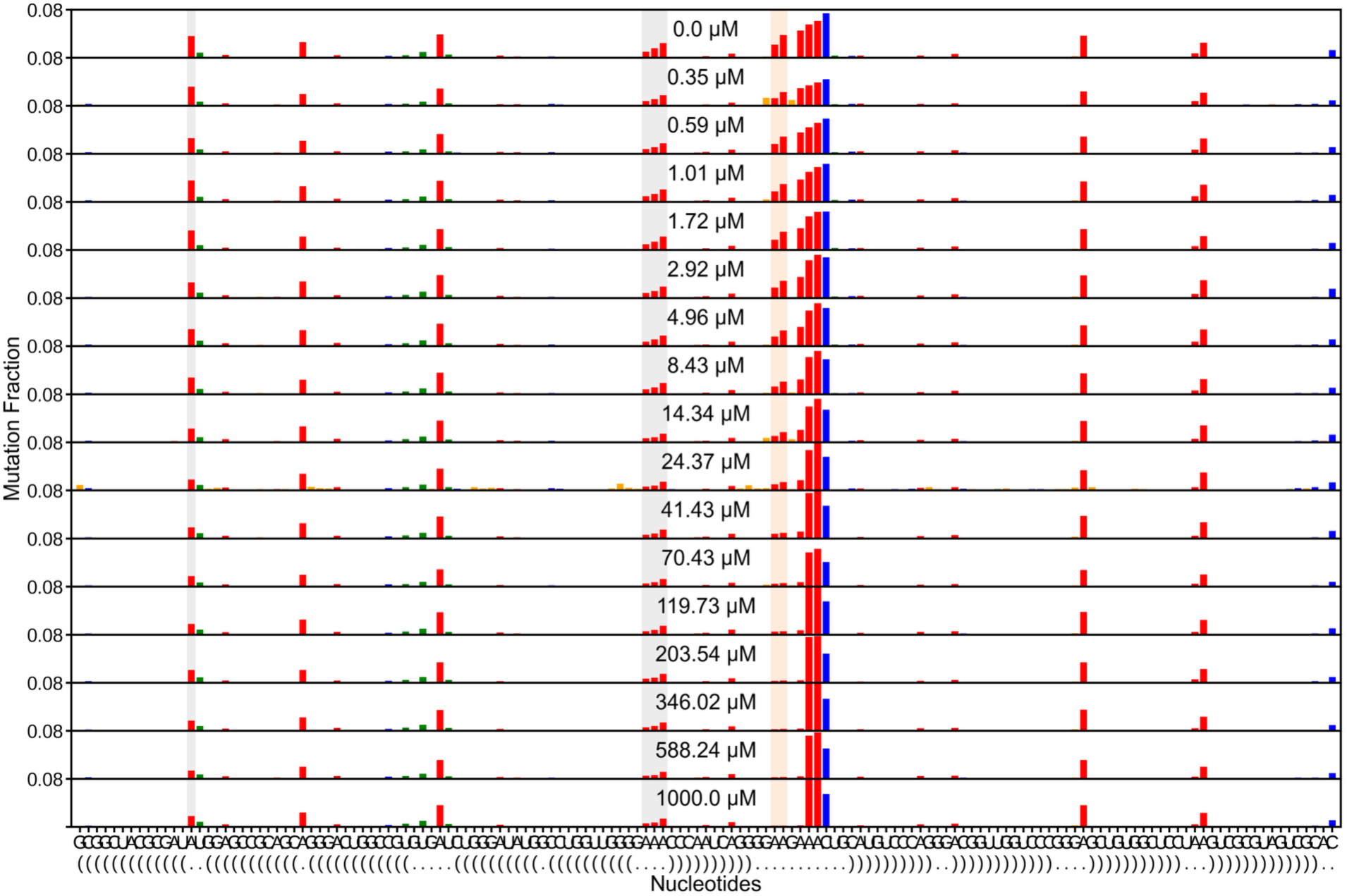
Switch-On per-nucleotide reactivity across the AMP titration, replicate 3. As in Figure S22 for the third Switch-On replicate. The aptamer and TL/TLR reporters lose reactivity together as AMP increases, consistent across all three replicates.

**Figure S30:**
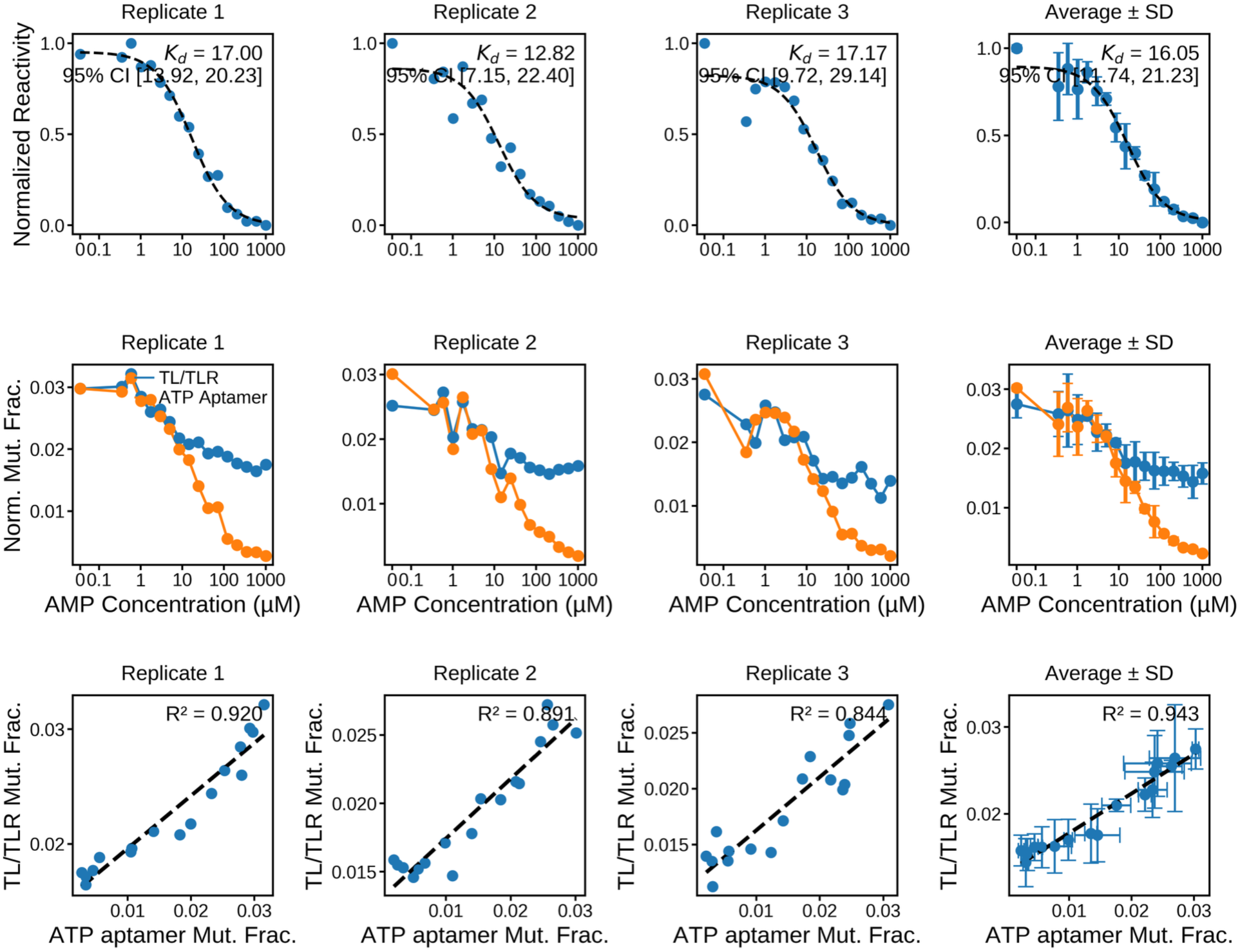
Switch-On AMP binding drives TL/TLR docking. Measured at 44 °C over a wider AMP range (to 1000 μM). (Top) Mean reactivity of the AMP-sensitive ATP-aptamer binding-site adenines, normalized to [0, 1], versus AMP concentration for each Switch-On replicate and for the inter-replicate average (± SD). Dashed lines are three-parameter Hill fits, with the apparent Kd annotated. The replicate fits give Kd = 17.00 ± 1.74, 12.82 ± 3.80, and 17.17 ± 5.01 μM, and averaging gives Kd = 16.05 ± 2.54 μM. (Middle) Per-nucleotide normalized mutation fraction across the same titration for the TL/TLR docking reporters (blue) and the ATP-aptamer binding-site adenines (orange). Both arms fall together as AMP rises, so AMP binding and TL/TLR docking track each other. (Bottom) TL/TLR docking reactivity versus ATP-aptamer reactivity across the titration, per replicate and averaged, with the coefficient of determination annotated. The strong average correlation (R² = 0.943) reflects the tight AMP-coupled docking, the positive coupling quantified in Figure 3E.

**Figure S31:**
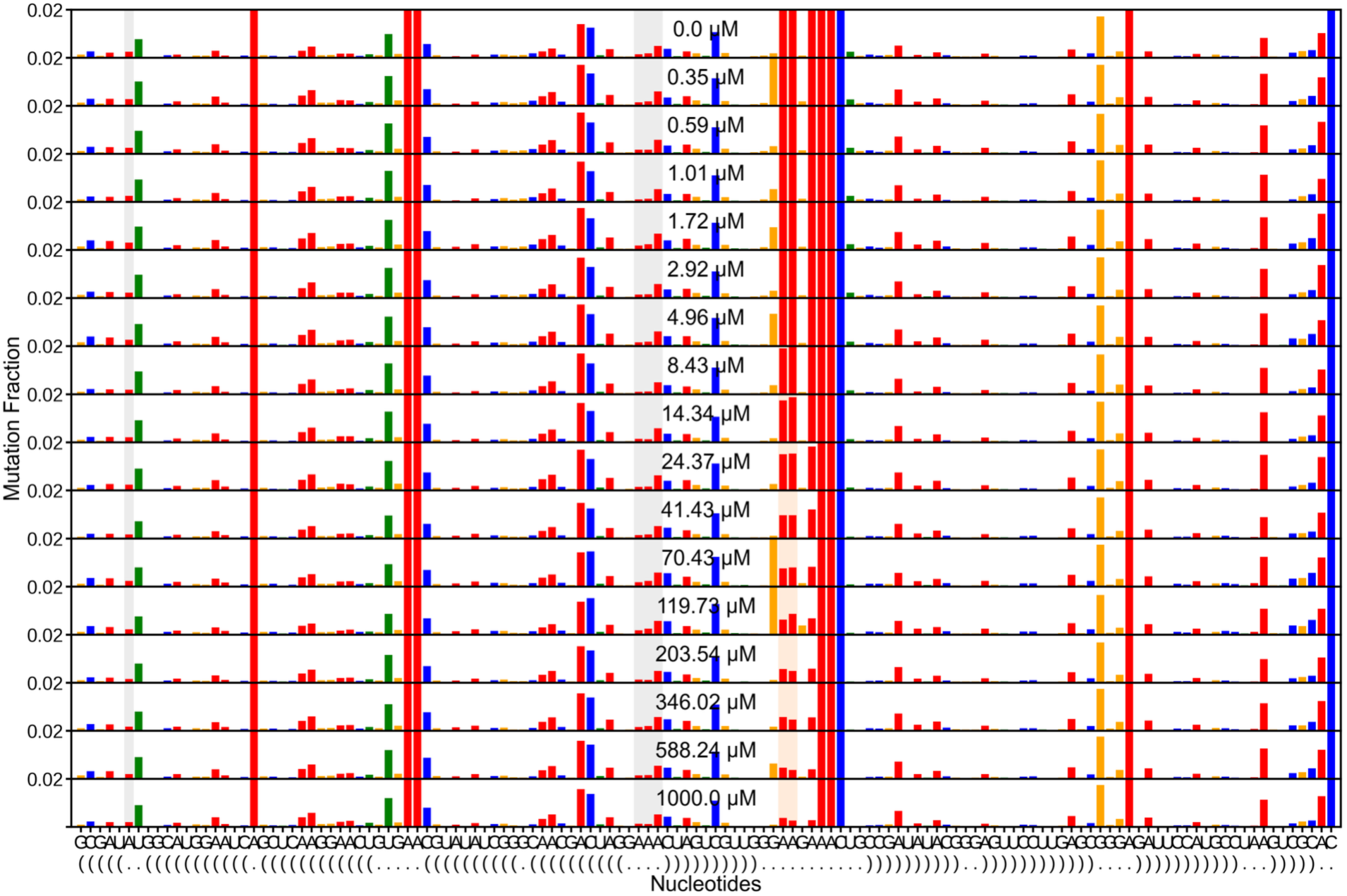
WT AMP titration at 44 °C, full per-nucleotide reactivity. Stacked population-average per-nucleotide reactivity across the AMP titration for the wild type at 44 °C (run 67, single replicate; n = 1). The y-axis is fixed to 0.02, the scale used for the 22 °C version (Figure S1), so the two temperatures compare directly. As at 22 °C, the TL/TLR reporters stay low across the titration, so the wild type remains docked at 44 °C.

**Figure S32:**
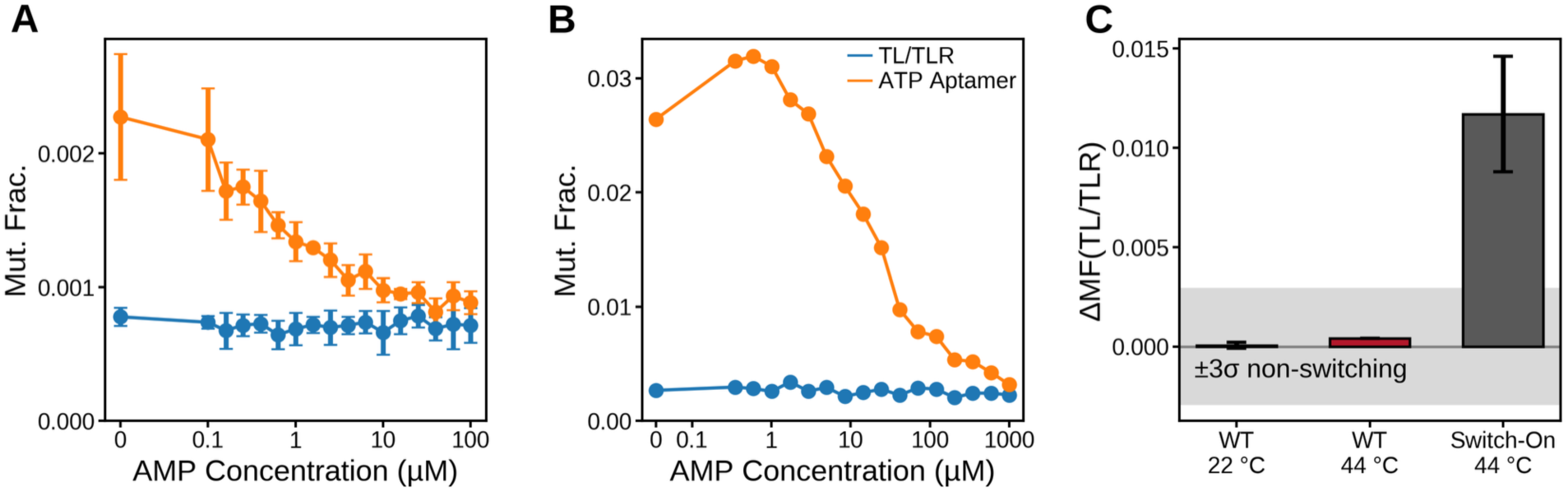
The wild type holds its regime at 44 °C. The Switch-On representative was measured at 44 °C, so this figure tests whether 44 °C by itself changes the wild-type regime. Dual AMP titration at (A) 22 °C (mean ± SD over replicates 2 to 4) and (B) 44 °C (single replicate from run 67; n = 1, no error bars), with the TL/TLR docking reporter (blue) and the ATP-aptamer reporter (orange). At both temperatures, the aptamer arm falls as AMP binds, while the TL/TLR arm stays flat. (C) ΔMF(TL/TLR), the TL/TLR reactivity without AMP minus that at saturating AMP, the same quantity used to classify the screen, against the ±3σ non-switching band (σ from the helix-only null distribution). Absolute DMS reactivity rises with temperature, so the regime is judged on this difference rather than on raw level. The wild type falls inside the band at both temperatures, while the Switch-On representative at 44 °C sits four times outside it.

**Figure S33:**
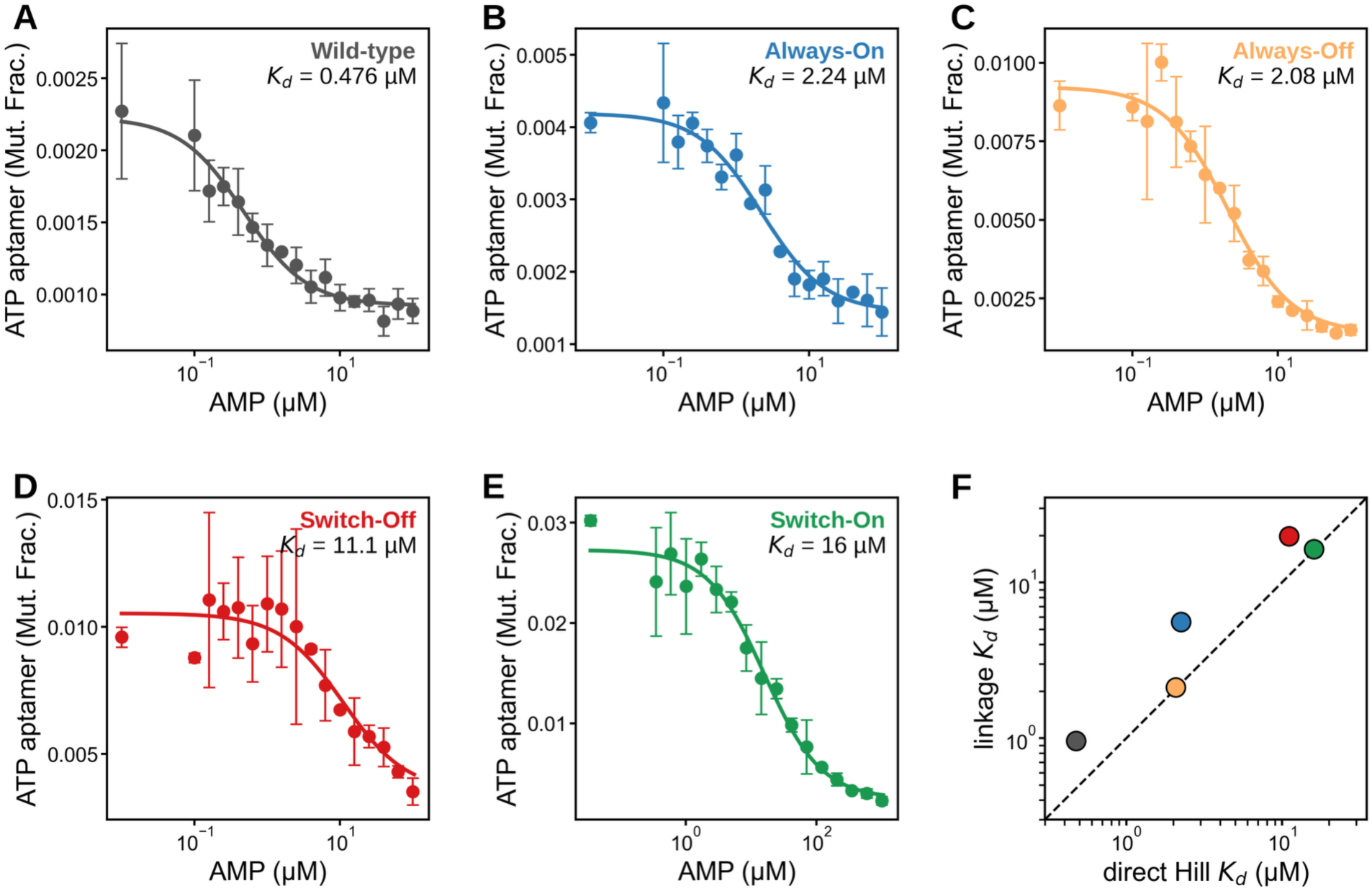
AMP affinity from direct Hill fits and Kd comparison. (A to E) The ATP-aptamer AMP titration for the wild type and the four representatives, each fit to a three-parameter Hill equation with the apparent K_d_ annotated, plotted on a log AMP axis in the class colors of Figure 4. Switch-On is measured at 44 °C. (F) Apparent AMP Kd from the direct Hill fit (x-axis) versus the four-state linkage-model K_d_ (y-axis). The two agree where the TL/TLR arm responds to AMP and diverge for the flat-arm constructs (wild type, Always-On), where the linkage K_d_ is underdetermined; the direct Hill value is reported for those cases.

**Figure S34:**
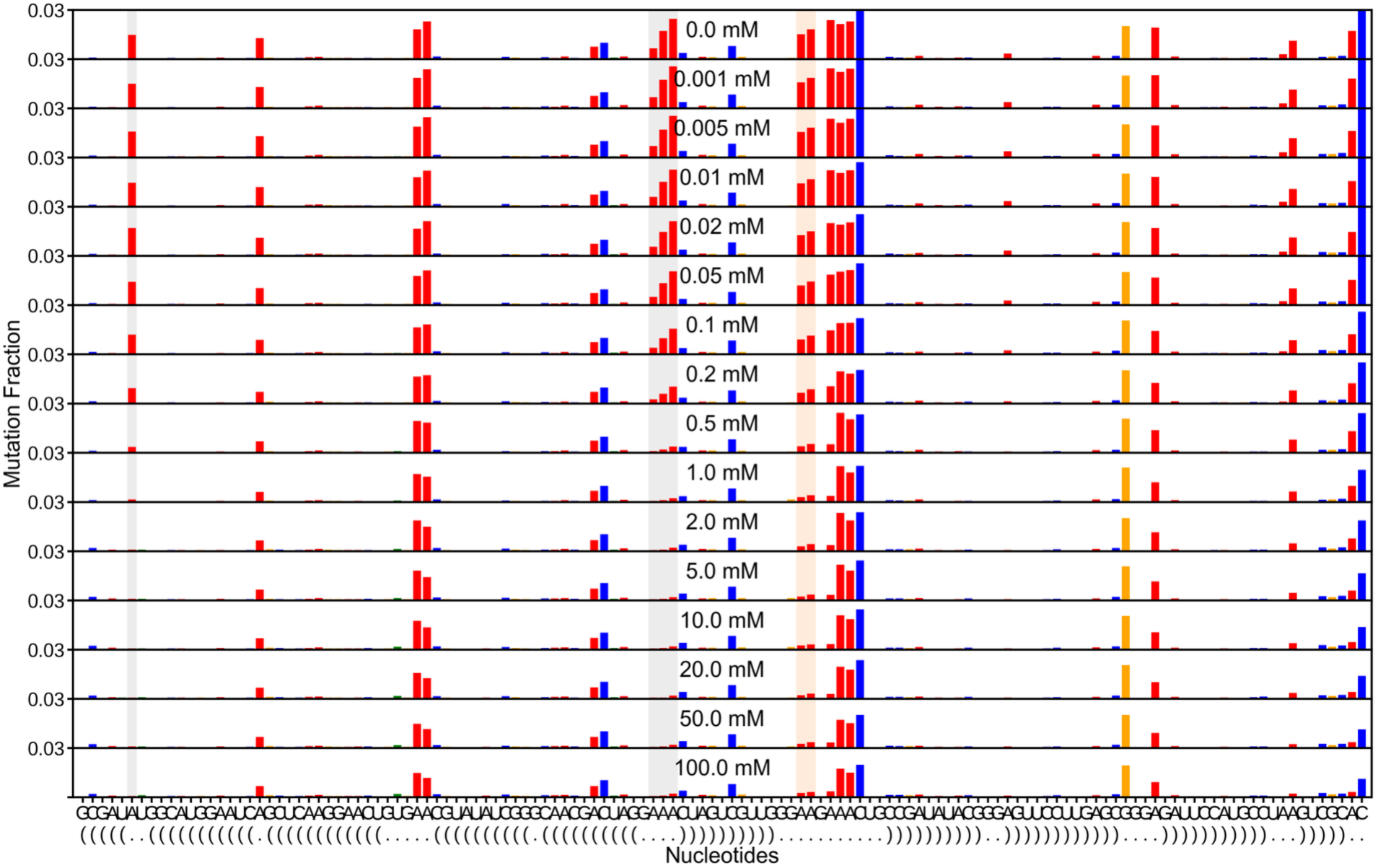
Wild type Mg^2+^ titration without AMP, replicate 1. Stacked population-average per-nucleotide reactivity at each Mg²⁺ concentration (0 to 100 mM) for Wild type without AMP; the y-axis is fixed to the construct’s 99.5th-percentile reactivity. The TL/TLR reporters drop as Mg²⁺ drives docking. Replicate 1 of the Mg-titration whitelist.

**Figure S35:**
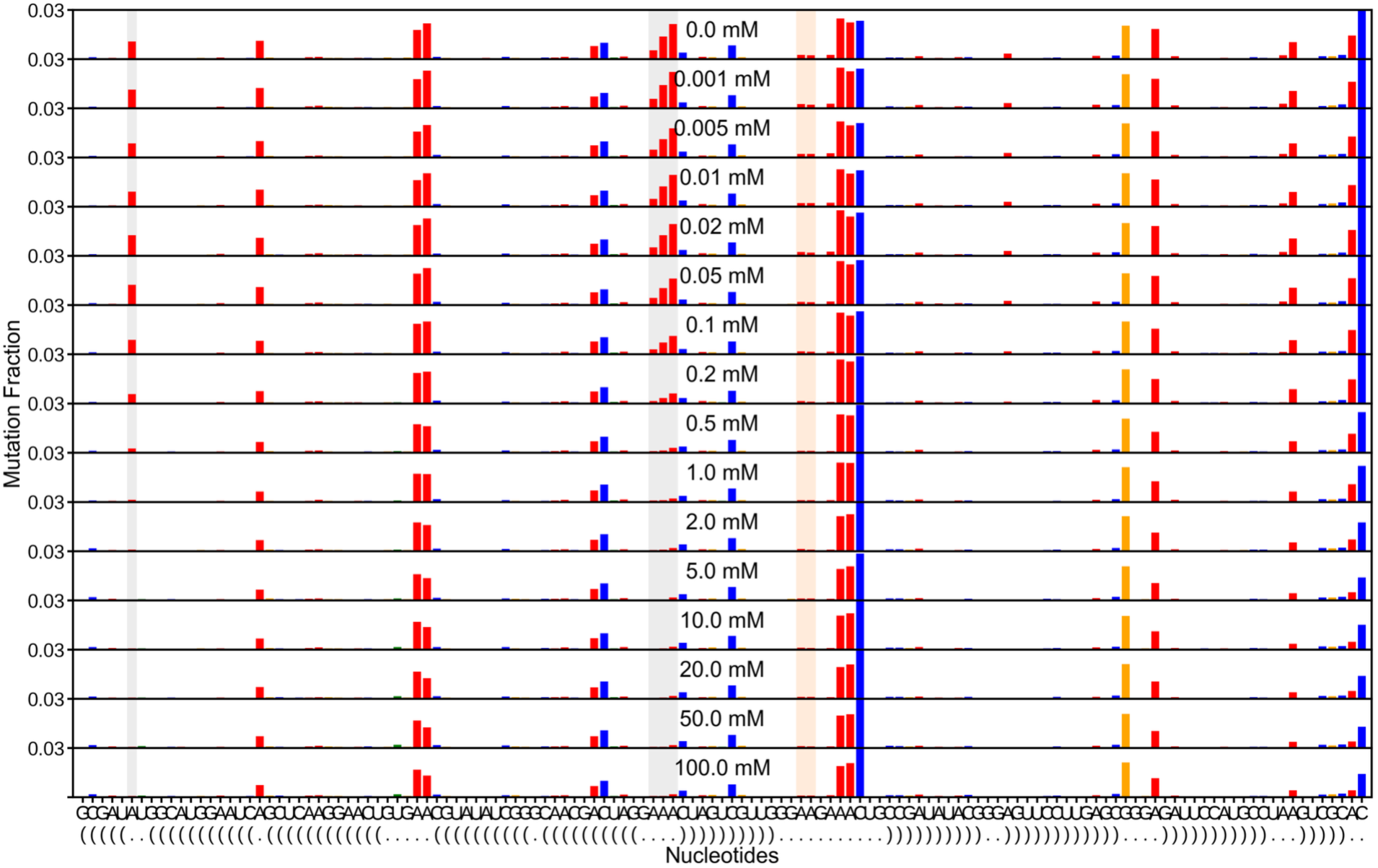
Wild type Mg^2+^ titration with saturating AMP, replicate 1. Stacked population-average per-nucleotide reactivity at each Mg²⁺ concentration (0 to 100 mM) for Wild type with saturating AMP; the y-axis is fixed to the construct’s 99.5th-percentile reactivity. The TL/TLR reporters drop as Mg²⁺ drives docking. Replicate 1 of the Mg-titration whitelist.

**Figure S36:**
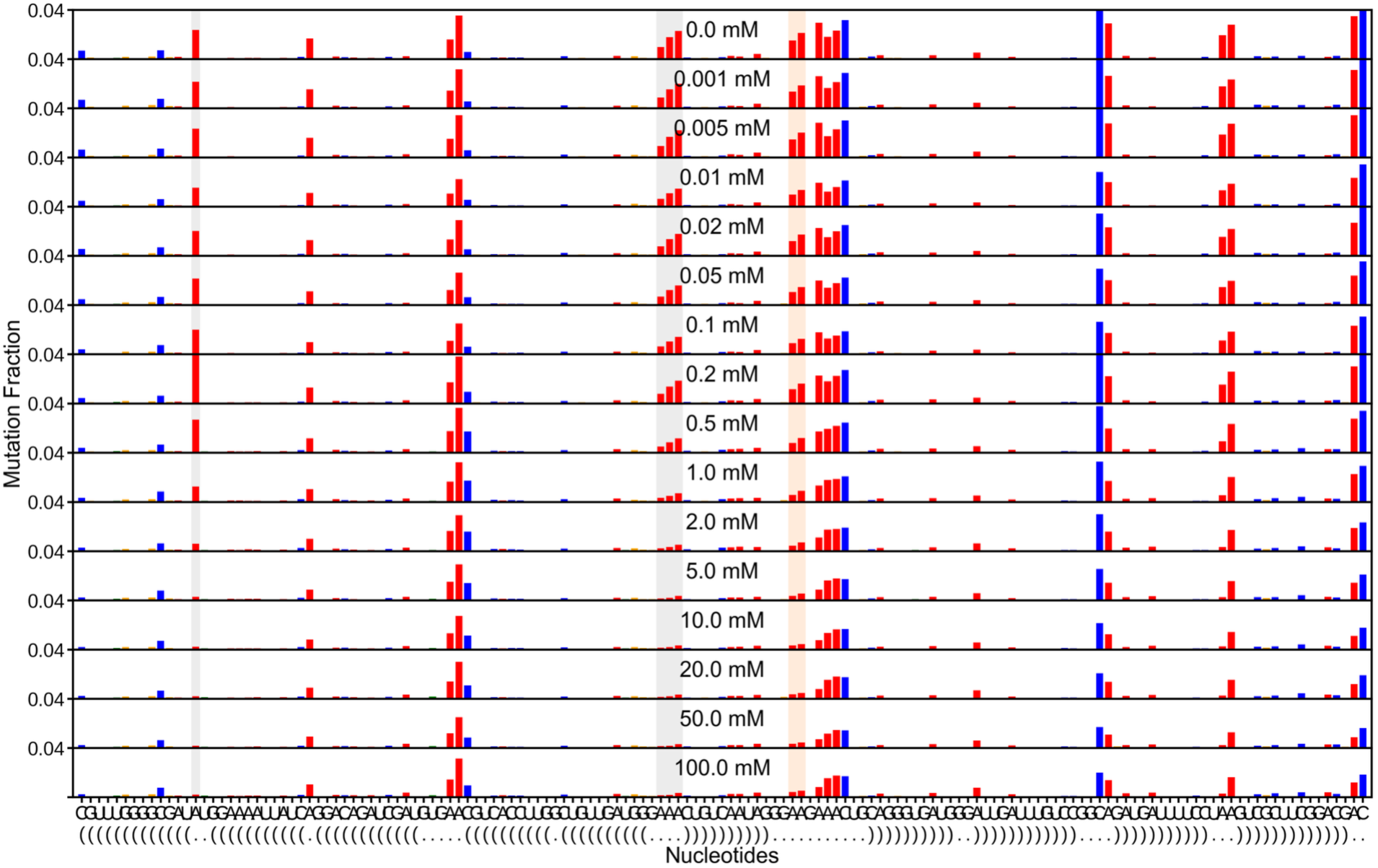
Always-On Mg^2+^ titration without AMP, replicate 1. Stacked population-average per-nucleotide reactivity at each Mg²⁺ concentration (0 to 100 mM) for Always-On without AMP; the y-axis is fixed to the construct’s 99.5th-percentile reactivity. The TL/TLR reporters drop as Mg²⁺ drives docking. Replicate 1 of the Mg-titration whitelist.

**Figure S37:**
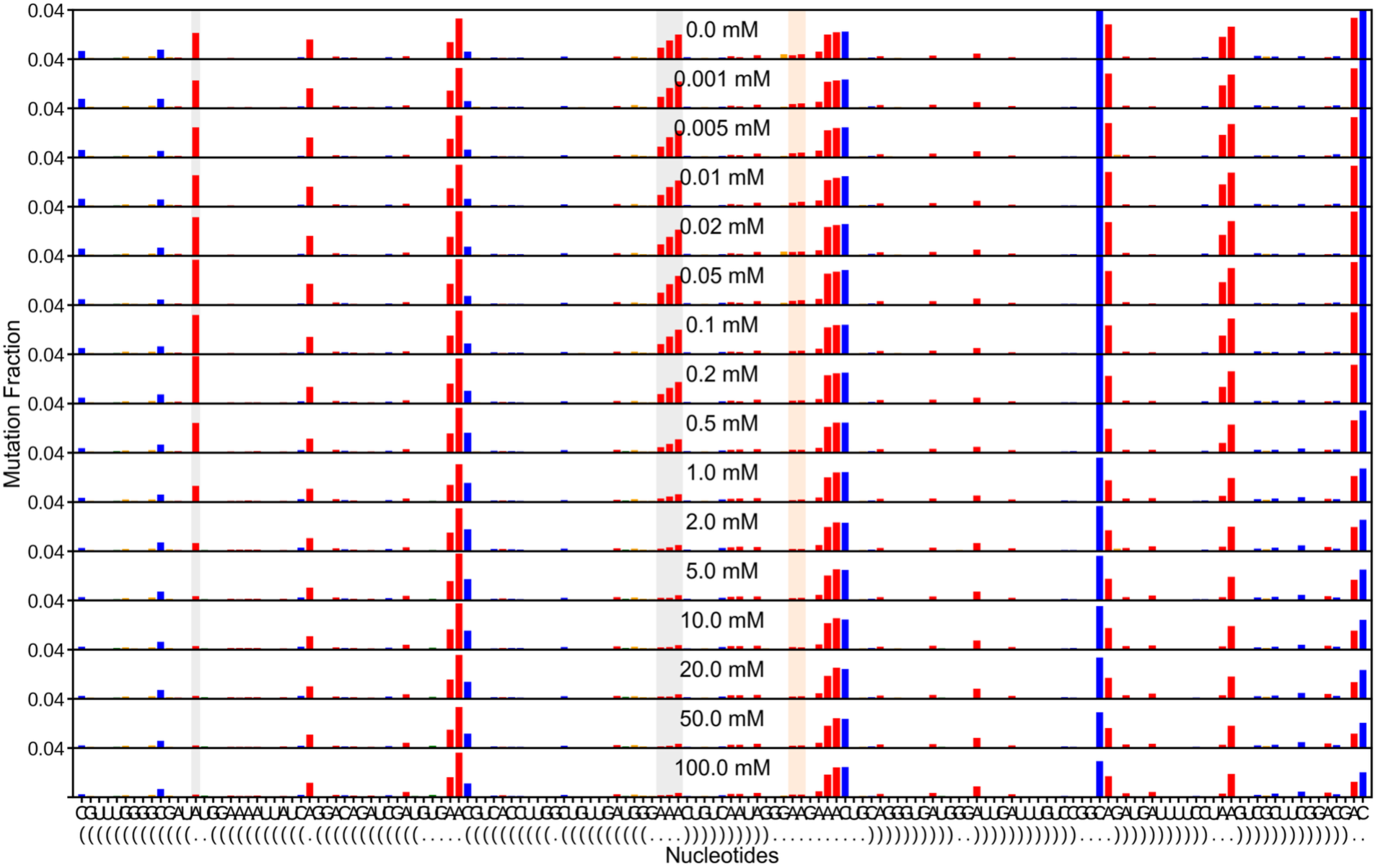
Always-On Mg^2+^ titration with saturating AMP, replicate 1. Stacked population-average per-nucleotide reactivity at each Mg²⁺ concentration (0 to 100 mM) for Always-On with saturating AMP; the y-axis is fixed to the construct’s 99.5th-percentile reactivity. The TL/TLR reporters drop as Mg²⁺ drives docking. Replicate 1 of the Mg-titration whitelist.

**Figure S38:**
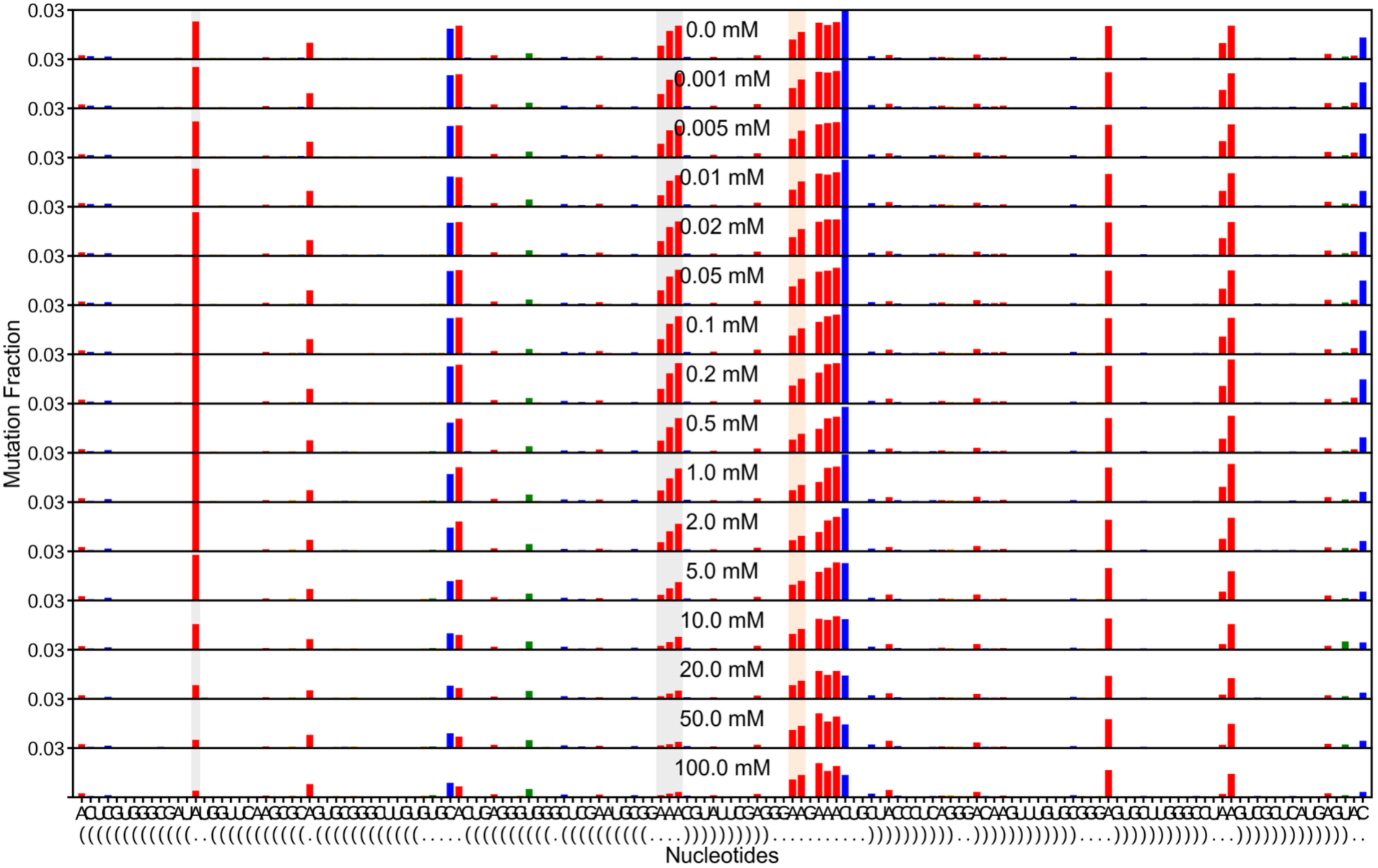
Always-Off Mg^2+^ titration without AMP, replicate 1. Stacked population-average per-nucleotide reactivity at each Mg²⁺ concentration (0 to 100 mM) for Always-Off without AMP; the y-axis is fixed to the construct’s 99.5th-percentile reactivity. The TL/TLR reporters drop as Mg²⁺ drives docking. Replicate 1 of the Mg-titration whitelist.

**Figure S39:**
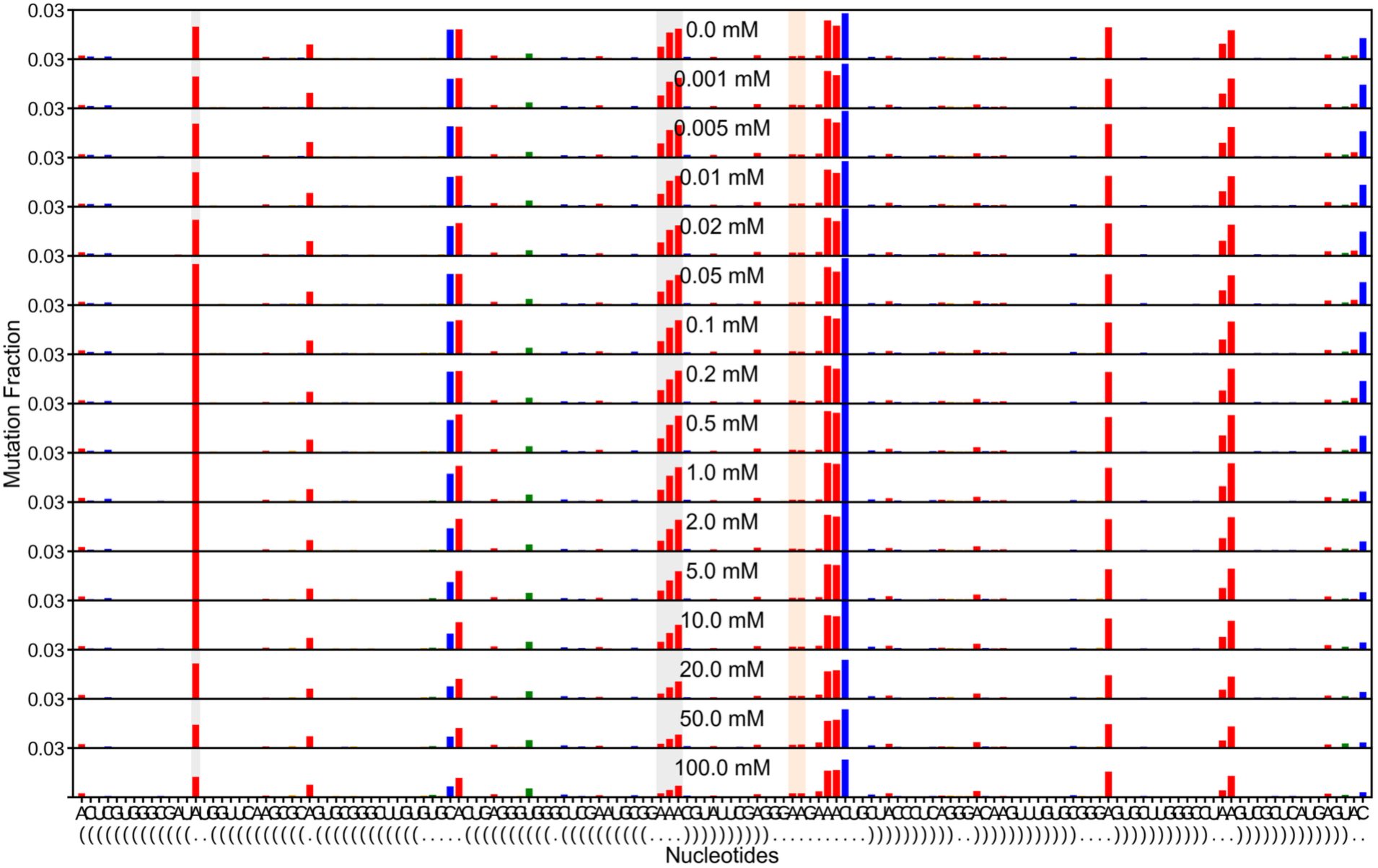
Always-Off Mg^2+^ titration with saturating AMP, replicate 1. Stacked population-average per-nucleotide reactivity at each Mg²⁺ concentration (0 to 100 mM) for Always-Off with saturating AMP; the y-axis is fixed to the construct’s 99.5th-percentile reactivity. The TL/TLR reporters drop as Mg²⁺ drives docking. Replicate 1 of the Mg-titration whitelist.

**Figure S40:**
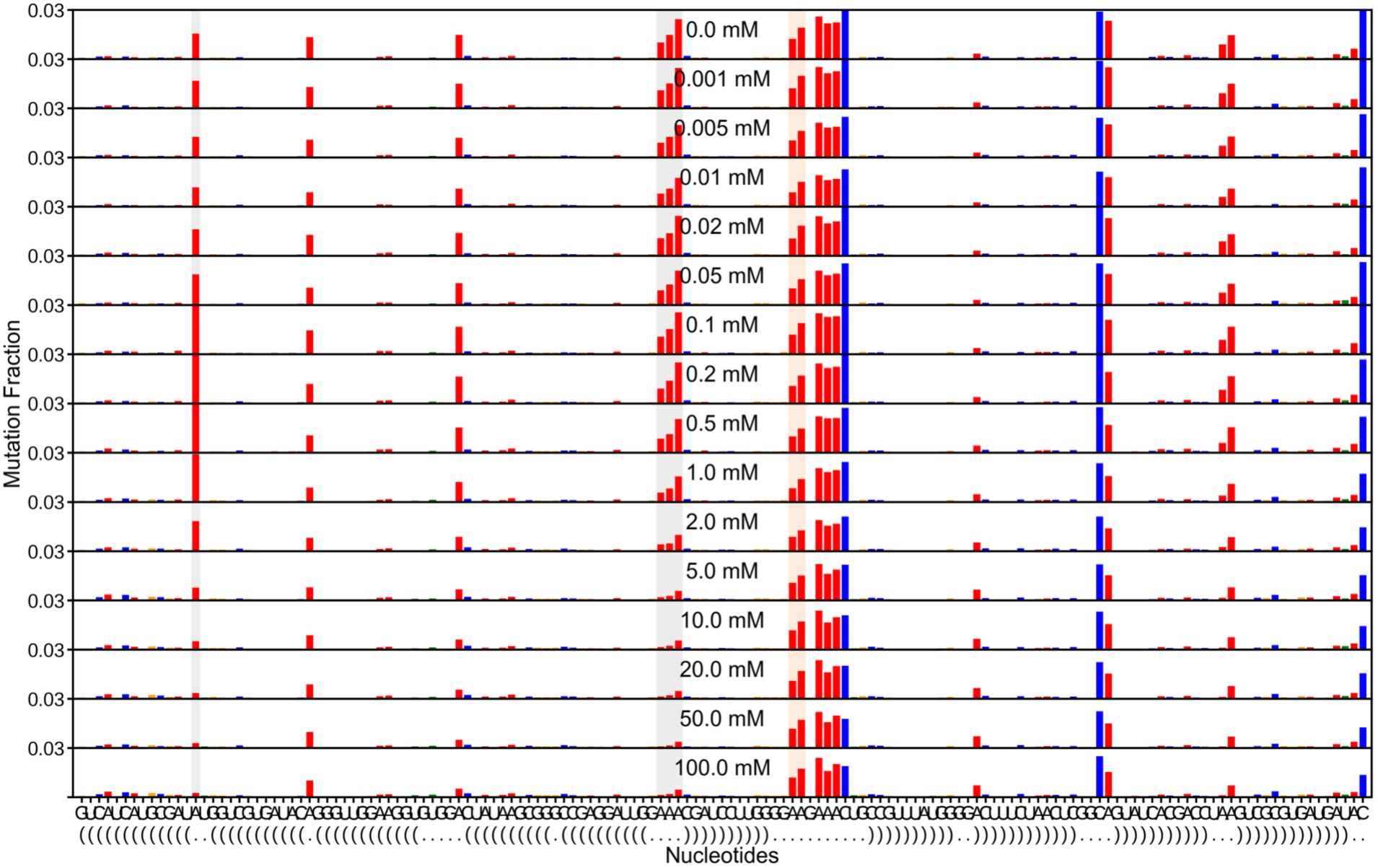
Switch-Off Mg^2+^ titration without AMP, replicate 1. Stacked population-average per-nucleotide reactivity at each Mg²⁺ concentration (0 to 100 mM) for Switch-Off without AMP; the y-axis is fixed to the construct’s 99.5th-percentile reactivity. The TL/TLR reporters drop as Mg²⁺ drives docking. Replicate 1 of the Mg-titration whitelist.

**Figure S41:**
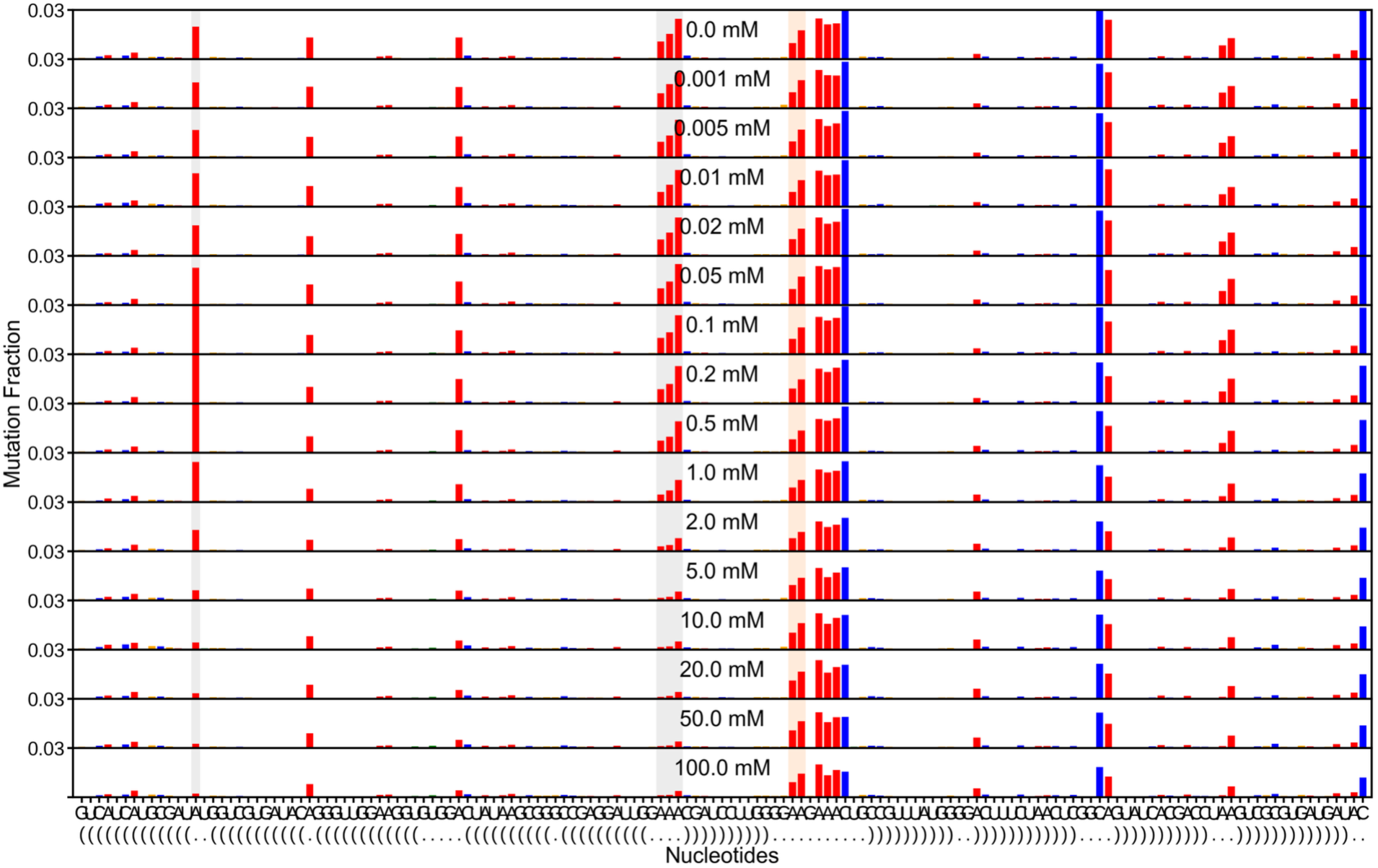
Switch-Off Mg^2+^ titration without AMP, replicate 2. Stacked population-average per-nucleotide reactivity at each Mg²⁺ concentration (0 to 100 mM) for Switch-Off without AMP; the y-axis is fixed to the construct’s 99.5th-percentile reactivity. The TL/TLR reporters drop as Mg²⁺ drives docking. Replicate 2 of the Mg-titration whitelist.

**Figure S42:**
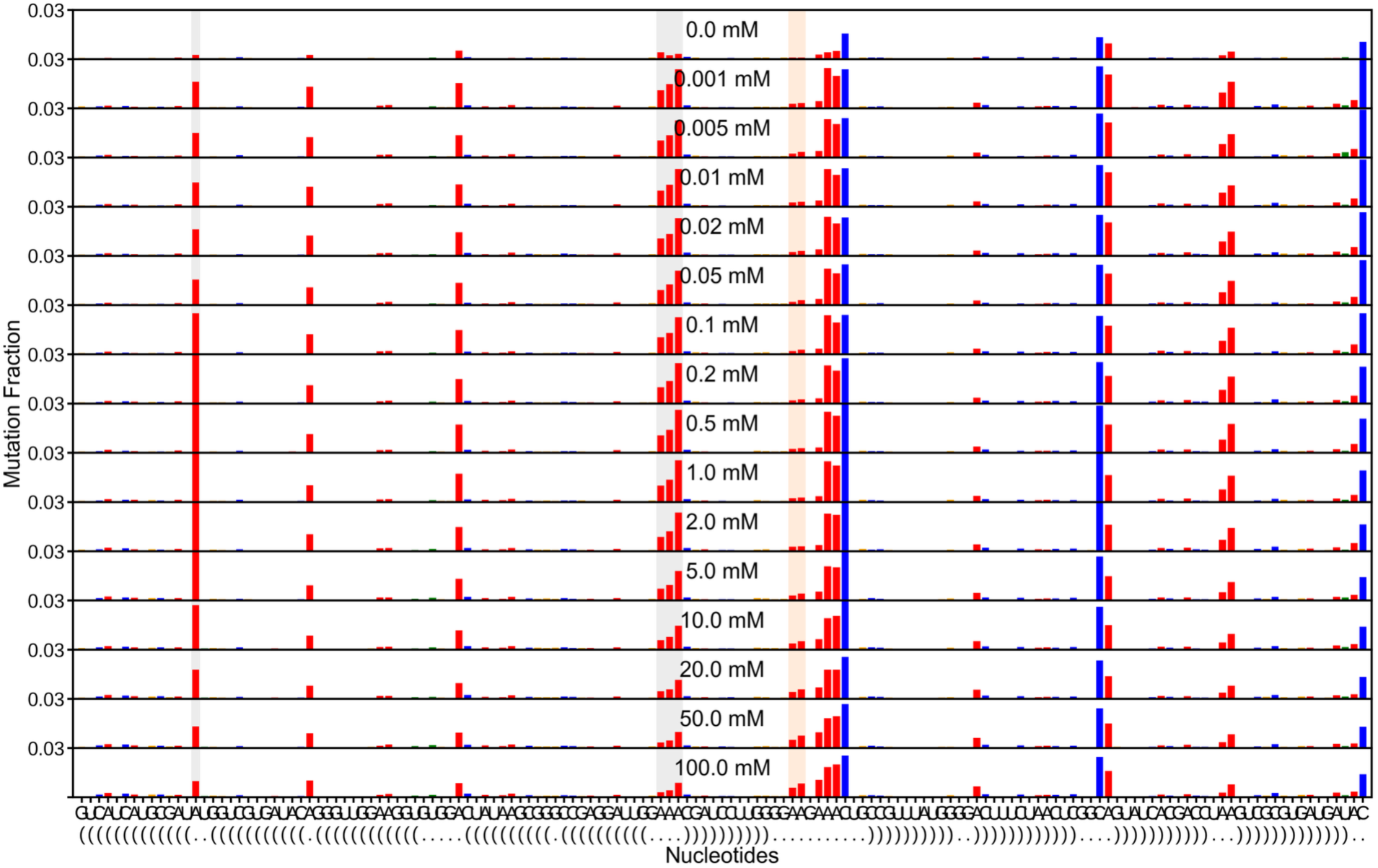
Switch-Off Mg^2+^ titration with saturating AMP, replicate 1. Stacked population-average per-nucleotide reactivity at each Mg²⁺ concentration (0 to 100 mM) for Switch-Off with saturating AMP; the y-axis is fixed to the construct’s 99.5th-percentile reactivity. The TL/TLR reporters drop as Mg²⁺ drives docking. Replicate 1 of the Mg-titration whitelist.

**Figure S43:**
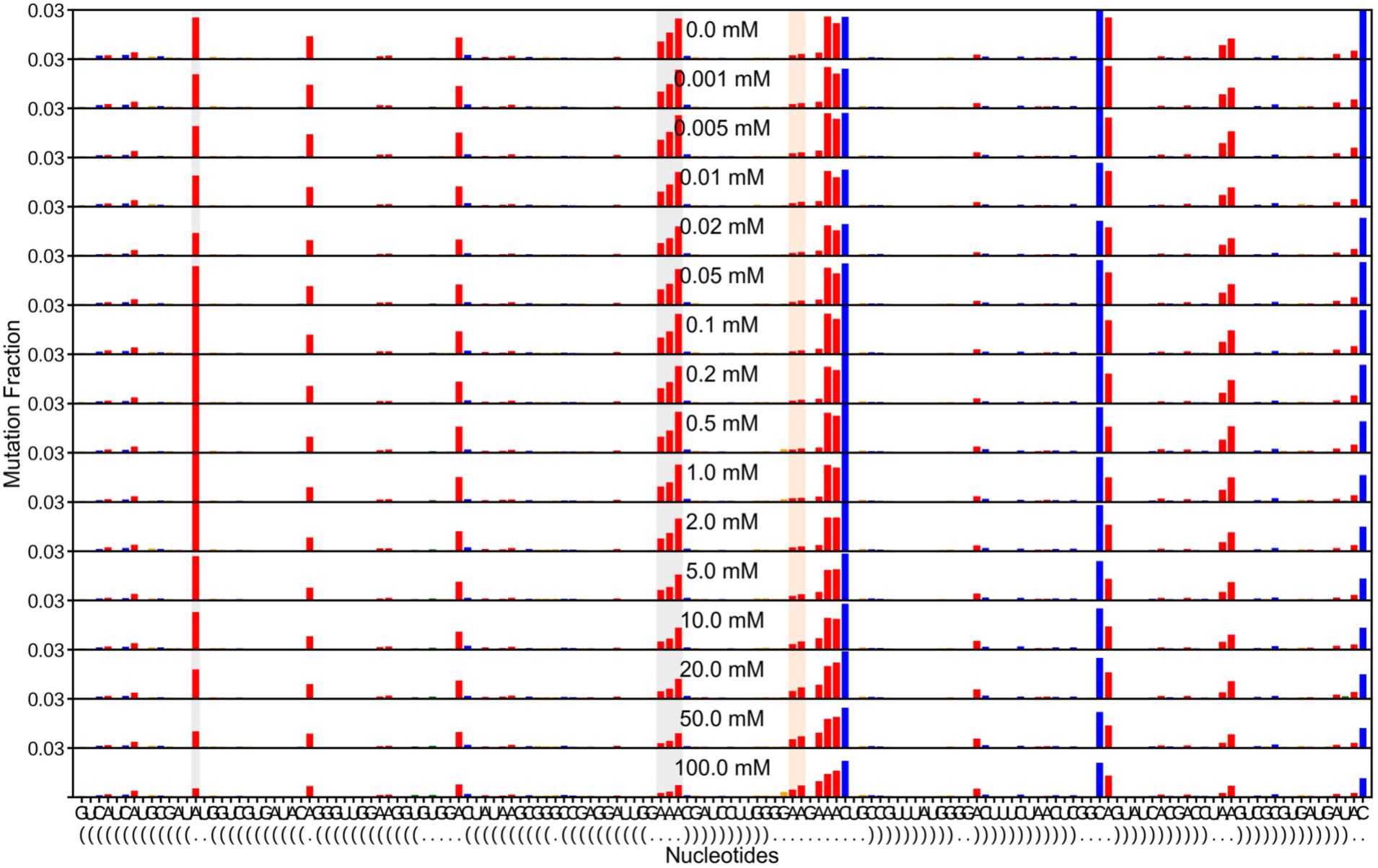
Switch-Off Mg^2+^ titration with saturating AMP, replicate 2. Stacked population-average per-nucleotide reactivity at each Mg²⁺ concentration (0 to 100 mM) for Switch-Off with saturating AMP; the y-axis is fixed to the construct’s 99.5th-percentile reactivity. The TL/TLR reporters drop as Mg²⁺ drives docking. Replicate 2 of the Mg-titration whitelist.

**Figure S44:**
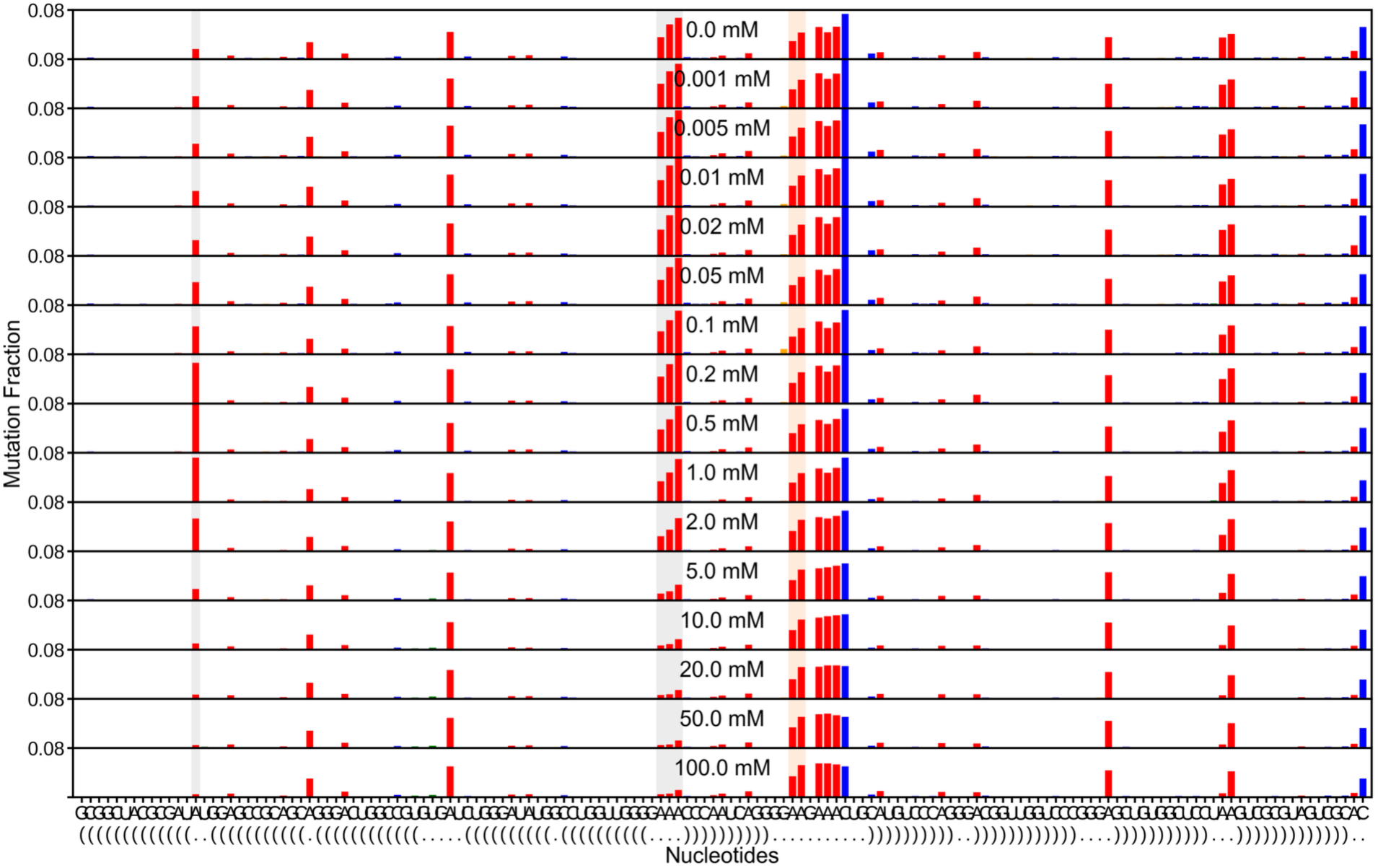
Switch-On Mg^2+^ titration without AMP, replicate 1. Stacked population-average per-nucleotide reactivity at each Mg²⁺ concentration (0 to 100 mM) for Switch-On without AMP; the y-axis is fixed to the construct’s 99.5th-percentile reactivity. The TL/TLR reporters drop as Mg²⁺ drives docking. Replicate 1 of the Mg-titration whitelist.

**Figure S45:**
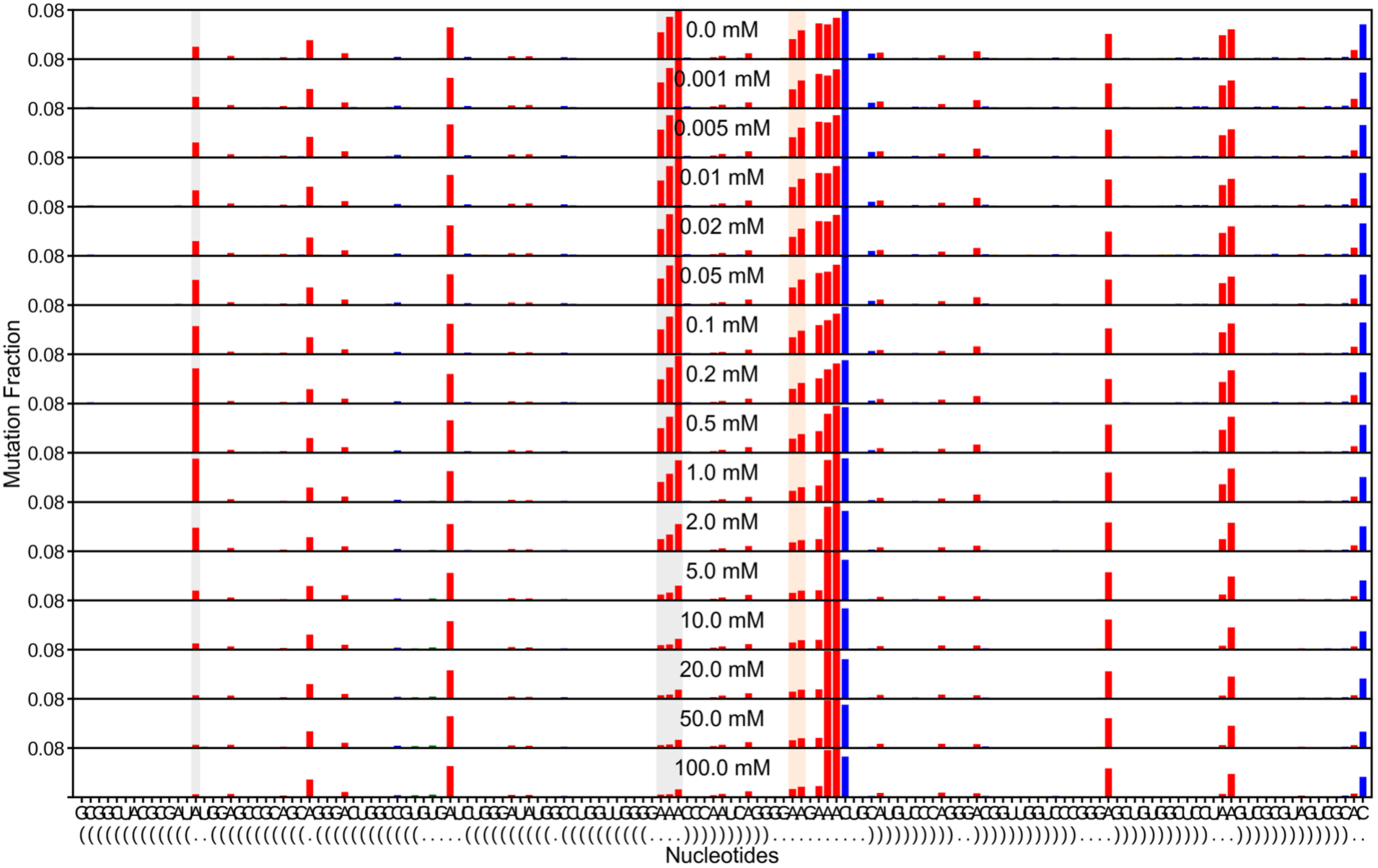
Switch-On Mg^2+^ titration with saturating AMP, replicate 1. Stacked population-average per-nucleotide reactivity at each Mg²⁺ concentration (0 to 100 mM) for Switch-On with saturating AMP; the y-axis is fixed to the construct’s 99.5th-percentile reactivity. The TL/TLR reporters drop as Mg²⁺ drives docking. Replicate 1 of the Mg-titration whitelist.

## Supplemental Tables

**Table S1: Construct sequences**

Sequences of the wild-type nanostructure and every library and representative construct, with their targeted mutations. Provided as the external spreadsheet Sequences.xlsx.

**Table S2: Primer sequences**

All DNA primers used for oligo-pool amplification, primer-assembly template construction, and reverse-transcription barcoding. Provided as the external spreadsheet Primers.xlsx.

## References

1. Nissen P, Hansen J, Ban N, Moore PB, Steitz TA. The Structural Basis of Ribosome Activity in Peptide Bond Synthesis. Science. 2000;289(5481):920–30.

2. Guerrier-Takada C, Gardiner K, Marsh T, Pace N, Altman S. The RNA moiety of ribonuclease P is the catalytic subunit of the enzyme. Cell. 1983;35(3):849–57.

3. Greider CW, Blackburn EH. Identification of a specific telomere terminal transferase activity in tetrahymena extracts. Cell. 1985;43(2):405–13.

4. Cech Thomas R, Steitz Joan A. The Noncoding RNA Revolution—Trashing Old Rules to Forge New Ones. Cell. 2014;157(1):77–94.

5. Brion P, Westhof E. HIERARCHY AND DYNAMICS OF RNA FOLDING. Annual Review of Biophysics and Biomolecular Structure. 1997;26(1):113–37.

6. Tinoco I, Bustamante C. How RNA folds. Journal of Molecular Biology. 1999;293(2):271–81.

7. Batey RT, Rambo RP, Doudna JA. Tertiary Motifs in RNA Structure and Folding. Angewandte Chemie International Edition. 1999;38(16):2326–43.

8. Butcher SE, Pyle AM. The Molecular Interactions That Stabilize RNA Tertiary Structure: RNA Motifs, Patterns, and Networks. Accounts of Chemical Research. 2011;44(12):1302–11.

9. Mandal M, Breaker RR. Adenine riboswitches and gene activation by disruption of a transcription terminator. Nature Structural & Molecular Biology. 2004;11(1):29–35.

10. Breaker RR. Riboswitches and the RNA World. Cold Spring Harbor Perspectives in Biology. 2012;4(2):a003566–a.

11. Serganov A, Nudler E. A Decade of Riboswitches. Cell. 2013;152(1-2):17–24.

12. Garst AD, Edwards AL, Batey RT. Riboswitches: Structures and Mechanisms. Cold Spring Harbor Perspectives in Biology. 2011;3(6):a003533–a.

13. Wickiser JK, Winkler WC, Breaker RR, Crothers DM. The Speed of RNA Transcription and Metabolite Binding Kinetics Operate an FMN Riboswitch. Molecular Cell. 2005;18(1):49–60.

14. Tang J, Breaker RR. Rational design of allosteric ribozymes. Chemistry & Biology. 1997;4(6):453–9.

15. Soukup GA, Breaker RR. Relationship between internucleotide linkage geometry and the stability of RNA. RNA. 1999;5(10):1308–25.

16. Chen X, Ellington AD. Design Principles for Ligand-Sensing, Conformation-Switching Ribozymes. PLoS Computational Biology. 2009;5(12):e1000620.

17. Lemay J-F, Penedo JC, Tremblay R, Lilley DMJ, Lafontaine Daniel A. Folding of the Adenine Riboswitch. Chemistry & Biology. 2006;13(8):857–68.

18. Frieda KL, Block SM. Direct Observation of Cotranscriptional Folding in an Adenine Riboswitch. Science. 2012;338(6105):397–400.

19. Treiber DK, Williamson JR. Exposing the kinetic traps in RNA folding. Current Opinion in Structural Biology. 1999;9(3):339–45.

20. Russell R, Zhuang X, Babcock HP, Millett IS, Doniach S, Chu S, et al. Exploring the folding landscape of a structured RNA. Proceedings of the National Academy of Sciences. 2002;99(1):155–60.

21. Greenleaf WJ, Frieda KL, Foster DAN, Woodside MT, Block SM. Direct Observation of Hierarchical Folding in Single Riboswitch Aptamers. Science. 2008;319(5863):630–3.

22. Wittmann A, Suess B. Engineered riboswitches: Expanding researchers’ toolbox with synthetic RNA regulators. FEBS Letters. 2012;586(15):2076–83.

23. Wachsmuth M, Findeiss S, Weissheimer N, Stadler PF, Morl M. De novo design of a synthetic riboswitch that regulates transcription termination. Nucleic Acids Research. 2013;41(4):2541–51.

24. Berens C, Suess B. Riboswitch engineering — making the all-important second and third steps. Current Opinion in Biotechnology. 2015;31:10–5.

25. Win MN, Smolke CD. A modular and extensible RNA-based gene-regulatory platform for engineering cellular function. Proceedings of the National Academy of Sciences. 2007;104(36):14283–8.

26. Win MN, Smolke CD. Higher-order cellular information processing with synthetic RNA devices. Science. 2008;322(5900):456–60.

27. Ceres P, Garst AD, Marcano-Velázquez JG, Batey RT. Modularity of select riboswitch expression platforms enables facile engineering of novel genetic regulatory devices. ACS Synthetic Biology. 2013;2(8):463–72.

28. Dixon N, Duncan JN, Geerlings T, Dunstan MS, McCarthy JEG, Leys D, et al. Reengineering orthogonally selective riboswitches. Proceedings of the National Academy of Sciences. 2010;107(7):2830–5.

29. Lynch SA, Desai SK, Sajja HK, Gallivan JP. A high-throughput screen for synthetic riboswitches reveals mechanistic insights into their function. Chemistry & Biology. 2007;14(2):173–84.

30. Wieland M, Hartig JS. Improved aptazyme design and in vivo screening enable riboswitching in bacteria. Angewandte Chemie International Edition. 2008;47(14):2604–7.

31. Frank J, Agrawal RK. A ratchet-like inter-subunit reorganization of the ribosome during translocation. Nature. 2000;406(6793):318–22.

32. Mondragón A. Structural Studies of RNase P. Annual Review of Biophysics. 2013;42(1):537–57.

33. Vicens Q, Cech TR. Atomic level architecture of group I introns revealed. Trends in Biochemical Sciences. 2006;31(1):41–51.

34. Marcia M, Pyle Anna M. Visualizing Group II Intron Catalysis through the Stages of Splicing. Cell. 2012;151(3):497–507.

35. Sassanfar M, Szostak JW. An RNA motif that binds ATP. Nature. 1993;364(6437):550–3.

36. Dieckmann T, Suzuki E, Nakamura GK, Feigon J. Solution structure of an ATP-binding RNA aptamer reveals a novel fold. RNA. 1996;2(7):628–40.

37. Costa M, Michel F. Frequent use of the same tertiary motif by self-folding RNAs. The EMBO Journal. 1995;14(6):1276–85.

38. Geary C, Baudrey S, Jaeger L. Comprehensive features of natural and in vitro selected GNRA tetraloop-binding receptors. Nucleic Acids Research. 2008;36(4):1138–52.

39. Yesselman JD, Eiler D, Carlson ED, Gotrik MR, d’Aquino AE, Ooms AN, et al. Computational design of three-dimensional RNA structure and function. Nature Nanotechnology. 2019;14(9):866–73.

40. Lorenz R, Bernhart SH, Höner zu Siederdissen C, Tafer H, Flamm C, Stadler PF, et al. ViennaRNA Package 2.0. Algorithms for Molecular Biology. 2011;6:26.

41. Jurich CP, Brivanlou A, Rouskin S, Yesselman JD. Web-based platform for analysis of RNA folding from high throughput chemical probing data. Nucleic Acids Res. 2022;50(W1):W266–W71.

42. Zubradt M, Gupta P, Persad S, Lambowitz AM, Weissman JS, Rouskin S. DMS-MaPseq for genome-wide or targeted RNA structure probing in vivo. Nature Methods. 2017;14(1):75–82.

43. Yesselman JD, Denny SK, Bisaria N, Herschlag D, Greenleaf WJ, Das R. Sequence-dependent RNA helix conformational preferences predictably impact tertiary structure formation. Proceedings of the National Academy of Sciences. 2019;116(34):16847–55.

44. Wyman J, Gill SJ. Binding and Linkage: Functional Chemistry of Biological Macromolecules. Mill Valley, CA: University Science Books; 1990.

45. Reinhart GD. Quantitative Analysis and Interpretation of Allosteric Behavior. Methods in Enzymology 2004. p. 187–203.

46. Draper DE, Grilley D, Soto AM. Ions and RNA Folding. Annual Review of Biophysics and Biomolecular Structure. 2005;34(1):221–43.

47. Draper DE. RNA Folding: Thermodynamic and Molecular Descriptions of the Roles of Ions. Biophysical Journal. 2008;95(12):5489–95.

48. Bisaria N, Greenfeld M, Limouse C, Pavlichin DS, Mabuchi H, Herschlag D. Kinetic and thermodynamic framework for P4-P6 RNA reveals tertiary motif modularity and modulation of the folding preferred pathway. Proceedings of the National Academy of Sciences. 2016;113(34).

49. Denny SK, Bisaria N, Yesselman JD, Das R, Herschlag D, Greenleaf WJ. High-Throughput Investigation of Diverse Junction Elements in RNA Tertiary Folding. Cell. 2018;174(2):377–90 e20.

50. Klein DJ, Schmeing TM, Moore PB, Steitz TA. The kink-turn: a new RNA secondary structure motif. The EMBO Journal. 2001;20(15):4214–21.

51. Goody TA, Melcher SE, Norman DG, Lilley DMJ. The kink-turn motif in RNA is dimorphic, and metal ion-dependent. RNA. 2004;10(2):254–64.

52. Lilley DMJ. The K-turn motif in riboswitches and other RNA species. Biochimica et Biophysica Acta (BBA) - Gene Regulatory Mechanisms. 2014;1839(10):995–1004.

53. Kappel K, Zhang K, Su Z, Watkins AM, Kladwang W, Li S, et al. Accelerated cryo-EM-guided determination of three-dimensional RNA-only structures. Nat Methods. 2020;17(7):699–707.

54. McPhee SA, Huang L, Lilley DMJ. A critical base pair in k-turns that confers folding characteristics and correlates with biological function. Nature Communications. 2014;5(1).

55. Daldrop P, Lilley DMJ. The plasticity of a structural motif in RNA: Structural polymorphism of a kink turn as a function of its environment. RNA. 2013;19(3):357–64.

56. Hilser VJ, Wrabl JO, Motlagh HN. Structural and Energetic Basis of Allostery. Annual Review of Biophysics. 2012;41(1):585–609.

